# A dimensional link between gain–loss economic preferences and psychopathology

**DOI:** 10.64898/2026.09.09.750300

**Authors:** Yang-Yang Feng, Ilya E. Monosov

## Abstract

Human decision-makers differ in how they value and trade off reward, risk, and information when evaluating future gains and losses, and these preferences are central to well-being. However, how multi-attribute economic valuation relates to psychopathology remains unclear. Here, 1,954 individuals completed a transdiagnostic symptom battery and a multi-attribute decision-making task that independently manipulated expected reward, reward uncertainty, and non-instrumental information under matched gain and loss contexts. On average, when choosing among gains, participants preferred larger rewards, avoided risk, and sought advance information. When choosing among losses, they still preferred better outcomes and advance information, but less strongly. Their risk preference, by contrast, reversed, from risk-averse to risk-seeking. Substantial individual variability accompanied these population-level context effects, and different patterns of variability were associated with separate dimensions of psychopathology. Anxious– depressive and social withdrawal symptoms were associated with increased reward sensitivity across contexts, whereas compulsive–intrusive symptoms were associated with reduced reward sensitivity. Symptom-related differences in risk and information preferences were not fully explained by a uniform scaling of overall economic sensitivity, revealing additional attribute- and context-specific variation. A data-driven analysis summarized these symptom–preference associations in a single latent dimension that contrasted compulsive–intrusive symptoms with anxious–depressive and social withdrawal symptoms. An independently identified subgroup that strongly sought information about gains but avoided it about losses mirrored this profile and was enriched for compulsive–intrusive symptoms. These findings reveal structured, low-dimensional variation linking multi-attribute decision-making to psychopathology and could support a dimensional, task-based approach to psychiatric nosology.

## 1 Introduction

Human decision-makers differ in how they value and trade off reward, risk, and information about the future when making choices about gains and losses [1–5]. These multi-attribute decision processes are thought to vary across individuals and contexts, shaping everyday choices and impacting mental health [1, 6–8]. Psychiatric disorders have been linked to alterations in economic preferences and their modulation by outcome valence—for example, in shifts in preferences between gain and loss contexts. Prior work has largely examined these preferences and context effects in isolation. But everyday choices seldom isolate a single attribute, instead decision options or offers usually differ in reward, risk, and information all at once, and these different attributes of each offer is often weighed against one another. However, how this integration is accomplished, and how it relates to psychopathology, remains unclear. Addressing this gap is a central goal in computational psychiatry [7, 9–14].

Indeed, prior work has examined how reward and risk influence choice and relate to psychiatric symptoms [4, 8, 14–17]. But beyond these extrinsic factors, humans are also intrinsically motivated to seek information to resolve uncertainty, even when it has no instrumental value—a process linked to neural circuits implicated in anxiety, compulsivity, and depression [18–25]. Individual differences in information-seeking have themselves been linked to psychological traits and symptoms, including anxious and obsessive–compulsive traits [25–29]. Yet how individuals integrate reward, uncertainty, and non-instrumental information across gain and loss contexts—and how this integration relates to psychopathology— remains poorly understood. Few studies have examined such multi-attribute decisions requiring trade-offs between reward, uncertainty, and information [21, 23, 30–32] and, to our knowledge, none have combined joint measurement of reward, risk, and information preferences with matched gain and loss contexts and a broad transdiagnostic symptom battery. Moreover, prior work has largely focused on gain-framed or mixed gain–loss gambles [6, 31, 33–35], with comparatively little attention to multi-attribute choices in loss contexts [31, 34]—despite evidence that loss framing plays a central role in psychiatric symptoms.

This gap is important given evidence that adaptive behavior requires modulating strategies depending on context [12, 36–40], and in particular, economic preferences may exhibit systematic changes between gain and loss contexts [6, 31, 33, 34]. For example, risk attitudes can reverse and information-seeking may be altered under loss or other negative contexts in both humans and animals [6, 20, 22, 33–35, 41–43]. Such asymmetries are especially relevant for psychiatry because many psychiatric illnesses are characterized by heightened sensitivity to loss, punishment, and aversive outcomes. It is therefore possible that gain–loss asymmetries could play an important role in psychiatric phenomena such as negativity bias and maladaptive avoidance [8, 15, 35, 44, 45].

Across mood, anxiety, and compulsive disorders, studies have reported changes in reward sensitivity, risk-seeking, and information-seeking, but these findings are often fragmented across tasks that isolate single attributes [15–17, 20, 25, 41, 46–48]. Some of this work has focused on learning-based paradigms, in which preferences are inferred from trial-by-trial updating from gains and losses [8, 15, 35, 42, 44, 49–52]. While such approaches have provided important insights into reinforcement learning processes and human behavior, they may not fully or easily isolate how individuals value reward, uncertainty, and information at the time of choice [7, 9, 36, 37]. Also, computational psychiatry has proposed that core symptom dimensions—such as anhedonia, intolerance of uncertainty, and negativity bias—reflect disturbances in underlying computations of value, uncertainty, and prediction error, which should manifest as systematic alterations in economic preferences [7, 9, 11, 15, 45, 53–55]. However, linking specific patterns of reward, risk, and information preferences to specific symptom dimensions has proven difficult, in part because studies often rely on heterogeneous tasks and small, diagnostically defined samples that obscure underlying dimensional structure (see discussion in [16, 56–58]).

Here, we employ a large-sample, multi-attribute decision-making task to jointly measure preferences for expected reward, reward uncertainty, and non-instrumental information under matched gain and loss contexts, alongside three transdiagnostic psychopathology dimensions: anxious–depressive (AD), compulsive–intrusive (CI), and social withdrawal (SW) [16]. This design allows us to (i) characterize how reward, risk, and information preferences and their scaling with uncertainty differ between gain and loss contexts at both population and individual levels; (ii) test how these preferences and their context-dependent changes relate to psychopathology, evaluating competing computational accounts based on motivation, uncertainty processing, and negativity bias; and (iii) search for computationally defined subgroups, including individuals who seek information about gains but avoid information about losses, providing a dimensional, task-based framework for understanding economic decision-making in mental health and disease.

## 2 Results

### 2.1 Measuring economic preferences and psychopathology

To capture joint variability in economic preferences and psychopathology, 1,974 adults recruited online performed a multiattribute choice task (Figure 1A) and completed a battery of symptom questionnaires (Figure 1B) previously shown to capture a 3-factor model of psychopathology; the 1,954 participants whose choice models converged in both contexts (Methods) contribute to all analyses below. This battery and approach have been validated in many recent studies [16, 17, 59–71]. The multi-attribute decision-making task used in this study measured economic preferences in gain and loss contexts using offers that varied independently in reward, risk, and informativeness (Figure 1). A gain-only version of this task was previously developed to assess attitudes toward risk and non-instrumental information in non-human primates and human participants [23] and used to study how the habenula-raphe circuit encodes subjective value across multiple attributes [23, 32].

**Figure 1.**
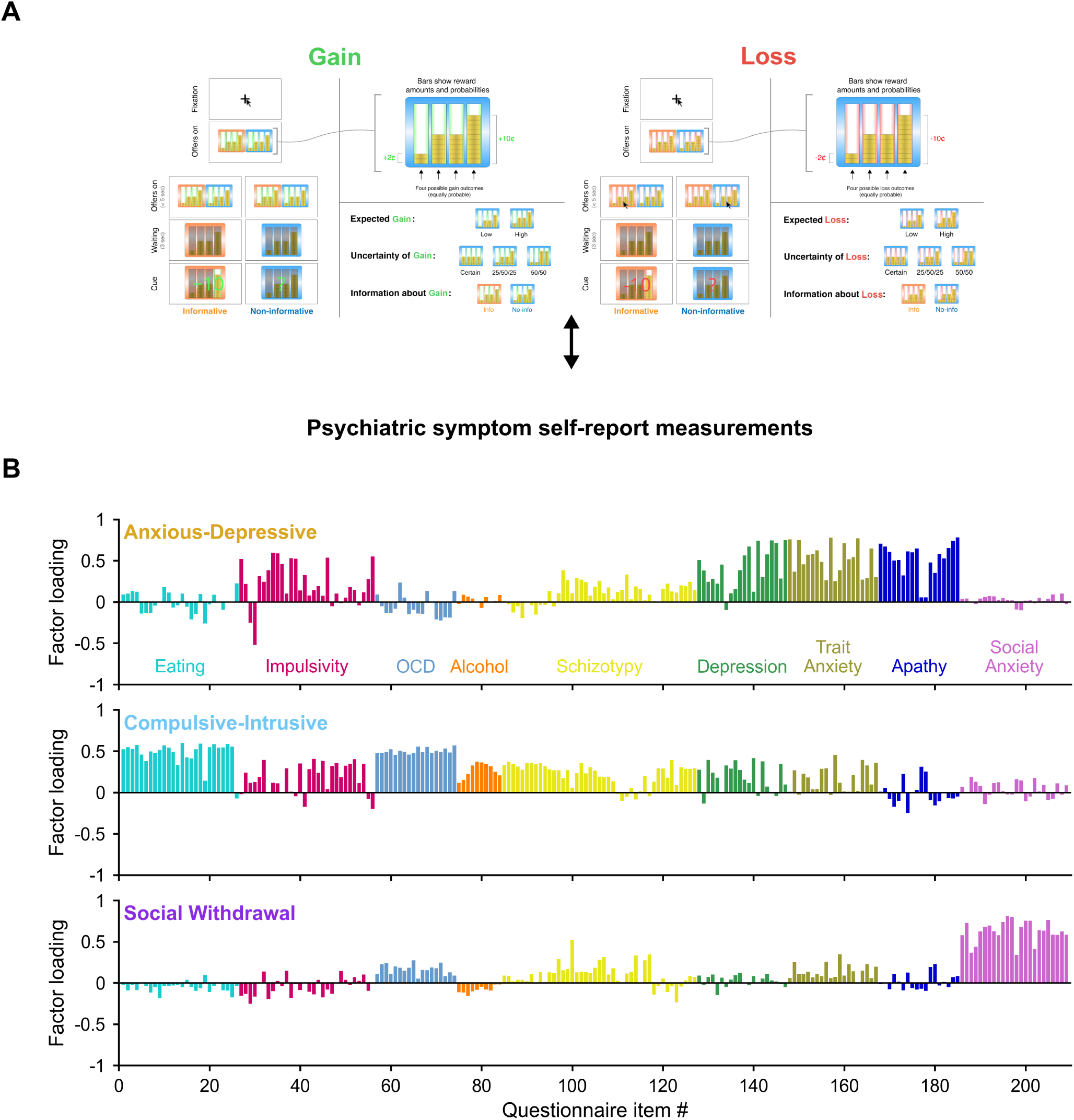
Measuring individual-level economic preferences and psychopathology in a large population (n=1954). We measured individual economic preferences—including the novel attribute of non-instrumental information-seeking— using a multi-attribute decision task performed in both gain and loss contexts, and characterized psychopathology using a 3-factor model (AD, CI, SW) derived from a 209-item battery of symptom questionnaires, providing the joint economic– psychiatric measurements needed to ask how information preferences and their context-dependence relate to dissociable dimensions of psychopathology. **(A)** Gain and loss multi-attribute decision task, drawn for the Gain (left, green) and Loss (right, red) contexts. The trial schematic in each context gives the sequence of screens: fixation, offers on (*<*5 s), a 3 s waiting period, and then the cue, which was either *Informative* or *Non-informative*. The enlarged offer at the top of each context shows how an offer was displayed: four bars, each a stack of coins worth $0.01, whose heights are the four possible monetary outcomes of that offer (labeled on the figure as “Bars show reward amounts and probabilities” and “Four possible gain (loss) outcomes (equally probable)”); outcomes ranged from $0.01 to $0.11 per trial in gain and from — $0.01 to —$0.11 in loss. On each trial participants chose between two such offers with a mouse click, and the chosen offer delivered one of its four outcomes uniformly at random. Offers therefore differed in expected reward (E[Rew]; the average bar height) and in reward uncertainty (Unc[Rew]; the spread across bars), and each offer’s frame color additionally marked it as an Info or a Noinfo offer: an Info offer delivered a cue that revealed which of its four outcomes would be obtained, while a Noinfo offer delivered an uninformative cue. This cue was non-instrumental—it had no influence on the outcome and could not be used to alter behavior—and participants were explicitly instructed of this. The offer-design panel at the right of each context shows the three attributes varied independently: expected gain/loss (Low vs. High), uncertainty of gain/loss (three levels, drawn as Certain, 25/50/25 and 50/50 outcome distributions, whose outcome standard deviations are the 0, 2.83 and 4.00 levels used in Figure 4A), and information about gain/loss (Info vs. No-info). Each participant completed 150 trials in each context in separate blocks, with task structure held constant and only the framing differing: in the gain context, participants were instructed that they could earn reward on each trial that accumulated toward their total payment; in the loss context, participants were instructed that they would start with $18 and gradually lose reward across trials. The starting amount was calibrated so that average earnings were equal between contexts (mean total earnings $9.26 ± $0.31 SD in gain, $9.25 ± $0.32 SD in loss). This design isolates preferences for each attribute independently, including non-instrumental information, whose value cannot be derived from any objective utility calculation. **(B)** 3-factor model of psychopathology, from the psychiatric symptom selfreport measurements (n=1954). Each row of bars is one factor—anxious-depressive (AD), compulsive-intrusive (CI) and social withdrawal (SW)—and each bar is the loading of one of the 209 individual questionnaire items (z-scored across participants) recovered by maximum-likelihood factor analysis with the number of factors fixed a priori at three (following Gillan et al. 2016; Methods). Items are grouped and colored along the x-axis by their source instrument, in the order eating disorders (EAT-26), impulsivity (BIS-11), obsessive-compulsive disorder (OCI-R), alcohol use (AUDIT), schizotypy (Short Scales for Schizotypy), depression (Zung SDS), trait anxiety (STAI-T), apathy (AES) and social anxiety (LSAS). The recovered factor structure qualitatively matches the original 3-factor model of Gillan et al. (2016): the AD factor is dominated by symptoms of trait anxiety, depression, apathy and impulsivity; the CI factor by symptoms of OCD, eating disorders and alcohol use; and the SW factor by symptoms of social anxiety. These three factors, each linked by theory to distinct computational processes (motivational dysregulation for AD, uncertainty processing for CI, and social-evaluative concerns for SW), form the psychiatric axes we relate to economic preferences in subsequent analyses. Takeaway: our task and 3-factor model together yield individual-level estimates of multi-attribute economic preferences in gain and loss alongside dimensional measures of three psychopathology axes that span common internalizing and compulsive symptom dimensions.

The task consisted of 150 trials in each context (gain and loss). And it was designed such that each context was performed as similarly as possible, with only different framing. At the start of the gain context, participants were instructed that on each trial they could earn reward based on their choices, accumulating towards their total payment, whereas at the start of the loss context, participants were instructed that they would start off with $18 and on each trial would gradually lose reward. Participants chose between a pair of offers on each trial with a mouse click. In the gain context, when chosen, each offer provided a monetary reward that was randomly drawn from a set of four possible outcomes, depicted as four stacks of coins. Participants were instructed that each coin obtained in this context would equate to real gains in their earnings. In the loss context, trials worked the same way, only participants were instructed that each coin obtained in this context would equate to real losses in their earnings, which would be subtracted from a starting total of $18. This starting amount was calibrated such that on average, participants earned the same total amount from each context. Each coin was worth $0.01. On average, participants earned a total of $9.26 ($0.31 standard deviation, SD) or $9.25 ($0.32 SD) after their choices in the gain or loss context, respectively. Different offers provided different probability distributions of rewards, which could have different levels of reward expectation (E[Rew]; the mean height of the stacks) and uncertainty (Unc[Rew]; the variability of the height of the stacks). Each offer also had a color that indicated whether it was an Info or Noinfo offer. When chosen, Info offers provided an informative cue indicating which of the four possible outcomes would be delivered into the participant’s winnings on that trial, whereas Noinfo offers did not. This information was non-instrumental and hence had no objective value because there was no way to use it to influence the outcome, and participants were clearly instructed on this following prior procedures [23]. Thus, each offer had multiple attributes— expected reward, reward uncertainty, and non-instrumental informativeness—that could elicit economic preferences in gain and loss decisions, enabling us to probe the value assigned to information alongside classical reward and risk.

To assess psychopathology, we measured each participant’s scores along three previously-identified factor dimensions of psychopathology: anxious–depressive (AD), compulsive–intrusive (CI), and social withdrawal (SW) [16]. These factors are derived from a battery of 209 questions across 9 surveys measuring symptoms of eating disorders (EAT-26), alcohol use disorder (AUDIT), trait anxiety (STAI-T), depression (Zung SDS), obsessive-compulsive disorder (OCI-R), schizotypy (Short Scales for Schizotypy; SCZ), apathy (AES), impulsivity (BIS-11), and social anxiety (LSAS). Re-estimating this three-factor structure in our sample gave loadings qualitatively consistent with the original factor structure, with the AD factor characterized by symptoms of trait anxiety, depression, apathy, and impulsivity, the CI factor characterized by symptoms of obsessive compulsive disorder (OCD), eating disorders, and alcohol use disorder, and the SW factor characterized by symptoms of social anxiety disorder. These three factors represent dissociable dimensions of psychopathology with distinct hypothesized relationships to core computational processes: AD maps onto internalizing symptoms theorized to involve heightened sensitivity to negative outcomes or altered motivational states; CI onto compulsive and intrusive cognition that may reflect aberrant uncertainty processing; and SW onto social-evaluative concerns linked to motivational withdrawal.

Together, the multi-attribute gain–loss decision task and 3-factor psychopathology model enable quantitative measures of individual-level economic preferences—for reward, risk, and non-instrumental information—and psychiatric symptomatology in a large population of participants.

### 2.2 Economic preferences for reward, risk, and information in gain and loss

We first set out to characterize how participants trade off reward, risk, and information in gain and loss contexts, establishing the baseline population pattern against which to examine individual variability. We applied a standard decision-theoretic approach of fitting psychometric functions describing how population choice preferences scale with reward and are shifted by the presence of reward uncertainty or the opportunity to gain information in gain or loss context. We confirmed that participants showed expected patterns of reward sensitivity, reliably scaling the percentage of choosing the offer (Right vs Left) based on the difference in expected reward (E[Rew]) in both gain and loss contexts (Figure 2A; E[Rew] slope: gain 0.835 ± 0.011 (*z* = +72.7), loss 0.746 ± 0.013 (*z* = +56.9), both *p*_BH_ *<* 0.0001; here and below, estimates are reported ± standard error (SE), with two-tailed cluster-robust Wald tests and Benjamini–Hochberg false-discovery-rate (BH–FDR) corrected p-values, Methods). To understand how participants traded E[Rew] off against the risk associated with reward uncertainty (Unc[Rew]), we examined the psychometric function predicting the choice of Risky (2 possible outcomes differ maximally) vs Certain (1 guaranteed outcome) offers as a function of their difference in E[Rew]. This analysis showed a tendency to trade Unc[Rew] off with E[Rew], with participants preferring Certain offers in gain and Risky offers in loss (Figure 2A; Unc[Rew] intercept: gain −0.561 ± 0.026 (*z* = −21.3), loss +0.144 ± 0.026 (*z* = +5.5), both *p*_BH_ *<* 0.0001). This is the risk-reflection effect that is classically observed and studied in behavioral economics: the risk aversion that typically governs choices among gains reverses toward risk seeking when the same choices are framed as losses [6, 33]. Finally, to assess trade offs between the opportunity to gain advance information (Info) and E[Rew], we examined the psychometric function predicting the choice of Info vs Noinfo offers as a function of their difference in E[Rew] and observed a strong preference for Info in both contexts (Figure 2A; Info intercept: gain +1.041 ± 0.018 (*z* = +59.1), loss +0.604 ± 0.020 (*z* = +30.9), both *p*_BH_ *<* 0.0001). Analyzing raw choice percentages showed similar results: the percentage of choices favoring the offer higher in E[Rew] was significantly above 50% in both contexts, as was the percentage favoring the more informative (Info) offer, whereas the percentage favoring the offer higher in Unc[Rew] reversed from significantly below 50% in gain to significantly above 50% in loss (Figure 2B; E[Rew]: 75.0% ± 0.26 in gain, 73.2% ± 0.32 in loss, both *t*(1953) *>* 72, both *p <* 0.0001; Unc[Rew]: 39.8% ± 0.40 in gain, *t*(1953) = −25.4, *p <* 0.0001; 53.1% ± 0.44 in loss, *t*(1953) = +7.1, *p <* 0.0001; Info: 71.7% ± 0.31 in gain, 63.5% ± 0.40 in loss; both *t*(1953) *>* 33, both *p <* 0.0001; one-sample *t*-tests vs. 50%). Thus, as a population, decision-makers were reward-seeking, risk-averse in gain and risk-seeking in loss, and information-seeking, consistent with prior work [6, 21, 23, 33].

**Figure 2.**
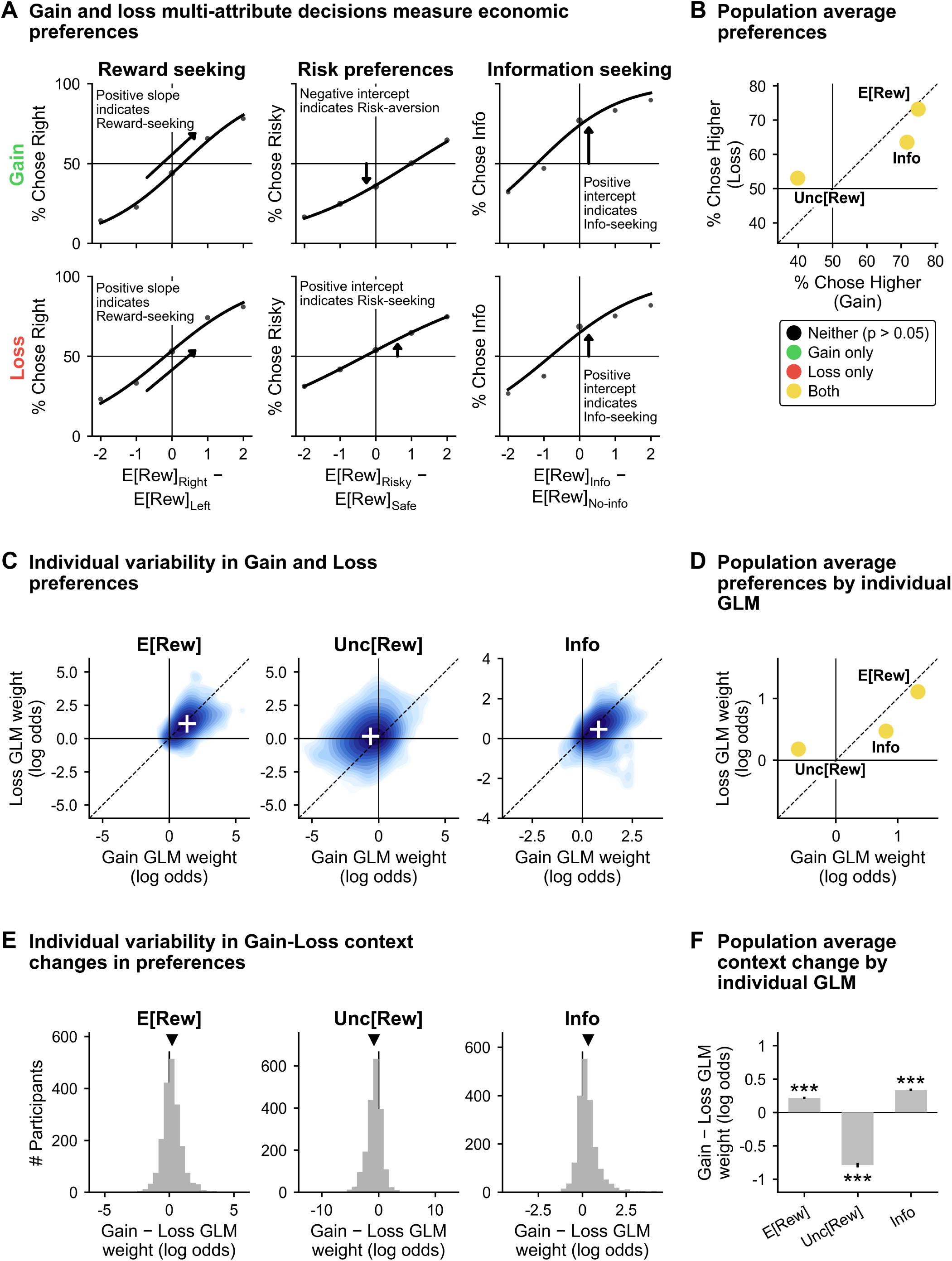
Economic preferences for reward, risk, and information in gain and loss contexts. Across the population (n=1954), all three economic preferences—for reward, risk, and non-instrumental information— differed systematically between gain and loss contexts: reward and information preferences were positive in both but weaker in loss, risk preference reversed sign (the classical reflection effect), and substantial individual heterogeneity persisted around all of these population averages, including a notable subgroup of individuals who reversed their information preference between contexts—establishing the rich preference landscape that psychopathology analyses must explain. Throughout this figure the loss-context E[Rew] measure is sign-flipped (scored on the higher-reward, i.e. less negative, offer) so that a positive value means reward-seeking in both contexts; Unc[Rew] and Info are not flipped. **(A)** Population psychometric curves for three choice comparisons, one per column (column headings *Reward seeking*, *Risk preferences*, *Information seeking*), with the Gain context on the top row (green row label) and Loss on the bottom row (red). The three comparisons are right vs. left offers (all trials; y-axis % Chose Right), risky vs. safe offers (trials pairing a certain with a maximally risky offer; % Chose Risky) and Info vs. Noinfo offers (trials pairing an informative with an uninformative offer; % Chose Info). In each tile choice probability is plotted against the difference in expected reward between the two offers (x-axis E[Rew]_Right_ - E[Rew]_Left_, E[Rew]_Risky_ - E[Rew]_Safe_, E[Rew]_Info_ - E[Rew]_No-info_; offers were not otherwise matched, Methods); gray dots are binned data and the black curve is a 2-parameter logistic fit (slope and intercept; no lapse). Horizontal and vertical reference lines mark 50% choice and a zero reward difference. One annotation per tile names the coefficient it marks: in the reward column an arrow drawn along the fitted tangent at zero marks the *slope*, and in the other two columns a vertical arrow running from the 50% line to the fitted curve marks the *intercept*, the choice bias on trials where the two offers are matched in expected reward. Gain and loss functions diverge most strongly for Unc[Rew], whose intercept reverses sign across contexts (gain −0.561 ± 0.026; loss +0.144 ± 0.026; both *p*_BH_ *<* 0.0001), and for Info, which stays positive but weakens (gain +1.041 ± 0.018; loss +0.604 ± 0.020; both *p*_BH_ *<* 0.0001), while E[Rew] sensitivity remains positive in both (gain slope 0.835 ± 0.011; loss slope 0.746 ± 0.013; both *p*_BH_ *<* 0.0001); p-values are two-tailed and BH–FDR corrected over the 24-test psychometric family shared with Figure 3B (6 population + 18 tertile tests; Methods). Note on the intercept arrows: an arrow’s length is the intercept it reads only ab ove a minimum drawn length of 9 percentage points; the loss risk-preference intercept is smaller than this (+0.144 log odds = +3.6 percentage points above 50%), so that one arrow is drawn at the floor and shows direction only, with its value readable from the curve and its axis. **(B)** Population average preferences: proportion of trials on which individuals chose the offer higher in E[Rew], Unc[Rew] or Info, computed over trials on which the two offers differed in that attribute (Methods), averaged across individuals and plotted as % Chose Higher in gain (x-axis) vs. loss (y-axis). Error bars show standard error of the mean (SE) across individuals (smaller than the drawn markers here); the dashed line is unity and the solid lines mark 50%. Point color indicates a significant difference from 50% in gain (green), loss (red), both (yellow) or neither (black) by one-sample *t*-tests, BH–FDR corrected over the 6 attribute × context tests. All three attributes are yellow: E[Rew] 75.0% ± 0.26 in gain (*t*(1953) = +95.0) and 73.2% ± 0.32 in loss (*t* = +72.4), Info 71.7% ± 0.31 in gain (*t* = +69.0) and 63.5% ± 0.40 in loss (*t* = +33.9, *p*_BH_ *<* 0.0001), and Unc[Rew] reversing from 39.8% ± 0.40 in gain (*t* = −25.4, *p*_BH_ *<* 0.0001) to 53.1% ± 0.44 in loss (*t* = +7.1, *p*_BH_ *<* 0.0001); all remaining *p*_BH_ *<* 0.0001. **(C)** Individual variability in gain and loss preferences: one tile per attribute (E[Rew], Unc[Rew], Info) showing a filled log-density contour (2D kernel density estimate) of the n=1954 individual GLM weights in gain (x-axis) vs. loss (y-axis), both in log odds. The white crosshair is the population mean ± SEM (standard error of the mean) of those weights, the dashed line is unity and the solid lines mark zero. Heterogeneity readouts are omitted from the figure for space; their values are, as observed SD of individual weights (SD_obs_) vs. the SD expected from estimation noise alone (SD_noise_), *I*^2^ and Cochran’s *Q* (Methods): E[Rew] gain 0.84, 0.35, 84%, *Q*(1953) = 12433 and loss 0.83, 0.30, 88%, *Q* = 16050; Unc[Rew] gain 1.41, 0.40, 90%, *Q* = 19459 and loss 1.26, 0.32, 91%, *Q* = 20600; Info gain 0.62, 0.21, 88%, *Q* = 16093 and loss 0.66, 0.18, 91%, *Q* = 21640; every test rejects pure-noise heterogeneity (*p <* 0.0001), so individual differences in preference strength are genuine and not merely estimation noise. The density fields also show directional heterogeneity: a small but visible protrusion of individuals had individually significant negative Info weights in loss (192/1954, 9.8%, vs. 2.5% chance, exact binomial *p <* 0.0001), and Unc[Rew] weights varied in direction as well as strength within a context (in gain, 1027/1954 (52.6%) individually negative and 302/1954 (15.5%) individually positive; both binomial *p <* 0.0001). **(D)** Population average preferences by individual GLM: the mean across individuals of the same individual GLM weights, in gain (x-axis) vs. loss (y-axis) log odds, with SE bars, a dashed unity line and zero rules. Point color is as in **(B)**, from one-sample *t*-tests against zero BH–FDR corrected over this panel’s 12 regressor × context tests (6 regressors × 2 contexts; Methods); all three points are yellow. E[Rew] +1.327 ± 0.019 in gain and +1.111 ± 0.019 in loss (*t*(1953) = +69.6, +59.3); Info +0.808 ± 0.014 and +0.470 ± 0.015 (*t* = +58.1, +31.7); Unc[Rew] changing sign from −0.610 ± 0.032 in gain (*t* = −19.1) to +0.180 ± 0.028 in loss (*t* = +6.3); all *p*_BH_ *<* 0.0001. **(E)** Individual variability in gain–loss context changes: one histogram per attribute of the individual gain-minus-loss difference in GLM weights across all n=1954 participants (gray bars; x-axis Gain - Loss GLM weight in log odds, y-axis number of participants), with a solid vertical line at zero and a black downward triangle marking the population mean. Heterogeneity of these differences (same four quantities as in **(C)**): E[Rew] SD_obs_ 0.83, SD_noise_ 0.46, *I*^2^ 66%, *Q*(1953) = 5714; Unc[Rew] 1.54, 0.52, 89%, *Q* = 17265; Info 0.71, 0.27, 81%, *Q* = 10266; all *p <* 0.0001. Nontrivial numbers of individuals shifted in the direct<u>ion opposite to</u> the population mean by an individual-level Wald test on the standardized contrast 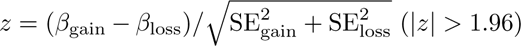: for Unc[Rew], 221/1954 (11.3%) were significantly gain *>* loss (exact binomial vs. 2.5% chance, *p <* 0.0001), and for Info, 87/1954 (4.5%) were significantly loss *>* gain (*p <* 0.0001). These counts are stated here rather than printed on the figure. **(F)** Population average context change by individual GLM: bars showing the population-average gain-minus-loss difference in individual GLM weights, with SE bars; *, **, *** indicate *p <* 0.05, 0.01, 0.001 from one-sample *t*-tests against zero, BH–FDR corrected over the 6 regressor difference tests of this panel. All three are significant: E[Rew] +0.216 ± 0.019 (*t*(1953) = +11.5, *p*_BH_ *<* 0.0001), Unc[Rew] −0.790 ± 0.035 (*t* = −22.8, *p*_BH_ *<* 0.0001) and Info +0.338 ± 0.016 (*t* = +21.0, *p*_BH_ *<* 0.0001), confirming that the gain/loss framing meaningfully alters the weight placed on each attribute. Takeaway: individuals on average preferred higher-reward and more-informative offers in both contexts and reversed from risk-aversion in gain to risk-seeking in loss, but each of these average preferences coexists with substantial individual heterogeneity, including a notable density of individuals who reverse their information preference between gain and loss.

To move beyond population-level summaries of choice preferences and quantify how individuals trade off multiple attributes concurrently, we fit individual-level logistic generalized linear models (GLMs) of choice for each participant, estimating separate weights for each economic attribute in gain and loss (Figure 2C–D). We first confirmed that this analysis produced similar aggregate-level results as the classic choice preference analyses by examining population averages of these individually-fit GLMs. Indeed, results were highly comparable, with decision-makers being reward-seeking, risk-averse in gain and risk-seeking in loss, and information-seeking (Figure 2D; all six regressors in Supplementary Figure 1A): E[Rew] +1.327 ± 0.019 in gain and +1.111 ± 0.019 in loss (*t*(1953) = +69.6 and +59.3), Info +0.808 ± 0.014 and +0.470 ± 0.015 (*t* = +58.1 and +31.7), and Unc[Rew] reversing from −0.610 ± 0.032 (*t* = −19.1) to +0.180 ± 0.028 (*t* = +6.3) (one-sample *t*-tests against zero, all *p*_BH_ *<* 0.0001). Then, we assessed heterogeneity in preferences across the population by examining the dispersion of individual-level GLM weights and found evidence of substantial variation across individuals in economic preferences (Figure 2C; all six regressors in Supplementary Figure 1D). This variation was significantly greater than what would be expected simply by estimation noise (Cochran’s *Q* tests against pure-noise heterogeneity, for the three attributes displayed in Figure 2C: E[Rew] gain *Q*(1953) = 12433, *I*_2_ = 84.3%; E[Rew] loss *Q*(1953) = 16050, *I*_2_ = 87.8%; Unc[Rew] gain *Q*(1953) = 19459, *I*_2_ = 90.0%; Unc[Rew] loss *Q*(1953) = 20600, *I*_2_ = 90.5%; Info gain *Q*(1953) = 16093, *I*_2_ = 87.9%; Info loss *Q*(1953) = 21640, *I*_2_ = 91.0%; all *p <* 0.0001; the corresponding statistics for the Info×Unc[Rew] interaction are reported with Figure 4B below and in the Supplementary Figure 1 caption). While individuals were overwhelmingly reward-seeking, there was substantial variation in the strength of this preference. Heterogeneity was particularly pronounced for Unc[Rew], which varied across individuals in both strength and direction within each context. In gain, while individuals were mostly risk-averse, with 1027 individuals (52.6%) having significantly negative Unc[Rew] weights, 302 (15.5%) had significantly positive ones (individual significance taken as twotailed *p <* 0.05 from each participant’s own GLM here and throughout; exact binomial tests against the 2.5% expected per direction by chance, both *p <* 0.0001). In loss, there was a weak tendency toward risk-seeking 736 (37.7%), but with a strong minority who were risk-averse 531 (27.2%) (both *p <* 0.0001, exact binomial tests vs. 2.5%). Finally, individuals were largely information-seeking across contexts, but again with substantial variation in strength. Choices were driven by the offers’ economic value rather than by trial history: in a population model carrying both, the value difference between the offers dominated choice (+1.05 log odds per unit in each context) whereas the tendency to repeat the previous choice, though detectable, was over forty-fold smaller (+0.018 and +0.026, gain and loss), the previous trial’s reward-prediction-error terms were smaller still, and no psychopathology factor interacted with any trial-history term (all *p*_BH_ ≥ 0.12; full statistics in Supplementary Figure 2). Thus, overall, our approach captured substantial variation in economic preferences in each context.

We next examined how preferences shifted between contexts. While risk-reflection is an expected pattern, it is less clear how reward and information preferences differ between gain and loss [21, 22, 34]. On average, reward-seeking was stronger in gain, the risk preference flipped from aversion in gain to seeking in loss, and information-seeking was stronger in gain on average, with GLM weights for all three attributes showing significant average gain–loss differences across the population (E[Rew] +0.216±0.019 (*t*(1953) = +11.5), Unc[Rew] −0.790±0.035 (*t*(1953) = −22.8) and Info +0.338±0.016 (*t*(1953) = +21.0); one-sample *t*-tests of the gain-minus-loss differences against zero, all *p*_BH_ *<* 0.0001; Figure 2F).

Next, we examined individual variability in how preferences change between gain and loss (Figure 2E). Just as individual preferences in each context exhibited substantial heterogeneity, so did individual changes in these preferences. While reward-seeking tended to be stronger in gain, the strength of this difference varied, and some individuals even exhibited stronger reward-seeking in loss (126 individuals, 6.4%, significantly loss*>*gain, more than the 2.5% expected by chance, *p <* 0.0001, binomial test). Individuals tended to become more risk-averse in gain, but the magnitude of shift in this preference varied, and a subset of individuals shifted toward stronger risk-seeking in gain relative to loss (221; 11.3% vs. 2.5% expected; *p <* 0.0001, binomial test). Information-seeking, like reward, tended to be stronger in gain but with variations in strength, and 87 individuals (4.5% vs. 2.5% expected) were significantly stronger in loss (*p <* 0.0001, binomial test). However, we also noted a relative abundance of individuals who were information-seeking in gain but information-averse in loss, with 130 of 1954 individually significant (6.7% vs. the 0.0625% expected by chance, *p <* 0.0001, binomial test; Supplementary Figure 3A), reminiscent of risk preferences reflecting across contexts.

In summary, individuals were on average strongly reward- and information-seeking in both contexts, became more risk-seeking in loss, and showed weaker reward and information preferences in loss than in gain. At the same time, individual differences in preference strength and even in the sign of context-shifts were substantial, including a density of individuals exhibiting an information-seeking to information-averse reversal across contexts.

### 2.3 Alterations in economic preferences in gain and loss associated with psychopathology

Having established that all three preference attributes—for reward, risk, and information—differed systematically between gain and loss and varied substantially across individuals, we asked whether these individual differences could be partially explained by differences in psychiatric symptoms. If psychopathological symptomatology is linked to distinct alterations in decision-related computations or processes, then psychopathological factors such as AD, CI, and SW may show distinct patterns of economic preferences in gain and loss.

To assess how economic preference variation between individuals may be associated with psychiatric symptoms, we examined whether and how groups of individuals high or low in each psychopathology factor differed in their preferences for reward, risk, and information across gain and loss contexts. We first compared individuals high vs. low in each factor—the top vs. bottom tertile of that factor’s scores, *n* = 651 per group—by fitting population psychometric functions separately for each group (Figure 3B), with example matched single-subject fits illustrating each pattern at the level of individual participants (Figure 3A). We observed a set of dissociable alterations across factors (all high-vs-low parameter comparisons below are two-tailed Wald tests; Methods). Individuals high in AD were more sensitive to reward magnitude than their low-AD counterparts in both gain and loss, with steeper E[Rew] psychometric slopes (gain 0.913 vs. 0.783, *p*_BH_ *<* 0.0001; loss 0.838 vs. 0.680, *p*_BH_ *<* 0.0001), but with no detectable difference in either the Unc[Rew] intercept or the Info intercept in either context (all *p*_BH_ *>* 0.05). High-CI individuals showed the opposite pattern for reward—weaker reward sensitivity in both contexts (E[Rew] slope: gain 0.754 vs. 0.886, *p*_BH_ *<* 0.0001; loss 0.654 vs. 0.810, *p*_BH_ *<* 0.0001)—together with significantly weaker risk aversion specifically in gain (Unc[Rew] intercept: −0.395 vs. −0.704, *p*_BH_ *<* 0.0001), and a shift away from information-seeking specifically in loss (Info intercept: 0.520 vs. 0.668, *p*_BH_ = 3.3 × 10^-3^). Like AD, high-SW individuals were more reward-sensitive across both contexts (E[Rew] slope: gain 0.891 vs. 0.795, *p*_BH_ = 1.1 × 10^-3^; loss 0.827 vs. 0.698, *p*_BH_ *<* 0.0001), but, unlike AD or CI, they showed increased risk-aversion in gain (Unc[Rew] intercept: −0.647 vs. −0.493, *p*_BH_ = 2.6 × 10^-2^) and significantly stronger information-seeking in loss (Info intercept: 0.680 vs. 0.578, *p*_BH_ = 4.4 × 10^-2^).

**Figure 3.**
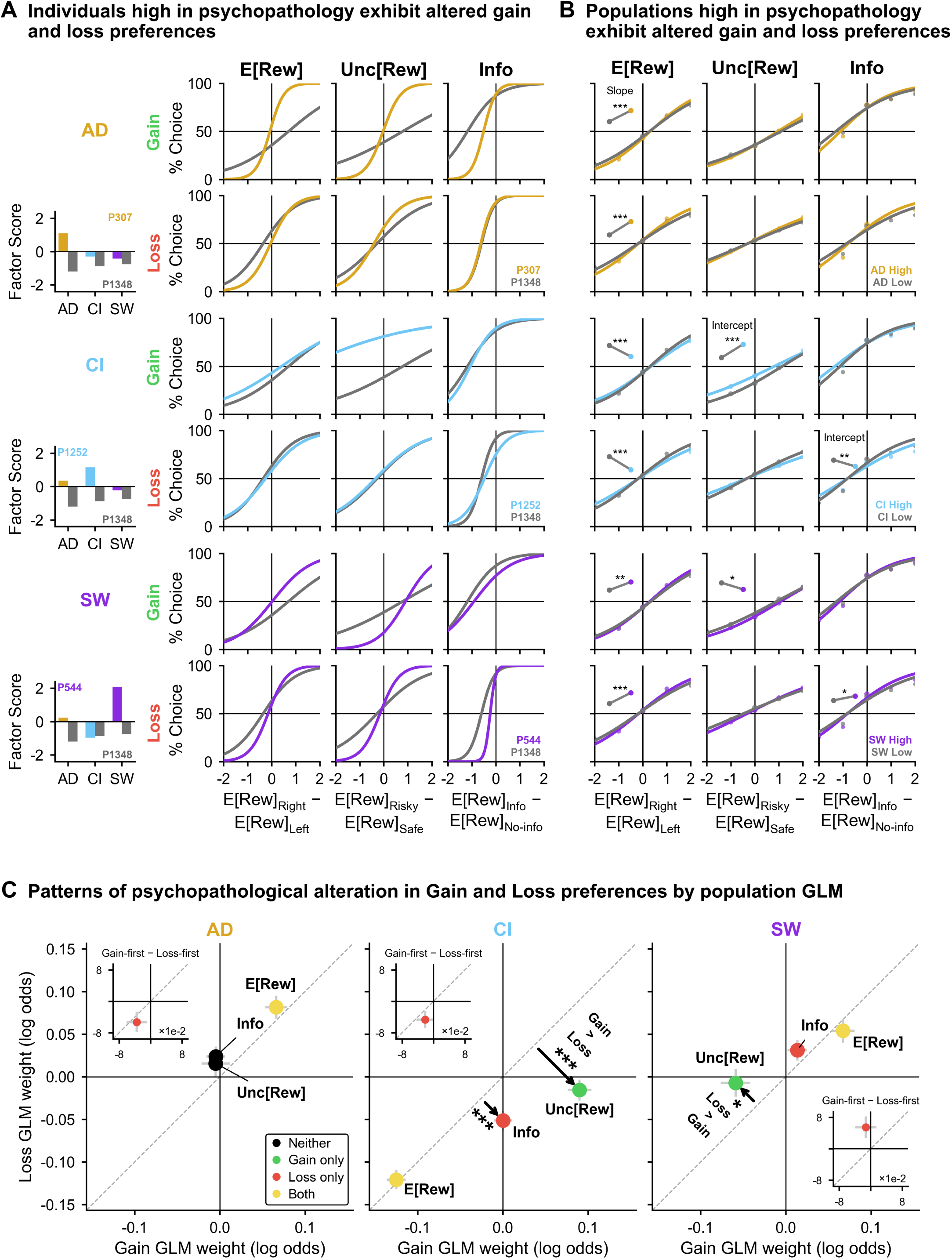
Dissociable alterations in gain and loss economic preferences associated with psychopathology. Linking economic preferences to dimensional psychopathology reveals three dissociable patterns: AD and SW are associated with heightened reward sensitivity across both contexts while CI is associated with broadly reduced reward sensitivity; in addition, CI is associated with reduced information-seeking in loss and reduced risk-aversion in gain, whereas SW shifts risk and information preferences in the opposite direction—a pattern of alterations too specific to be explained by a simple global scaling account and pointing to CI as the dimension most distinctively linked to information preferences. **(A)** Individuals high in psychopathology exhibit altered gain and loss preferences. Six rows pair each psychopathology factor (AD, CI, SW) with each context (Gain, Loss), and the three columns are the attribute psychometric functions of **Figure 2A** (E[Rew], Unc[Rew], Info; same x-axes, shared y label % Choice). In every tile one selected high-factor individual is drawn in that factor’s color and a single matched control individual in gray. The bar plot at the left of each factor block gives the two individuals’ AD, CI and SW factor scores and their participant indices (P307 vs. control P1348 for AD, P1252 vs. P1348 for CI, P544 vs. P1348 for SW). These examples show that the population-level dissociations in **(B)** are visible at the level of single participants, consistent with the aggregate patterns reflecting real individual differences rather than artifacts of averaging. **(B)** Populations high in psychopathology exhibit altered gain and loss preferences. Same 6 × 3 grid of rows and columns as **(A)**, now with population psychometric curves for the top vs. bottom tertile of each factor: the high-factor tertile is drawn in the factor’s color (AD gold, CI light blue, SW violet) and the low-factor tertile in gray, with binned points and 2-parameter logistic fits (slope and intercept; no lapse). Context identity is carried by the row labels (Gain green, Loss red) rather than by curve color. An inset in the upper-left of a tile shows the high vs. low tertile group estimates of that tile’s designated parameter as a paired dot–line–dot plot (low group gray, high group in the factor color) with cluster-robust SE error bars; the parameter is named on each column’s first surviving inset (*Slope* for the E[Rew] column, *Intercept* for the Unc[Rew] and Info columns). Insets are drawn only where the high-vs-low comparison survives correction, so a tile without an inset is a tile whose test was not significant. P-values are from two-tailed Wald tests, BH–FDR corrected jointly over the 24-test psychometric family (6 population tests from **Figure 2A** + 18 tertile contrasts, one designated parameter per tile; Methods); *, **, *** indicate *p*_BH_ *<* 0.05, 0.01, 0.001. AD-high individuals showed steeper E[Rew] slopes than AD-low in both contexts (gain 0.913 vs. 0.783, *p*_BH_ *<* 0.0001; loss 0.838 vs. 0.680, *p*_BH_ *<* 0.0001) but no detectable change in risk preference or information preference in either context (all *p*_BH_ *>* 0.05; the closest is the AD loss Info intercept, *p*_BH_ = 0.078). CI-high individuals showed the opposite pattern for reward (gain 0.754 vs. 0.886, *p*_BH_ *<* 0.0001; loss 0.654 vs. 0.810, *p*_BH_ *<* 0.0001), significantly weaker risk aversion in gain (Unc[Rew] intercept −0.395 vs. −0.704, *p*_BH_ *<* 0.0001) and a shift away from information-seeking in loss (Info intercept 0.520 vs. 0.668, *p*_BH_ = 3.3 × 10^-3^). SW-high individuals showed steeper reward sensitivity in both contexts (gain 0.891 vs. 0.795, *p*_BH_ = 1.1 × 10^-3^; loss 0.827 vs. 0.698, *p*_BH_ *<* 0.0001), increased risk-aversion in gain (Unc[Rew] intercept −0.647 vs. −0.493, *p*_BH_ = 2.6 × 10^-2^) and significantly stronger information-seeking in loss (Info intercept 0.680 vs. 0.578, *p*_BH_ = 4.4 × 10^-2^). **(C)** Patterns of psychopathological alteration by population GLM: scatters of the population (cluster-robust) logistic GLM interactions of E[Rew], Unc[Rew] and Info with each psychopathology factor, one panel per factor (AD, CI, SW), plotted in gain (x-axis) vs. loss (y-axis) log odds on one shared square scale. Coefficients are decomposed from the joint population model into context-specific estimates (*β*_gain_ = *β*_main_ + 0.5 · *β*_context_, *β*_loss_ = *β*_main_ - 0.5 · *β*_context_) with SEs propagated using the full coefficient covariance matrix; gray arms show cluster-robust SE and the gray dashed line is unity. Point color marks a two-tailed Wald test against zero that is significant in gain (green), loss (red), both (yellow) or neither (black), BH–FDR corrected over the population model’s 48 context-decomposed effects (Methods); quoted p-values are raw unless labeled *p*_BH_. A black arrow drawn from the identity line to a point marks an attribute × context × factor interaction that survives BH–FDR correction over that 12-test family (4 attributes × 3 factors; Methods), and carries the test’s star tier plus, once per panel, a bold note giving the direction of the difference (*Gain > Loss* in the CI panel, *Loss > Gain* in the SW panel). Three such terms survive: Unc[Rew] × context × CI (*β* = +0.106, *z* = 6.94, *p*_BH_ *<* 0.0001), Info × context × CI (*β* = +0.052, *z* = 3.83, *p*_BH_ = 7.8 × 10^-4^) and Unc[Rew] × context × SW (*β* = −0.052, *z* = −2.71, *p*_BH_ = 2.7 × 10^-2^). The dissociations of **(A)** and **(B)** are recapitulated here: AD and SW interacted positively with E[Rew] in both contexts (AD gain *β* = +0.066, *p <* 0.0001; loss +0.082, *p <* 0.0001; SW gain +0.067, *p <* 0.0001; loss +0.054, *p* = 1.2 × 10^-4^), whereas CI interacted negatively in both (gain −0.125; loss −0.121; both *p <* 0.0001); in the AD panel the Unc[Rew] and Info points sit at the origin and are black (both n.s.). For Unc[Rew], CI interacted positively in gain—i.e. less risk-averse—(+0.090, *p <* 0.0001) but not in loss (−0.016, *p* = 0.21), which is the context dependence marked by the arrow in the CI panel, while SW interacted negatively—i.e. more risk-averse—specifically in gain (−0.059, *p* = 6.6 × 10^-4^), marked by the arrow in the SW panel. For Info, CI interacted negatively specifically in loss (−0.051, *p <* 0.0001), marked by the second CI arrow, whereas SW interacted positively in loss (+0.031, *p* = 0.011). An inset in each panel, titled *Gain-first* - *Loss-first* and drawn on ×10^-2^ axes, is a task-order robustness check for that panel’s own E[Rew] × factor interaction: the gain-first and loss-first cohorts were fit as separate submodels and their coefficients differenced within the gain context (x-axis) and within the loss context (y-axis), with color from an independent-variance two-tailed Wald test (raw, uncorrected p). All three task-order differences were small relative to the corresponding main interactions (|Δ*β*| ≤ 0.055, all *p* ≥ 0.038, each nominally significant in the loss context), indicating that the psychopathology effects are not primarily an artifact of task order. Takeaway: AD, CI and SW map onto dissociable, partially opposing patterns of alteration in reward, risk and information preferences across gain and loss contexts, with CI standing out for the breadth and direction of its effects (weaker reward, less risk-averse in gain, less info-seeking in loss) and SW shifting risk and information preferences in the opposite direction.

To model these group-level effects jointly while controlling for the covariance among economic preferences and psychopathology factors, we fit a population GLM in which all three psychopathology factors entered as continuous covariates and were allowed to interact with the economic attributes and with context (Figure 3C; Methods). Mixed models with individual random effects estimated comparable fixed effects (Supplementary Figure 4; the factor interactions are shown per regressor, for all six regressors, in Supplementary Figure 5). To assess economic changes associated with each psychopathological factor in each context, we recombined linear weights to define gain or loss effects (Methods). These gain and loss effects estimated by the population model recapitulated the group-level dissociations (two-tailed cluster-robust Wald tests; p-values reported raw, and every effect called significant here also survives BH correction; Methods). AD interacted positively with E[Rew] across both contexts (E[Rew]×AD: gain *β* = +0.066, *z* = +5.11, *p <* 0.0001; loss *β* = +0.082, *z* = +6.09, *p <* 0.0001), as did SW (E[Rew]×SW: gain *β* = +0.067, *z* = +4.99, *p <* 0.0001; loss *β* = +0.054, *z* = +3.84, *p* = 1.2 × 10^-4^), indicating stronger reward-seeking with higher AD or SW psychopathology. Conversely, CI interacted negatively with E[Rew] in both contexts (E[Rew]×CI: gain *β* = −0.125, *z* = −11.26; loss *β* = −0.121, *z* = −10.70; both *p <* 0.0001), indicating weaker reward-seeking with higher CI psychopathology. For Unc[Rew], CI interacted positively in gain, (Unc[Rew]×CI: gain *β* = +0.090, *z* = +6.49, *p <* 0.0001), indicating less risk-aversion with higher CI psychopathology in gain, while SW interacted negatively in gain (Unc[Rew]×SW: gain *β* = −0.059, *z* = −3.41, *p* = 6.6 × 10^-4^), indicating more risk-aversion with higher SW psychopathology in gain. For Info, CI interacted negatively in loss (Info×CI loss: *β* = −0.051, *z* = −4.47, *p <* 0.0001), indicating weaker information-seeking with higher CI psychopathology whereas SW interacted positively in loss (Info×SW loss: *β* = +0.031, *z* = +2.54, *p* = 0.011), indicating stronger information-seeking with higher SW psychopathology. To ask which of these factor effects were themselves significantly context-dependent, we analyzed the attribute×context×factor interactions. CI’s shift in risk preference was strongly context-dependent (Unc[Rew]×context×CI: *β* = +0.106, *z* = 6.94, *p*_BH_ *<* 0.0001; Figure 3C, middle panel), as was its shift in information preference (Info×context×CI: *β* = +0.052, *z* = 3.83, *p*_BH_ = 7.8 × 10^-4^), while SW’s risk-preference effect was significantly stronger in gain than in loss (Unc[Rew]×context×SW: *β* = −0.052, *z* = −2.71, *p*_BH_ = 2.7 × 10^-2^). Thus the two factors with context-specific risk and information effects, CI and SW, are associated with these attributes in a genuinely context-dependent fashion rather than uniformly across framings. Psychopathology effects on reward-seeking in loss differed modestly but significantly between participants who performed gain first and those who performed loss first, which could suggest a difference in change-point detection [72, 73] (Figure 3C, E[Rew] inset; AD −0.053, *z* = −2.01, *p* = 0.044; CI −0.046, *z* = −2.07, *p* = 0.038; SW +0.055, *z* = +1.99, *p* = 0.047; Wald tests). Similar directions of preference alteration are broadly visible when participants are split (median split, high vs. low) on the individual symptom questionnaires rather than on the three factors, and separately within each task-order cohort (Supplementary Figure 6): the CI-loaded instruments (OCI-R, EAT-26) show the reward-sensitivity reduction in both cohorts, whereas the AD-loaded instruments show the corresponding elevation most clearly in the loss-first cohort, consistent with raw questionnaire scores mixing variance from factors with opposing associations. The factor-level associations also survive controlling for questionnaire engagement (Supplementary Figure 7).

To further assess whether any of these effects could be due to generalized psychomotor or engagement differences instead of true changes in economic behavior, we examined task reaction speeds. Choice reaction speed tracked the economic choice variables and recent reward prediction errors—faster with larger total offer value and larger previous-trial RPE, slower after larger previous outcomes (all *p*_BH_ *<* 0.0001 in both contexts; Supplementary Figure 8)—as expected if participants were engaging in economic decisions [74, 75]. In contrast, the speed of the trial-initiating fixation response was minimally related to task variables and showed no association with any psychopathology factor (all interactions *p*_BH_ ≥ 0.056; Supplementary Figure 9), and the post-outcome collection response was related to neither task variables (smallest *p*_BH_ = 0.42) nor psychopathology (no factor term approached significance, smallest *p*_BH_ = 0.77; Supplementary Figure 10), further suggesting that changes in behavior are not a by-product of generalized psychomotor or engagement differences.

Together, these analyses suggest that AD, CI, and SW psychopathology factors map onto dissociable patterns of alteration in economic preferences across gain and loss contexts. The SW profile resembles AD in its reward-sensitivity elevation but diverges in the risk and information dimensions suggesting that social-withdrawal symptoms track a distinct, if partially overlapping, preference phenotype. CI preference changes largely opposed AD and SW, with opposite reward sensitivity from both and shifting risk and information preferences in the opposite direction from CI.

### 2.4 Scaling of information value with reward uncertainty in gain and loss

The analyses above examined how strongly participants valued advance information overall. However, our previous work showed that information value is itself computed as a function of other attributes of the anticipated outcome. In particular, reward uncertainty strongly scales the subjective value assigned to information: humans and monkeys were willing to pay more for information when it resolved greater uncertainty, and analogous uncertainty-dependent information signals were observed in the lateral habenula and pallidum [23]. This scaling was not simply a consequence of preference for uncertainty itself, suggesting that it reflects a characteristic computation governing the value assigned to information. We therefore asked whether this computation is preserved across gain and loss contexts and whether its strength varies across individuals and with psychopathology.

We first examined how the population choice preferences for information scaled with the level of reward uncertainty to be resolved by this information and found that % choice of Info increased—if modestly in absolute terms—with Unc[Rew] in both contexts, monotonically in gain while in loss the rise was carried by the high-uncertainty level (Figure 4A; Page’s *L* trend test: gain *L* = 23762, *p <* 0.0001; loss *L* = 23621, *p* = 6.4 × 10^-3^; *n* = 1954; gain mean 76.2% at SD= 0 to 77.7% at SD= 4; loss 68.2% to 69.0%). Notably, information preference sat far above the 50% chance level even for certain offers (SD= 0; 76.2% in gain, 68.2% in loss), for which the advance cue resolves no outcome uncertainty. Thus, uncertainty scaling was not the sole determinant of information preference: participants valued information even when there was no uncertainty to resolve, while assigning it progressively more value as reward uncertainty increased. That this scaling persists even in loss—despite weaker average information preference—extends this previously described computation of information value across gain and loss framing; whether its strength differs between contexts is addressed next with individual-level model estimates.

**Figure 4.**
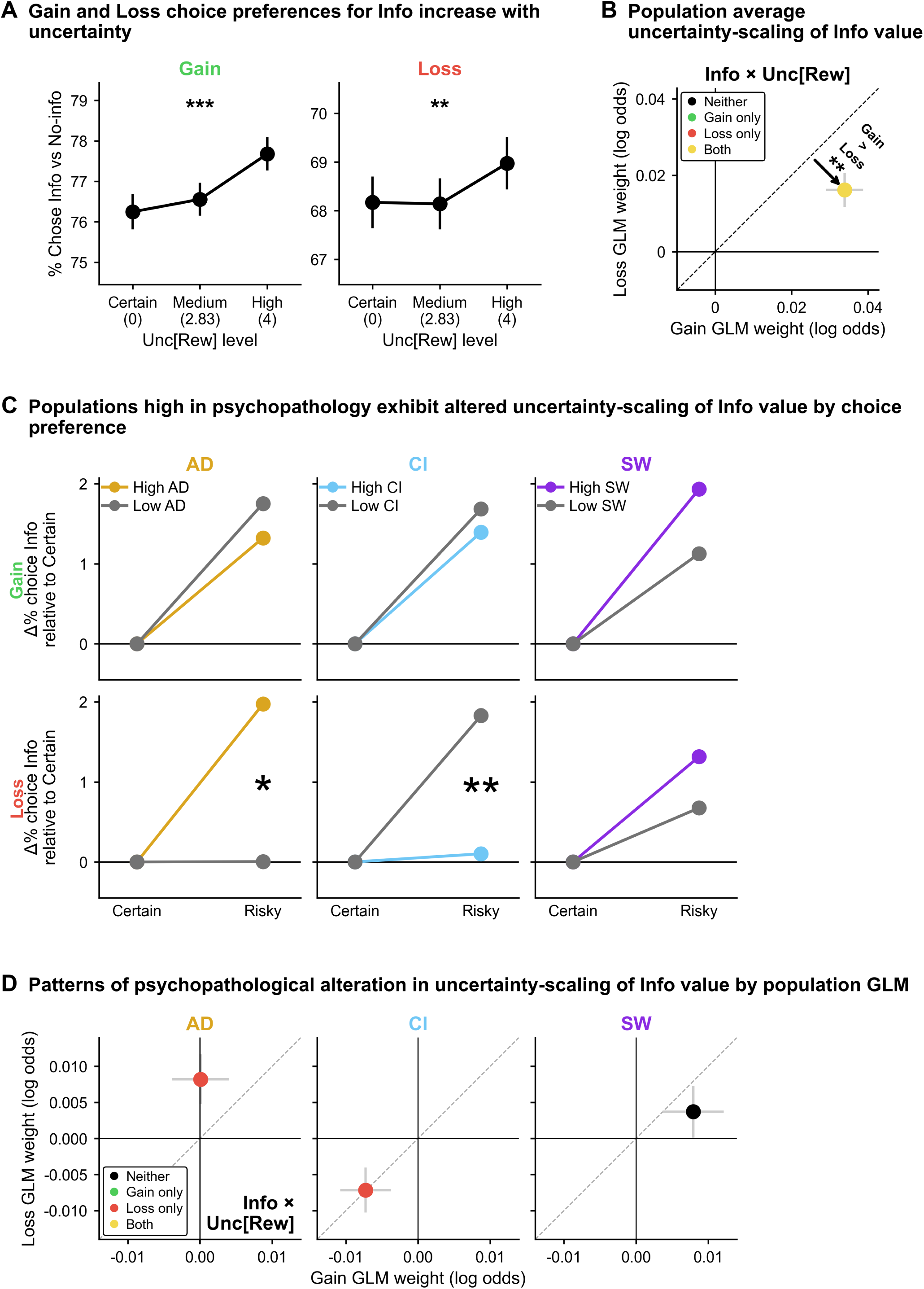
Scaling of information value with reward uncertainty in gain and loss. Information preference scales positively with reward uncertainty in both gain and loss contexts, but more strongly in gain—a context asymmetry with genuine individual variability that parallels the psychopathology-related differences in reward sensitivity: in the loss context, high-AD participants scale information value more strongly and high-CI participants scale it more weakly, deepening the link between AD and CI psychopathology and information processing established in **Figure 3. (A)** Gain and loss choice preferences for Info increase with uncertainty. Two tiles, Gain (green heading, left) and Loss (red heading, right), plot the population average proportion of trials on which individuals chose the more informative offer when the two offers differed only in informativeness (y-axis % Chose Info vs No-info), separated by the reward-distribution width shared by the two offers (x-axis Unc[Rew] level, with ticks Certain (0), Medium (2.83) and High (4), the outcome standard deviations of **Figure 1A**’s three uncertainty designs). Error bars show SE across individuals. Stars above each tile are Page’s L trend test on the per-participant three-point curves, with significance from within-participant permutation of the uncertainty-level labels (10,000 permutations, two-sided, uncorrected; Methods). Information preference was far above chance at every level in both contexts and rose across levels in both (gain *L* = 23762, *p <* 0.0001; loss *L* = 23621, *p* = 0.0064; *n* = 1954); the rise is monotonic in gain (76.2% → 76.6% → 77.7%) whereas in loss the certain and medium levels are indistinguishable and the increase is carried by the high-uncertainty level (68.2% → 68.1% → 69.0%). **(B)** Population average uncertainty-scaling of Info value: the population average of the individual Info × Unc[Rew] GLM weights (panel title *Info* × *Unc[Rew]*), plotted in gain (x-axis) vs. loss (y-axis) log odds with gray SE arms and a dashed unity line; conventions as in **Figure 2D**. The point is yellow: Info × Unc[Rew] scaling is significantly positive in both contexts (gain +0.0339 ± 0.0048, *t*(1953) = +7.02, *p*_BH_ *<* 0.0001; loss +0.0162 ± 0.0045, *t* = +3.61, *p*_BH_ = 3.4 × 10^-4^). A black arrow runs perpendicular from the identity line to the point; its length is the gain-minus-loss difference and it carries that difference’s star tier and a bold *Gain > Loss* note. Scaling is significantly stronger in gain (+0.0177±0.0057, *t*(1953) = +3.09, *p* = 0.0020). **(C)** Populations high in psychopathology exhibit altered uncertainty-scaling of Info value. Model-predicted change in information seeking from the certain to the risky condition for participants high vs. low in each psychopathology factor (columns AD, CI, SW; per-column legend High/Low in the factor color and gray), in gain (top row) and loss (bottom row); y-axis Δ% choice Info relative to Certain, x-axis Certain vs. Risky. Predicted probabilities come from a trial-level cluster-robust logistic GLM with a risky-vs-certain indicator, continuous psychopathology factor scores and their interactions, evaluated at the mean factor score of the above- and below-median groups (*n* = 977 each; Methods); each line is normalized to its own certain-condition value, so the displayed quantity is the change in information seeking caused by introducing reward uncertainty. Stars are the risky × factor interaction term (continuous) in that context, raw and uncorrected, and are drawn at double the paper’s usual star size because each cell carries at most one test; a cell without a star had no significant interaction and nothing is printed. In the loss context, high-AD participants showed a stronger increase in information seeking with uncertainty than low-AD participants (*z* = 2.53, *p* = 0.012) and high-CI participants a weaker increase (*z* = −2.68, *p* = 7.4 × 10^-3^); the corresponding gain-context tests were not significant (AD *z* = −0.59, *p* = 0.56; CI *z* = −0.43, *p* = 0.67) and SW did not differ in either context (gain *z* = 1.12, *p* = 0.26; loss *z* = 0.85, *p* = 0.39). The AD and CI patterns in loss parallel the opposing modulations of reward sensitivity in **Figure 3**. **(D)** Patterns of psychopathological alteration in uncertainty-scaling by population GLM: the population cluster-robust GLM interaction of the Info × Unc[Rew] term with each psychopathology factor, one panel per factor (AD, CI, SW) on one shared square scale, in gain (x-axis) vs. loss (y-axis) log odds; the regressor is named once, in the AD panel. Decomposition, error bars, dashed unity line and the four-class point coloring (BH–FDR over the population model’s 48 context-decomposed effects) are as in **Figure 3C**. AD interacted with Info × Unc[Rew] specifically in loss (+0.008, *p*_raw_ = 1.7 × 10^-2^, *p*_BH_ = 0.034; gain +0.0001, n.s.), CI interacted negatively in loss (−0.007, *p*_raw_ = 2.1 × 10^-2^, *p*_BH_ = 0.041) with a similar nominal effect in gain that does not survive correction (−0.007, *p*_raw_ = 3.8 × 10^-2^, *p*_BH_ = 0.071), and SW was not significant in either context (gain *p*_raw_ = 0.062; loss *p* = 0.30). No arrows are drawn: no Info × Unc[Rew] × context × factor term survives BH–FDR over the same 12-test family used in **Figure 3C** (the closest is AD, *p*_BH_ = 0.17). Takeaway: across the population, information preference scales positively with reward uncertainty in both contexts but more strongly in gain; across-individual variability in this scaling is substantial and partially structured by psychopathology, with high-AD participants scaling more strongly in loss and high-CI participants scaling more weakly, most reliably in loss — paralleling AD’s and CI’s opposing modulations of monetary reward sensitivity.

To more precisely quantify this effect at the level of individuals while controlling for cross-attribute covariances, we used individual-level GLMs that included an Info×Unc[Rew] interaction term. Averaged across individuals, Info×Unc[Rew] weights were positive in both contexts, consistent with a tendency for information value to scale up with reward uncertainty (Figure 4B; one-sample *t*-tests against zero: gain +0.0339 ± 0.0048, *t*(1953) = +7.02, *p*_BH_ *<* 0.0001; loss +0.0162 ± 0.0045, *t* = +3.61, *p*_BH_ = 3.4 × 10^-4^). At the individual level, more individuals than expected by chance had detectable Info×Unc[Rew] weights in only one context (151*/*1954 = 7.7% gain-only, *p <* 0.0001, and 131*/*1954 = 6.7% loss-only, *p* = 1.2 × 10^-4^, one-tailed binomial enrichment tests vs. the 4.75% expected if the two contexts’ significance patterns were independent) and a small but significant proportion had negative scaling in both contexts (8*/*1954 = 0.4% vs. 0.0625% expected under the same independence null; one-tailed binomial enrichment test, *p <* 0.0001; Supplementary Figure 3A and Methods). This heterogeneity, too, exceeded estimation noise (Cochran’s *Q* tests as in Figure 2C: Info×Unc[Rew] gain *Q*(1953) = 2978, *I*^2^ = 34%; loss *Q*(1953) = 2822, *I*^2^ = 31%; both *p <* 0.0001), and the per-participant gain-loss difference in this scaling was itself more variable than estimation noise alone would produce (*Q*(1953) = 2228, *I*^2^ = 12%, *p* = 1.2 × 10^-5^; Supplementary Figure 1D), indicating genuine individual variation not only in the strength of information– uncertainty scaling but in how it changes between contexts. Thus, overall, participants scale the value of information with reward uncertainty, but with substantial individual heterogeneity in both the strength of this scaling and its context dependence.

To more directly test the context dependence of information value scaling with reward uncertainty, we examined differences in individual GLM weights for gain vs loss. Averaging the individual weights, Info×Unc[Rew] scaling was significantly stronger in gain than in loss (+0.0177 ± 0.0057, *t*(1953) = +3.09, *p* = 0.0020, paired *t*-test on the gain-minus-loss difference; Figure 4B).

We next asked whether individual variation in this computation was related to psychopathology. Interestingly, just as high-AD individuals were more sensitive to monetary reward, they also scaled the value of information more strongly with reward uncertainty than did low-AD individuals; and just as high-CI individuals were less sensitive to monetary reward, they scaled information value with reward uncertainty less strongly—a parallel between reward sensitivity and information–uncertainty scaling that holds for both factors. In a trial-level cluster-robust logistic GLM of information choice with continuous risky × factor interactions (Figure 4C displays the model’s predictions for high- and low-factor groups; cluster-robust Wald tests, uncorrected; Methods), the AD interaction was significant in the loss context (*β* = +0.058 ± 0.023, *z* = 2.53, *p* = 0.012), with higher-AD individuals scaling information value more strongly with reward uncertainty; the reverse pattern held for CI, whose interaction was significant in the loss context (*β* = −0.055 ± 0.021, *z* = −2.68, *p* = 7.4 × 10^-3^), with higher-CI individuals scaling information value less strongly. The corresponding gain-context interactions were not significant for AD (*z* = −0.59, *p* = 0.56) or CI (*z* = −0.43, *p* = 0.67), and the SW interaction was not significant in either context (gain *z* = 1.12, *p* = 0.26; loss *z* = 0.85, *p* = 0.39); the gain–loss comparison of these factor effects is quantified by the population-model three-way terms below. When we instead jointly assessed economic and psychiatric covariates by fitting these factor effects in the population GLM (Info × Unc[Rew]×factor terms, cluster-robust Wald tests; Figure 4D), the AD effect was significant only in loss (Info × Unc[Rew]×AD in loss: *β* = +0.008, *p*_raw_ = 1.7 × 10^-2^, *p*_BH_ = 0.034; gain *β* = +0.0001, *p*_raw_ = 0.99)—although the corresponding attribute×context×factor term did not survive BH correction (Figure 4D), so this loss-specificity should be interpreted cautiously—whereas the CI effect was significant in loss (Info × Unc[Rew]×CI loss: *β* = −0.007, *p*_raw_ = 2.1 × 10^-2^, *p*_BH_ = 0.041) with a similar nominal effect in gain that did not survive BH correction (*β* = −0.007, *p*_raw_ = 3.8 × 10^-2^, *p*_BH_ = 0.071), and SW did not show significant effects (gain *β* = +0.008, *p*_raw_ = 0.062; loss *β* = +0.004, *p*_raw_ = 0.304).

Together, these results extend the previously described scaling of information value with reward uncertainty to loss decisions and show that its strength is context dependent, with stronger scaling in gain. The strength of this computation also varied with psychopathology: higher AD was associated with stronger information–uncertainty scaling, whereas higher CI was associated with weaker scaling, most clearly in loss. Notably, these opposing associations paralleled the corresponding increases and decreases in reward sensitivity associated with AD and CI, linking individual variation in a computation governing information value to broader variation in economic value sensitivity.

### 2.5 Patterns of psychopathology-related changes in economic preferences partially align with theory-driven predictions

The analyses above revealed systematic associations between psychopathology and multiple economic preferences across gain and loss. Measuring several economic attributes within the same task provides an important advantage for interpreting these associations in the interest of identifying computations that underlie decision-making and changes in psychopathology. A change in sensitivity to any single attribute could either reflect a specific alteration in how that attribute is valued, but it could also arise from changes in mental computations that affect decisions more broadly [7, 9, 11].

One possibility is that psychopathology alters a scalar parameter of economic sensitivity, potentially related to motivation or engagement, which acts by steepening or dampening the population-average preference profile, rather than by selectively altering particular attributes (Supplementary Figure 11A,B). Separate scalars could exist for gain and loss that allow context-dependent modulation of choice behavior. To formally test this account, we compared the effects fit by the unconstrained population GLM, which included separate effects of psychopathology factors in separate attributes, against an alternative constrained scaling model in which each psychopathology factor was allowed only to scale the population-average attribute weights up or down by a single scalar for each context (Supplementary Figure 11C,D; Methods).

Using this approach, we first examined the predictions made by the constrained model (Figure 5A). For easier comparison, we arbitrarily fixed the scaling of E[Rew] weights to match the unconstrained model. At a glance, the constrained model provided an account of how psychopathology factors would be expected to affect preferences for each attribute in gain and loss. All three psychopathology dimensions showed significant broad scaling in both contexts (Figure 5B). AD and SW steepened the population-average attribute weights (two-sided Wald *z* tests of scaling factor *a* = 1, all six *p <* 0.001; AD: *a*_gain_ = 1.082, *z* = +5.11, *a*_loss_ = 1.126, *z* = +6.09; SW: *a*_gain_ = 1.083, *z* = +4.99, *a*_loss_ = 1.083, *z* = +3.84), whereas CI dampened them (CI: *a*_gain_ = 0.845, *z* = −11.26; *a*_loss_ = 0.813, *z* = −10.70). Notably, the magnitude of this scaling depended on context. AD’s steepening and CI’s dampening were both significantly stronger in loss than in gain (Wald *z* tests of *a*_gain_ - *a*_loss_; AD: *z* = −2.50, *p* = 0.012; CI: *z* = +2.11, *p* = 0.035), whereas SW’s steepening did not differ significantly between contexts (*z* = 0.005, *p* = 0.996; maximum-likelihood counterpart in Supplementary Figure 12A).

**Figure 5.**
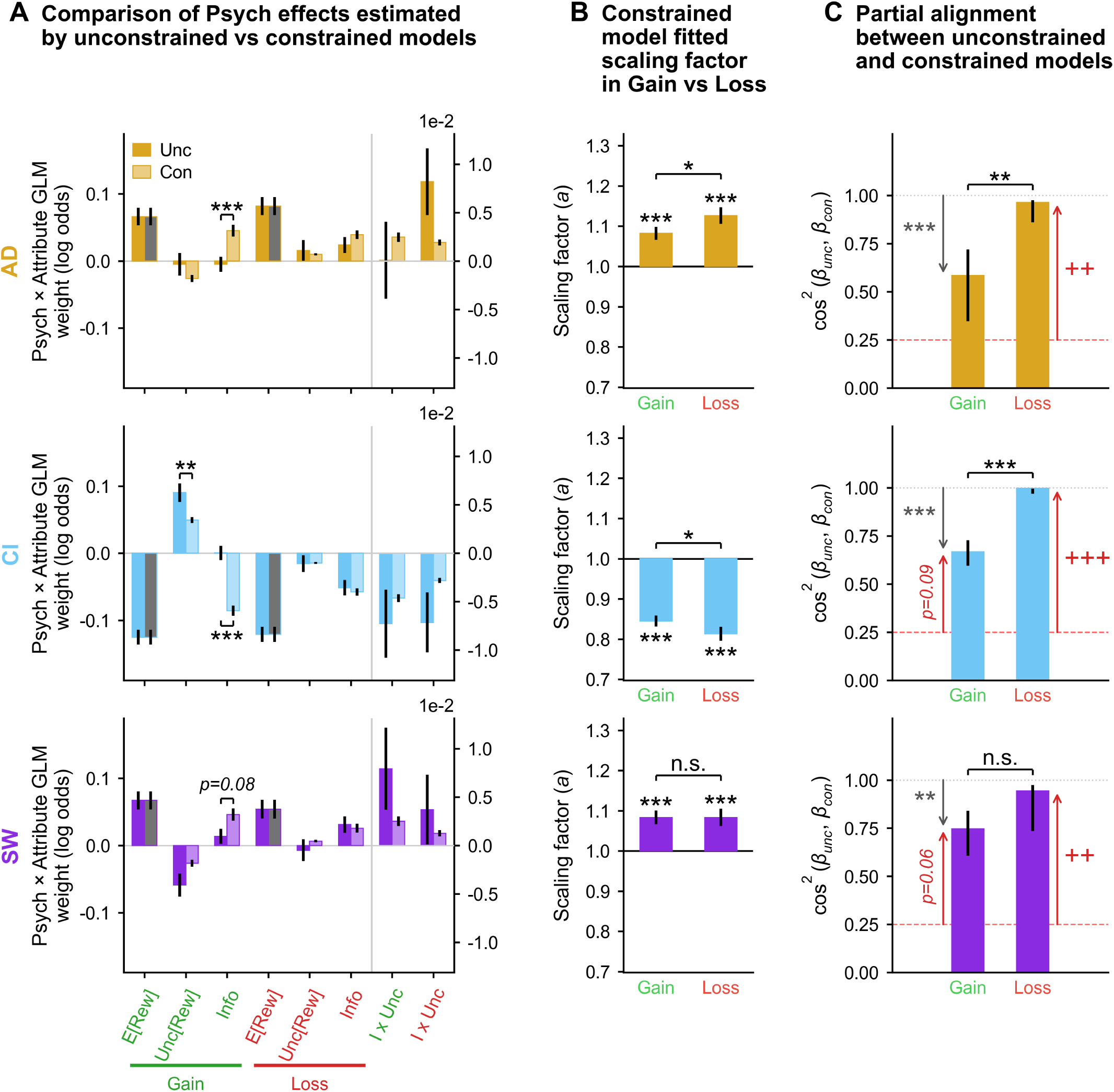
Testing a scaling account of psychopathology-related changes in economic preferences. We tested a “scaling” account in which each psychopathology factor broadly increases or decreases the overall strength with which the economic attributes influence choice, multiplying the population-average attribute weights by a single scalar per context (motivation, engagement, attention, and choice consistency are possible but untested interpretations of such scaling). The unconstrained comparison model fits free factor × attribute interactions in each context (8 per factor: 4 attributes × 2 contexts); the constrained model instead pins each context’s scalar to the factor’s own fitted E[Rew] interaction rather than fitting it freely (Methods). Both models use a shared base model (attribute main effects + context interactions, no psychopathology terms) as a fixed offset, with cluster-robust standard errors throughout (Methods). The figure is a transposed 3 × 3 grid: rows are the three psychopathology factors, named in the left margin in the factor colors (AD gold, CI light blue, SW violet), and columns are the three views **(A)**–**(C)** described below. Each cell carries its own y axis word and scale; columns **(B)** and **(C)** label their two bars Gain (green) and Loss (red) on every row, while column **(A)**’s attribute names, context underlines and Gain/Loss block labels are drawn on the bottom row only and apply to all three. **Supplementary Figure 11** walks through the scaling hypothesis and the alignment statistic schematically; the complementary “context-change” account is tested in **Supplementary Figures 13 and 14. (A)** Comparison of psychopathology effects estimated by the unconstrained vs. the constrained model: for each factor, the eight attributecontext interaction coefficients drawn as pairs of bars—unconstrained (dark, legend *Unc*) against the pinned-constrained prediction (light, *Con*)—with cluster-robust SE bars, on the y axis *Psych* × *Attribute GLM weight (log odds)*. Slot order is E[Rew], Unc[Rew], Info in gain, then the same three in loss, then, past a vertical divider and drawn against the righthand axis at ×10^-2^ because they are an order of magnitude smaller, the two Info × Unc[Rew] slots (*I* × *Unc*) in gain and loss. The two E[Rew] constrained bars are drawn in gray, not in the factor’s light tier, because the scalar is solved so that the constrained model reproduces the unconstrained E[Rew] weight exactly: those bars are the pin itself, are equal to their neighbor by construction, and carry no test. Brackets and stars mark attribute-level departures from the scaling prediction (exact linear contrast under the population-model covariance, BH-corrected within factor over the 8 attribute-context effects per factor, of which the two pinned E[Rew] comparisons enter at *p* ≡ 1): three departures are significant, all in the gain context—AD’s Info comparison (*p*_BH_ = 3.4 × 10^-4^) and CI’s Unc[Rew] and Info comparisons (*p*_BH_ = 8.2 × 10^-3^ and *<*0.0001)—with SW’s Info comparison marginal and drawn with its literal value (*p*_BH_ = 0.08) and no other significant deviations. See **Supplementary Figure 12B** for the corresponding maximum-likelihood comparison (significant for CI’s E[Rew], Unc[Rew], and Info in gain). **(B)** Constrained-model fitted scaling factor in gain vs. loss: the scalar pinned to each factor’s own fitted E[Rew] interaction in that context (*a*_ctx_ = 1 + *β*_E[Rew],ctx_*/µ*_E[Rew],ctx_), drawn as a bar from a baseline of 1 on the y axis *Scaling factor (a)*. Per-bar stars are two-sided *z* tests of *a* = 1 and sit outside the error bar on the side away from the baseline. AD and SW significantly steepened the population-average attribute weights in both contexts (AD *a*_gain_ = 1.082, *a*_loss_ = 1.126; SW *a*_gain_ = 1.083, *a*_loss_ = 1.083), whereas CI dampened them (*a*_gain_ = 0.845, *a*_loss_ = 0.813); all six *p <* 0.001 (*z* = 5.11, 6.09, 4.99, 3.84, −11.26, −10.70). The bracket above each cell tests the gain–loss difference in scaling using the exact cross-context covariance of the two E[Rew] coefficients: scaling departs further from 1 in loss for AD (*a*_gain_ - *a*_loss_ = −0.044, *p* = 0.012) and for CI (+0.031, *p* = 0.035) but not for SW (+0.0001, *p* = 0.996, printed n.s.). Error bars show cluster-robust SE; *, **, *** indicate *p <* 0.05, 0.01, 0.001. See **Supplementary Figure 12A** for the corresponding maximum-likelihood version. **(C)** Partial alignment between the unconstrained and constrained models: the squared cosine between each context’s unconstrained 4-attribute coefficient vector and the population-average pattern for that context, on the y axis cos^2^ (*β*_unc_*, β*_con_). The gray dotted line at 1.00 marks perfect alignment; the red dashed line at 0.25 marks the expected value of cos^2^ under chance, not a significance threshold. Error bars are the central 68% percentile interval of a 10,000-draw bootstrap over the full coefficient covariance, drawn as that interval rather than anchored on the bar (near the ceiling the bootstrap distribution sits below the observed value, because noise can only rotate a vector away from alignment). A gray downward arrow with gray asterisks marks a significant departure from perfect alignment (cluster-robust Wald test, *χ*^2^(3)); a red upward arrow with crosses marks significantly above-chance alignment under the calibrated effect-direction null (+/++/+ + + at *p <.*05*/.*01*/.*001; Methods), and a chance *p* in [.05,.1) is drawn as a red arrow labeled with its literal *p*; an absent arrow denotes a non-significant test. Brackets show the gain-versus-loss bootstrap comparison. Loss-context alignment was statistically indistinguishable from perfect for all three factors (AD = 0.964, *p* = 0.12; CI = 0.997, *p* = 0.51; SW = 0.944, *p* = 0.69) and significantly above chance (AD *p* = 0.003; CI *p <* 0.0001; SW *p* = 0.006); gain-context alignment departed significantly from perfect for all three (AD = 0.584, *p* = 3.6 × 10^-4^; CI = 0.667, *p <* 0.0001; SW = 0.747, *p* = 2.5 × 10^-3^) and none individually exceeded chance (AD *p* = 0.14, no arrow; CI *p* = 0.09; SW *p* = 0.06). The gain-versus-loss difference was itself significant for AD and CI but not SW (AD *p* = 0.009; CI *p <* 0.0001; SW *p* = 0.35). See **Supplementary Figure 12C** for the corresponding joint (8-D) maximum-likelihood statistic (AD = 0.81, *p* = 0.0064; CI = 0.80, *p* = 0.0076; SW = 0.80, *p* = 0.0081; all ab ove chance). Takeaway: pinned scaling factors capture most of each factor’s effect pattern — AD and SW globally steepen and CI globally dampens economic preferences, more strongly in loss for AD and CI — with alignment statistically indistinguishable from perfect scaling in loss for all three factors but significantly departing from it in gain for all three; these gain-context residuals, most extensive for CI, indicate structure beyond uniform scaling and motivate the data-driven analysis in **Figure 6**.

**Figure 6.**
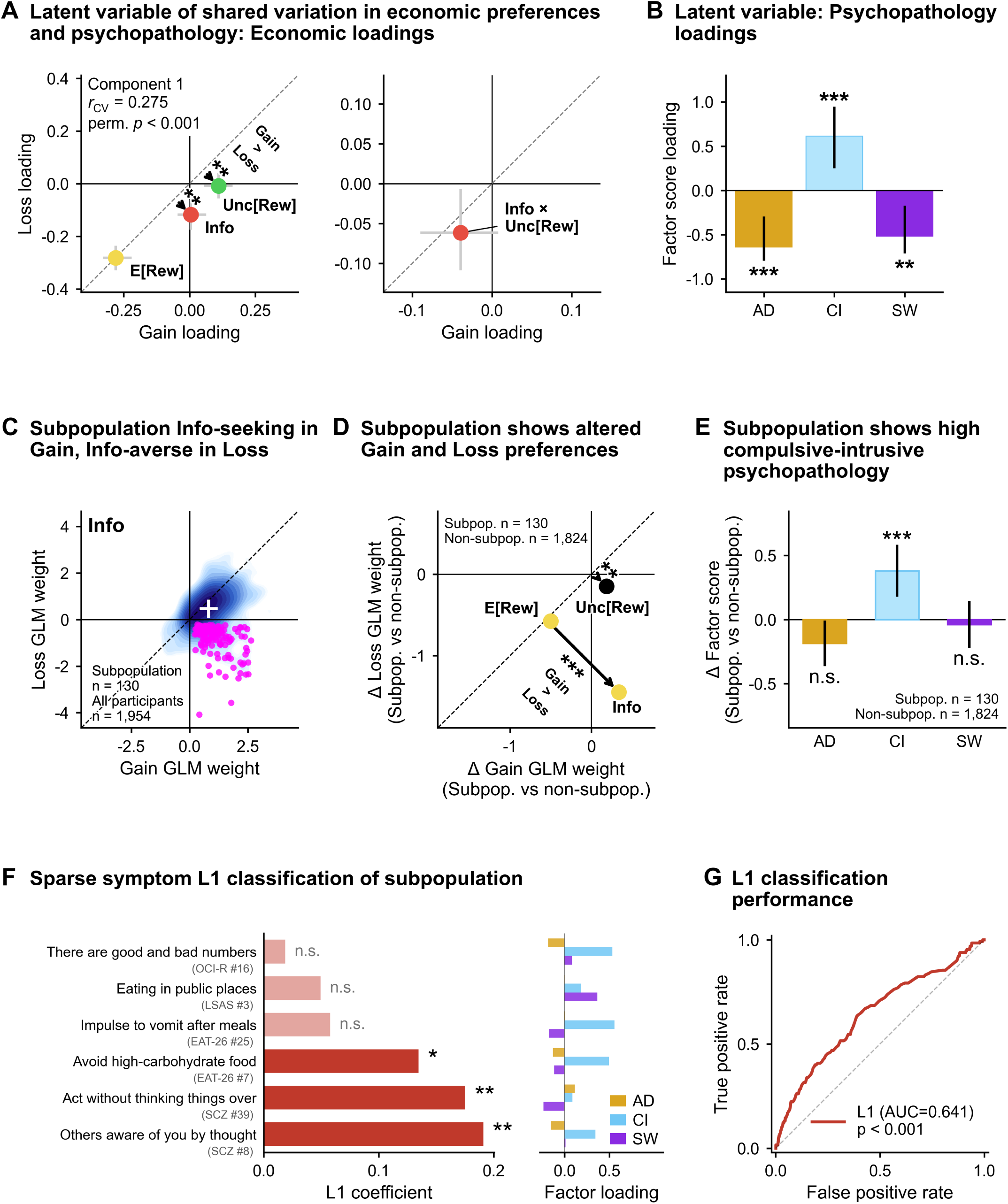
A latent dimension of shared economic-psychopathological variation converges with an independently identified information-reflecting subpopulation high in compulsive-intrusive psychopathology. Multivariate (partial least squares, PLS) analysis of economic preferences and psychopathology identifies a leading latent dimension of shared variation that converges, without theoretical guidance, on the same compulsive-intrusive factor highlighted by prior analyses—pairing weaker reward and information preferences (especially information-avoidance in loss) with elevated CI. A subgroup of n=130 information-reflecting individuals, identified directly from the choice data and first apparent in the **Figure 2C** Info distributions, independently of and prior to the PLS solution, matches that profile, is selectively enriched on CI, and is predictable above chance from a sparse set of 6 psychiatric symptom items. **(A)** Latent variable of shared variation: Component 1 economic loadings from a PLS regression predicting individual economic preference GLM weights (E[Rew], Unc[Rew], Info and Info × Unc[Rew] in gain and loss; 8 responses) from psychopathology factor scores (AD, CI, SW) for n=1954 participants. Two scatters share the axes *Gain loading* (x) and *Loss loading* (y) with a gray dashed identity line; the left scatter holds E[Rew], Unc[Rew] and Info, and the right one holds Info × Unc[Rew] alone, at its own finer scale. Gray crosses are 95% bias-corrected and accelerated (BCa) bootstrap intervals (2,000 resamples) and point color marks a loading significantly different from zero in gain (green), loss (red), both (yellow) or neither (black), by exact two-sided bootstrap p-values BH-corrected over the 8 economic loadings. Component 1 loads strongly negative on reward in both contexts (E[Rew]_gain_ −0.281, 95% CI [−0.329, −0.220]; E[Rew]_loss_ −0.282, [−0.329, −0.236]), negative on information in loss (Info_loss_ −0.118, [−0.175, −0.062]; Info_gain_ +0.005, [−0.046, +0.063], n.s.), positive on reward uncertainty in gain (Unc[Rew]_gain_ +0.110, [+0.054, +0.163]; Unc[Rew]_loss_ −0.009, [−0.056, +0.041], n.s.), and negative on Info × Unc[Rew] scaling in loss only (−0.061, [−0.109, −0.007], *p*_BH_ = 0.038; the gain loading is negative but not significant, −0.039, [−0.090, +0.008], *p*_BH_ = 0.167). Arrows drawn from the identity line to a point, with their star tier and one bold *Gain > Loss* note, mark loadings whose gain and loss values differ significantly from each other, tested by a paired within-replicate bootstrap contrast (gain minus loss, computed from the same 2,000 PLS bootstrap replicates used for the loading CIs; exact two-sided *p* floored at 1*/*2000; BH-FDR over the 4 regressor contrasts as their own family; Methods): Unc[Rew] (+0.119, *p*_BH_ = 0.0020) and Info (+0.123, *p*_BH_ = 0.0020) differ, while E[Rew] (*p*_BH_ = 0.980) and Info × Unc[Rew] (*p*_BH_ = 0.656) do not; this contrast is a distinct test from the vs.-zero loading significance above and from the analogous arrows in **(D)**. We adopted a one-component model as primary: cross-validated MSE (5-fold × 10-shuffle CV; mean MSE = 1.008 → 0.991 → 0.990 → 0.991 from 0 to 3 components, SE ≈ 0.0005) was minimized at two components but by a negligible margin (ΔMSE ≈ 0.0008), an elbow-point criterion favored one dominant component, and the second component generalized only weakly to held-out participants (Methods; component-selection diagnostics in **Supplementary Figure 16**). The overall one-component regression was significant (in-sample *R*^2^ = 0.019, equal-weighted across the eight responses; 1000-permutation *p* = 0.001). The block at the upper left of the left scatter gives the held-out latent-score correlation of Component 1 — *r*_CV_ = 0.275: the correlation, across the held-out participants of the same 5-fold × 10-shuffle cross-validation, between the psychopathology-side and preference-side Component-1 scores computed with weights learned on the training folds alone (in-sample counterpart *r* = 0.289), with its two-sided permutation *p <* 0.001 from 1,000 permutations of the full cross-validation pipeline. This statistic characterizes the robustness of the latent association itself; it is distinct from the model’s prediction of the eight response variables (cross-validated *R*^2^ = 0.0156, annotated in **Supplementary Figure 16A**) and from the in-sample *R*^2^ = 0.019 quoted in the preceding sentence. **(B)** Latent variable, psychopathology loadings: bars of the Component 1 X-loading of each psychopathology factor, y axis *Factor score loading*, with 95% BCa bootstrap CIs (2,000 resamples) and stars from exact two-sided bootstrap p-values BH-corrected over the three factor loadings. CI loads strongly positive (+0.614, [+0.249, +0.946]) while AD and SW load negative (AD −0.639, [−0.795, -0.297]; SW −0.518, [−0.713, −0.173], *p*_BH_ = 0.003; AD and CI *p*_BH_ *<* 0.001), forming a CI-vs-(AD+SW) contrast. Combined with the economic loadings in **(A)**, the dominant latent dimension pairs a compulsive-intrusive pattern of economic preferences (weaker reward and information preferences, less risk-aversion in gain, weaker scaling of information value with reward uncertainty) with an elevated CI-and-not-AD/SW psychiatric profile. **(C)** Subpopulation info-seeking in gain, info-averse in loss: individual Info GLM weights in gain (x-axis) vs. loss (y-axis), with the whole population (n=1954) drawn as a filled log-density contour and the n=130 information-reflecting individuals (“info-reflecting subpopulation”) plotted in magenta; the white crosshair is the population mean ± SEM (+0.808 ± 0.014 gain, +0.470 ± 0.015 loss), the dashed line is unity and the solid lines mark zero. The info-reflecting subpopulation was defined operationally from the choice data alone, as those individuals whose individual-level GLM Info weights were both significantly positive in gain and significantly negative in loss (*p <* 0.05 in both). Among the 1,374 individuals with individually significant Info weights in both contexts, 130 showed this information-reflecting pattern (gain *>* 0, loss *<* 0) while zero showed the opposite (gain *<* 0, loss *>* 0; binomial test, *p <* 0.0001), so the sign asymmetry is strongly unidirectional rather than a symmetric tail artifact. **(D)** Subpopulation shows altered gain and loss preferences: the difference in population-average individual GLM weights (info-reflecting subpopulation minus rest of population) for E[Rew], Unc[Rew] and Info, plotted as the gain-context difference (x-axis Δ Gain GLM weight) vs. the loss-context difference (y-axis Δ Loss GLM weight); both axes are in log odds, and loss-context E[Rew] weights are sign-flipped before differencing so that positive indicates reward-seeking. The dashed line marks equality of the two differences. Point color is the per-context info-reflecting subpopulation-vs-rest comparison (two-tailed Welch’s *t*-test), BH–FDR corrected over the 16 tests in this panel (4 displayed regressors × 2 contexts × 2 test types, within-subgroup and subgroup-vs-rest). Arrows from the identity line, their stars and the bold *Gain > Loss* note mark attributes whose gain- and loss-context differences themselves differ, by a separate two-tailed Welch’s *t*-test on the gain-minus-loss contrast; these three arrow tests are reported uncorrected: Unc[Rew] (*p* = 0.0022) and Info (*p <* 0.0001) differ significantly between contexts, but E[Rew] does not (*p* = 0.21, no arrow). *, **, *** indicate *p <* 0.05, 0.01, 0.001. The underlying per-context comparisons: the info-reflecting subpopulation had weaker reward sensitivity in both contexts (E[Rew] gain, in +0.86 ± 0.07 vs. out +1.36 ± 0.02, *t* = −6.80, *p*_BH_ *<* 0.0001; E[Rew] loss, in +0.57 ± 0.06 vs. out +1.15 ± 0.02, *t* = −8.54, *p*_BH_ *<* 0.0001), the defining sharp negative shift in Info in loss (in −0.88 ± 0.06 vs. out +0.57 ± 0.01, *t* = −23.77, *p*_BH_ *<* 0.0001) and stronger information-seeking in gain (in +1.12 ± 0.06 vs. out +0.79 ± 0.01, *t* = +5.72, *p*_BH_ *<* 0.0001). Group differences in risk preference and in Info × Unc[Rew] scaling did not survive the 16-test correction (Unc[Rew] gain, in −0.44 ± 0.08 vs. out −0.62 ± 0.03, *t* = +2.11, *p*_raw_ = 0.036, *p*_BH_ = 0.053; Info × Unc[Rew] loss, in −0.004 ± 0.017 vs. out +0.018 ± 0.005, *t* = −1.21, *p*_raw_ = 0.23). The info-reflecting subpopulation’s economic profile thus closely mirrors the Component 1 loadings in **(A)**. **(E)** Subpopulation shows high compulsive-intrusive psychopathology: bars of the difference in mean AD, CI and SW factor score between the n=130 info-reflecting subpopulation members and the n=1,824 remaining participants (y axis Δ *Factor score*, subtitled *Subpop. vs non-subpop.*; bar colors AD goldenrod, CI baby blue, SW blue violet), with both group sizes printed in the panel. Whiskers are the 95% confidence interval of the difference—not SE, so they are roughly twice as long as those of the equivalent one-sample display. Factor scores are the frozen full-sample z-scores used throughout and subgroup membership is unchanged from **(C)**; the contrast is therefore between the subgroup and the rest of the sample on the same scores. Stars are two-tailed Welch two-sample *t*-tests of subpopulation against non-subpopulation, BH-corrected across the three factors, and *n.s.* is printed where the test does not clear 0.05. CI is substantially elevated (+0.380, 95% CI [+0.178, +0.583]; *t*(143.1) = +3.71, *p*_raw_ = 2.9 × 10^-4^, *p*_BH_ = 8.7 × 10^-4^), AD is nominally lower but does not clear this panel’s BH criterion (−0.186, [−0.364, −0.007]; *t*(148.2) = −2.06, *p*_raw_ = 0.041, *p*_BH_ = 0.062, unstarred) and SW does not differ between the groups (−0.038, [−0.222, +0.146]; *t*(146.9) = −0.41, *p* = 0.69) — so the info-reflecting subpopulation is selectively and positively associated with compulsive-intrusive psychopathology, not with a general internalizing profile. **(F)** The kind of symptom content the classifier draws on. This panel is illustrative rather than a definitive symptom list: it shows which individual items the sparse model retained and how strongly each is supported, not the full set of symptoms on which the subgroup differs. Left: signed L1 coefficients of an L1-regularized logistic classifier predicting info-reflecting subpopulation membership from the 209 individual psychiatric symptom questionnaire items, fit with nested cross-validation (10-fold outer × 5-fold inner; class-balanced loss; L1 strength selected on the inner folds); one horizontal bar per item with a nonzero penalty-selected weight, each labeled with a short form of the item and, beneath it, its instrument and item number. Stars give each item’s scale-matched permutation *p* under maximumstatistic family-wise correction (*, **, *** for *p <* 0.05, 0.01, 0.001; Methods), and items that do not reach *p <* 0.05 are drawn in a lighter tint and labeled *n.s.* — the tint reinforces the label rather than replacing it. Right: the same six items’ loadings on AD (goldenrod), CI (baby blue) and SW (blue violet). The L1 solution selected only 6 of 209 items, and all six coefficients are positive: “Others aware of you by thought” (SCZ #8, 0.191), “Act without thinking things over” (SCZ #39, 0.175 — this item is reverse-keyed, so the drawn label states the scored direction rather than the question’s literal wording), “Avoid high-carbohydrate food” (EAT-26 #7, 0.135), “Impulse to vomit after meals” (EAT-26 #25, 0.058), “Eating in public places” (LSAS #3, 0.049) and “There are good and bad numbers” (OCI-R #16, 0.019). All six have positive CI factor loadings, four of them substantial (+0.56, +0.53, +0.50, +0.34), with the remaining two—a schizotypy and a social-anxiety item—loading only weakly on CI (+0.09, +0.19). The sparsity of this solution shows that a small item combination carries the predictive signal; because L1 retains only a minimal predictive subset among correlated items, it does not imply that only these six symptoms are elevated in the info-reflecting subpopulation. The identity of the six is accordingly weaker evidence than the model-level result, and the six are not equally well supported: against a scale-matched permutation null the three largest coefficients are individually significant under family-wise error control (SCZ #8 and #39, both *p* = 0.003; EAT-26 #7, *p* = 0.019), while the three smallest are not (*p* = 0.35, 0.45 and 0.81; Methods). **(G)** L1 logistic classifier ROC: receiver operating characteristic curve for the classifier in **(F)**, evaluated on held-out outer-fold predictions (n=1954; 130 info-reflecting subpopulation, 1824 non-info-reflecting subpopulation), with the dashed diagonal marking chance and the in-panel legend giving the area under the curve and its permutation p-value. The classifier exceeded chance in held-out participants (AUC = 0.641, 1000-iteration model-level label-permutation test with a full nested-CV refit per permutation, *p <* 0.001; balanced accuracy 0.596). Sensitivity 0.546 and specificity 0.646 (held-out TP = 71, FN = 59, TN = 1178, FP = 646) describe the single operating point at a 0.5 threshold and are reported for completeness only: neither exceeds its own nested-CV label-permutation null (*p* = 0.38 and *p* = 0.25; a class-balanced classifier on a 93%-negative sample attains specificity near 0.6 under shuffled labels), so they should not be read as separate evidence of performance. Together these indicate that info-reflecting subpopulation membership is predictable from individual symptom items alone, not only from the aggregate CI factor. Takeaway: a leading PLS latent dimension captures co-variation between economic preferences and psychopathology and pairs a compulsive-intrusive economic-preference pattern with a CI-vs-(AD+SW) psychiatric contrast; an independently defined n=130 information-reflecting subpopulation—identified directly from the choice data—matches this dimension’s profile, is selectively elevated on CI, and has membership predictable above chance from a sparse 6-item combination of individual psychiatric symptoms.

We next asked how well this broad scaling account captured the full multivariate pattern. Overall, while the constrained and unconstrained models made similar predictions (Figure 5A), there were a few significant differences in how psychopathology affected risk and information preferences (two-sided Wald *z* tests, BH-corrected). While the apparently steepened preferences in AD would predict a substantial increase in information preference across contexts, AD showed essentially no unconstrained modulation of Info in gain, significantly below the constrained prediction (*z* = −4.09, *p*_BH_ = 3.4 × 10^-4^). Conversely, CI’s broad dampening predicted a weakened influence of every attribute, yet in gain both its risk and information effects sat significantly above that prediction: risk aversion was reduced more than dampening alone would produce (Unc[Rew]: *z* = +3.08, *p*_BH_ = 8.2 × 10^-3^), and information preference was largely spared rather than dampened (Info: *z* = +7.24, *p*_BH_ *<* 0.0001). SW showed the same qualitative shortfall in gain-context information preference as AD, though only marginally (Info: *z* = −2.59, *p*_BH_ = 0.076), and there were no other significant deviations. The corresponding maximum-likelihood comparison (Supplementary Figure 12B) showed the same qualitative pattern, with all three of CI’s gain-context comparisons (E[Rew], Unc[Rew], and Info) significant.

Quantifying alignment directly and separately by context revealed a clear asymmetry (Figure 5C). In loss, alignment was statistically indistinguishable from perfect scaling for all three factors (deviation-from-perfect cluster-robust Wald tests: AD cos^2^ = 0.964, *χ*^2^(3) = 5.88, *p* = 0.118; CI cos^2^ = 0.997, *χ*^2^(3) = 2.32, *p* = 0.509; SW cos^2^ = 0.944, *χ*^2^(3) = 1.45, *p* = 0.694) and significantly above the chance expected value of 1*/*4 under the calibrated directional null (Methods; AD *p* = 0.003; CI *p <* 0.0001; SW *p* = 0.006). In gain, by contrast, alignment departed from perfect scaling for all three factors (AD cos^2^ = 0.584, *χ*^2^(3) = 18.43, *p* = 0.0004; CI cos^2^ = 0.667, *χ*^2^(3) = 68.04, *p <* 0.0001; SW cos^2^ = 0.747, *χ*^2^(3) = 14.35, *p* = 0.0025) and did not individually exceed chance (AD *p* = 0.140; CI *p* = 0.092; SW *p* = 0.061). A gain-versus-loss comparison of these per-context cos^2^ values (bootstrap percentile test; Methods) confirmed this asymmetry was itself significant for AD and CI but not SW (AD *p* = 0.009; CI *p <* 0.0001; SW *p* = 0.349). The corresponding joint (8-D) alignment statistic remained high for all three factors (Supplementary Figure 12C; cos^2^: AD = 0.81; CI = 0.80; SW = 0.80; calibrated above-chance *p*: AD = 0.0064; CI = 0.0076; SW = 0.0081; joint cluster-robust Wald test of perfect alignment (*χ*^2^(6)) rejected for all three: AD *W* = 22.6, *p* = 9.2 × 10^-4^; CI *W* = 72.2, *p <* 0.0001; SW *W* = 15.9, *p* = 0.015), and scoring alignment instead as the orthogonal projection onto the two-scalar hypothesis space gave the same picture (AD = 0.82; CI = 0.81; SW = 0.81; Supplementary Figure 12D). Thus, broad scaling captured a substantial portion of the psychopathology-related pattern, particularly in loss, but left additional attribute-specific structure, most prominently for CI.

Alternatively, psychopathology could alter the representation of outcome context or reference point, causing preferences to shift in a more gain-like or loss-like direction. We therefore hypothesized that psychopathology-related changes might also resemble the average multivariate change in preferences produced by gain versus loss framing itself (Supplementary Figures 13 and 14). We formalized this context-change hypothesis by fitting a constrained model in which each psychopathology factor’s attribute interactions were replaced by scalar interactions with *δ*, with separate gain and loss coefficients allowing gain-directed or loss-directed effects in either context (Supplementary Figure 14A; Methods). This context-change account explained substantially less of the observed variation (Supplementary Figure 14B-E), partially because the context-change and scaling axes themselves overlap. Thus, although AD and CI showed clear evidence that psychopathology-related changes were stronger under loss framing, those effects were not well characterized as shifts along the particular multivariate direction that normally distinguishes gain from loss.

We also asked whether the observed psychopathology effects on reward and risk could be captured within a classic behavioral economics framework. To this end, we fit choice behavior using a cumulative prospect-theory (CPT) model in which the magnitudes of probabilistic reward outcomes were nonlinearly transformed according to a power parameter *α* (Methods). Probability weighting was fixed at its population estimate for these fits, while information-related effects were retained from the corresponding GLM. At the population level, *α* was below 1 in both gain (0.970 ± 0.005) and loss (0.908 ± 0.007; cluster-robust Wald tests vs. 1, both *p <* 0.0001), consistent with the canonical pattern of concavity over gains and convexity over losses that contributes to risk aversion for gains and risk seeking for losses [6, 33] (Supplementary Figure 15C). Across individuals, fitted *α* was related to the linear E[Rew] and Unc[Rew] weights in directions broadly consistent with this interpretation (multivariate regression over the *n* = 1642 participants whose CPT fits converged in both contexts; Supplementary Figure 15B): E[Rew] was positively associated with *α* in both contexts (*β* = +0.225 ± 0.004, *t*(1639) = +51.9 in gain; +0.288 ± 0.005, *t* = +58.8 in loss; both *p <* 0.0001), whereas Unc[Rew] was positively associated with *α* in gain and negatively associated in loss (+0.076 ± 0.002, *t* = +30.9; −0.085 ± 0.003, *t* = −29.5; both *p <* 0.0001). Forward simulations across theoretical values of *α* reproduced the model-implied relationships directly (Supplementary Figure 15A; Methods). In the range of outcome magnitudes and reward distributions used in our task, increasing *α* strongly increased the model-implied E[Rew] choice weight in both contexts (*dβ/dα* ≈ +2.0 in gain and +1.7 in loss), while leaving the model-implied Unc[Rew] weight nearly unchanged (derivatives 14–53-fold smaller) and, if anything, directed at the fitted probability weighting toward less risk aversion in gain and less risk seeking in loss (Methods). Thus, within the range relevant to this task, magnitude curvature primarily acts as a reward-sensitivity parameter and has comparatively little influence on the linear uncertainty weight.

We then asked whether psychopathology was associated with variation in *α* by allowing the parameter to vary with each psychopathology factor in the population CPT model (Methods). AD and SW were associated with higher *α* in both contexts, whereas CI was associated with lower *α* (AD: gain +0.026 ± 0.005, loss +0.043 ± 0.008; SW: gain +0.023 ± 0.005, loss +0.033 ± 0.008; CI: gain −0.047 ± 0.005, loss −0.074 ± 0.008; all six *p <* 0.0001; Supplementary Figure 15D), closely paralleling their respective increases and decreases in E[Rew] sensitivity. Thus, variation in prospect-theoretic magnitude distortion provides one plausible parameterization of the broad psychopathology-related differences in reward sensitivity. However, it does not account for the full pattern of risk-related effects. AD showed no detectable change in Unc[Rew], while CI and SW showed opposing, gain-specific alterations in uncertainty preference. Because the forward simulation shows that variation in *α* leaves the model-implied Unc[Rew] weight nearly unchanged (Supplementary Figure 15A), these risk-preference alterations cannot be attributed to the same differences in magnitude distortion; indeed, to the extent that *α* predicts any change in uncertainty preference at all, the predicted direction is opposite to the observed effects— higher *α*, as observed with SW, would if anything shift choices toward less risk aversion in gain, whereas SW showed greater risk aversion, and lower *α*, as observed with CI, predicts the converse, whereas CI showed reduced risk aversion. Prospect theory also does not specify the observed changes in non-instrumental information valuation.

Together, these analyses show that psychopathology-related changes in economic preferences contain a substantial broad component: AD and SW generally increased, whereas CI decreased, the influence of economic attributes on choice, with this scaling stronger under loss framing for AD and CI. This structure was better characterized by context-specific scaling than by shifts along the ordinary gain-to-loss preference axis, and its reward-sensitivity component could also be expressed as variation in prospect-theoretic magnitude curvature. However, systematic attribute- and context-specific deviations—particularly in risk and information preferences—show that these low-dimensional accounts do not fully explain the observed multivariate pattern.

### 2.6 A latent dimension highlights a compulsive-intrusive economic profile and motivates characterization of an information-reflecting subgroup

The analyses above characterized psychopathology associations attribute by attribute and tested whether their multivariate structure could be summarized by several theory-driven computational accounts. While these accounts revealed some overlap with our observed results, there were key differences, especially in risk and information preferences. We hypothesized that a data-driven dimensional approach could summarize these differences in how psychopathology is related to economic preferences. To this end, we used partial least squares (PLS) regression, a technique that identifies latent components that maximize covariance between two sets of variables, in order to summarize the shared variability in individual GLM weights for E[Rew], Unc[Rew], Info, and Info×Unc[Rew] in gain and loss, and AD, CI, and SW factor scores.

PLS regression furnished one component, which we examined as a latent dimension linking gain-loss decision-making and psychopathology (Methods). This component exhibited substantial cross-validated correlation between economic and psychopathology variables (*r*_CV_ = 0.275; two-sided permutation *p <* 0.001; Figure 6A) and the overall PLS regression was significant (in-sample *R*^2^ = 0.019, permutation *p* = 0.001; cross-validated *R*^2^ = 0.016; Methods). This latent dimension captured a pattern reminiscent of the attribute-level associations aligned with CI psychopathology and in opposition to AD and SW psychopathology. (Figure 6A). On the economic side, it had significantly negative loadings on reward in both contexts (E[Rew]_gain_ −0.281, 95% bootstrap CI [−0.329, −0.220]; E[Rew]_loss_ −0.282, [−0.329, −0.236]), a significantly negative loading on information in loss (Info_loss_ −0.118, [−0.175, −0.062]), a significantly positive loading on reward uncertainty in gain (Unc[Rew]_gain_ +0.110, [+0.054, +0.163]), and a significantly negative loading on Info×Unc[Rew] scaling in loss ((Info × Unc[Rew])_loss_ −0.061, [−0.109, −0.007], *p*_BH_ = 0.038; the corresponding gain loading was negative but not significant, −0.039, [−0.090, +0.008], *p*_BH_ = 0.167). On the psychopathology side, the component contrasted CI (X-loading +0.61, 95% bootstrap CI [+0.25, +0.95]) with AD (−0.64, [−0.79, −0.30]) and SW (−0.52, [−0.71, −0.17], *p*_BH_ = 0.003), indicating that this dimension distinguished CI from the other two factors (Figure 6B; loading significance by exact two-sided bootstrap *p*, BH-corrected within block). Thus, the PLS analysis highlighted a CI-associated profile combining weaker reward sensitivity across contexts with context-specific alterations in uncertainty and information preference.

We next asked whether individuals showing an especially strong expression of one of the behavioral features highlighted by this component might define a more interpretable subgroup. Earlier analyses of the behavioral distributions had already identified individuals who were information-seeking in gain but information-averse in loss (Figure 2C), a pattern resembling the component’s negative loading on loss-context information preference. We therefore asked whether these individuals also exhibited the broader economic profile summarized by the latent component and whether they were enriched for CI psychopathology. We refer to this sign reversal as information-reflection, by direct analogy to risk-reflection: just as risk attitudes can reverse sign between gain and loss, these individuals’ valuation of non-instrumental information reversed sign across contexts. To our knowledge, though previous work has examined information-seeking for appetitive and aversive outcomes [21, 22, 43], such a reversal of information preference has not been described before. We operationally defined an information-reflecting subpopulation as individuals whose individual-level GLM weights were both significantly positive for Info in gain and significantly negative for Info in loss (individual GLM, *p <* 0.05 for both; Figure 6C; *n* = 130 of 1954). This subgroup was substantially more abundant than the 0.0625% expected if significance in the two contexts were independent (6.7%; binomial test *p <* 0.0001; Supplementary Figures 1D and 3A). The directional asymmetry was striking: among the 1,374 individuals with individually significant Info weights in both contexts (*p <* 0.05 for each), 130 exhibited the information-reflecting pattern (Info gain *>* 0, Info loss *<* 0) while zero exhibited the opposite pattern (Info gain *<* 0, Info loss *>* 0; exact binomial sign test of 130 vs. 0 against an even split, *p <* 0.0001).

Examining other economic preferences relative to the rest of the population (Figure 6D; Welch’s *t*-tests, BH-corrected), information-reflecting individuals had significantly weaker reward-seeking in both contexts (E[Rew] gain: in +0.86 ± 0.07 vs. out +1.36 ± 0.02, *t*(149.2) = −6.80, *p*_BH_ *<* 0.0001; E[Rew] loss: in +0.57 ± 0.06 vs. out +1.15 ± 0.02, *t*(152.9) = −8.54, *p*_BH_ *<* 0.0001, the subgroup’s weaker positive weight again indicating weaker reward sensitivity; the subgroup’s own six weights are shown in Supplementary Figure 17B). The defining Info-loss difference was very large (Info loss: in −0.88±0.06 vs. out +0.57 ± 0.01, *t*(140.7) = −23.77, *p*_BH_ *<* 0.0001), and the subgroup was also significantly more information-seeking in gain (Info gain: in +1.12 ± 0.06 vs. out +0.79 ± 0.01, *t*(145.3) = +5.72, *p*_BH_ *<* 0.0001), consistent with an amplified information-reflection pattern rather than general indifference to information. Group differences in risk preference and in Info×Unc[Rew] scaling did not survive the figure’s BH correction (Unc[Rew] gain: in −0.44 ± 0.08 vs. out −0.62 ± 0.03, *t*(176.8) = +2.11, *p*_raw_ = 0.036, *p*_BH_ = 0.053; Info×Unc[Rew] loss: *t*(149.2) = −1.21, *p*_raw_ = 0.23). Psychopathologically, information-reflecting individuals scored significantly higher on CI than the rest of the population (difference = +0.380, 95% CI [+0.178, +0.583], Welch’s *t*(143.1) = +3.71, *p*_raw_ = 2.9 × 10^-4^, *p*_BH_ = 8.7 × 10^-4^), nominally lower on AD (−0.186, [−0.364, −0.007], *t*(148.2) = −2.06, raw *p* = 4.1 × 10^-2^, not significant after BH correction across the three factors, *p*_BH_ = 0.062), and did not differ from the rest of the population on SW (−0.038, [−0.222, +0.146], *t*(146.9) = −0.41, *p* = 0.69) (Figure 6E). Thus, a subgroup defined solely by a strong context reversal in information preference also showed reduced reward sensitivity and selective enrichment in CI psychopathology, partially recapitulating the broader profile highlighted by the latent analysis.

We assessed whether this distinct pattern of information-reflection behavior could be an artifact of inattention, poor task comprehension, or poor model fit rather than a true decision-related phenotype (Supplementary Figure 17). At the item level, only two of the nine instruments separated the groups, and both are CI-loaded (EAT-26, median 58 vs. 51, *z* = 2.76, *p* = 5.8 × 10^-3^; OCI-R, median 17 vs. 11, *z* = 3.02, *p* = 2.5 × 10^-3^; two-tailed rank-sum tests, uncorrected), with every AD-loaded instrument non-significant (SDS *p* = 0.88, STAI-T *p* = 0.45, AES *p* = 0.65; Supplementary Figure 17A) — the item-level counterpart of the selective CI elevation in Figure 6E. The subgroup was not slower: choice reaction times were in fact faster than the rest of the sample in both contexts (medians 1373 vs. 1599 ms in gain, *U* = 87,287, and 1536 vs. 1712 ms in loss, *U* = 90,946; both *p <* 0.0001, two-sided Mann–Whitney *U* tests), with no significant group difference in the collection or fixation responses (*p* ≥ 0.30 and *p* ≥ 0.069, respectively) and no group × context interaction (*p* = 0.94 for choice; Supplementary Figure 17C), and it did not differ in total questionnaire completion time (*U* = 118,954, *p* = 0.95; Supplementary Figure 17D), so membership is not a by-product of hurried responding. Their individual choice GLMs were fit as well as everyone else’s in gain and only slightly less well in loss (median accuracy 0.833 vs. 0.833, *p* = 0.69 in gain; 0.780 vs. 0.807, *p* = 9.2 × 10^-3^ in loss; two-sided rank-sum tests, with a non-significant group × context interaction, *p* = 0.063; Supplementary Figure 17E), so they are not a pool of noisy participants. Restricting to the 1,526 participants who answered the task-comprehension question correctly in both contexts, 73 of the 130 members are retained and occupy the same region of the preference plane (Supplementary Figure 17G), so the pattern is attenuated but not created by comprehension failures. In a post-hoc, exploratory check we also refit the population GLM with the comprehension indicator added to the moderator block; across the 30 psychopathology-interaction terms the adjusted and published coefficients correlate at *r* = 0.994, with 29 of the 30 terms keeping their sign (the single exception is the smallest term, |*β*| ≈ 10^-4^), the adjustment acting as a near-uniform shrinkage of about 12% (Supplementary Figure 17H). The subgroup also did not differ from the rest of the sample by age or gender (age: two-sided Mann–Whitney *U* = 117,858, *p* = 0.92; gender: *χ*^2^(1) = 0.06, *p* = 0.81, over the *n* = 1,913 participants reporting binary gender; Supplementary Table 1). Overall, these analyses suggest that this subpopulation reflects a true decision-related phenotype.

Finally, we asked whether this behaviorally defined subgroup was associated not only with a broad elevation in CI psychopathology, but also with a more specific pattern of individual symptoms. We trained an L1-regularized logistic-regression classifier to classify in vs. out of this subpopulation based on the 209 questionnaire items, using nested crossvalidation (Methods). Symptom responses predicted subgroup membership modestly but significantly above chance in held-out participants (area under the receiver operating characteristic curve; AUC = 0.64; balanced accuracy = 0.60; Figure 6G). Repeating the entire nested procedure on 1000 label permutations yielded no permutation reaching the observed AUC (AUC *p <* 0.001; balanced accuracy *p* = 0.002; Supplementary Figure 17F), so neither overfitting nor the selection of the regularization strength is a plausible account of the result. Although this discrimination is far below what would be needed for diagnostic classification, it shows that individual symptom profiles carry reproducible information about membership in this independently defined behavioral phenotype.

L1 regularization allows feature selection by setting many coefficients to zero, and the fitted solution retained only 6 of the 209 items (Figure 6F): two from the schizotypy scale (SCZ; “Can some people make you aware of them just by thinking about you?”; acting without thinking things over, from a reverse-keyed item), two from EAT-26 (“Avoid food with a high carbohydrate content”; “Have the impulse to vomit after meals”), one from OCI-R (“I feel that there are good and bad numbers”), and one from LSAS (“Eating in public places”); all six items had positive CI factor loadings, four of them substantial, with the remaining two endorsing schizotypy and social-anxiety symptoms whose CI loadings were small. Against a scale-matched permutation null (Methods), the three largest coefficients were individually significant under family-wise error control (SCZ #8 and SCZ #39, both *p* = 0.003; EAT-26 #7, *p* = 0.019), while the three smallest were not (*p* = 0.35, 0.45 and 0.81). Because L1 regularization retains only a minimal predictive subset among many correlated items, even these three should be read as a sparse combination sufficient to carry the signal rather than a uniquely identified or exhaustive list of the symptoms on which the subgroup differs; items correlated with the retained set are likely also elevated. More broadly, the retained features provide an initial illustration of the types of individual symptoms enriched in this behavioral phenotype rather than a set of proposed diagnostic symptoms.

Overall, these analyses highlight a latent dimension of economic changes that discriminate compulsive-intrusive psychopathology. Motivated by one especially salient feature of this pattern, a reduction in information-seeking in loss, we characterized an information-reflecting subgroup whose valuation of non-instrumental information reversed from positive in gain to negative in loss. These individuals also showed weaker reward sensitivity across contexts and higher CI psychopathology, and their individual symptom responses predicted subgroup membership modestly but reproducibly out of sample. Thus, using a dimensional approach to link multi-attribute economic preferences and psychopathology, we identified a computationally defined phenotype in association with a potential psychiatric symptom profile.

## 3 Discussion

Human decision-making about uncertain gains and losses is central to behavior and is profoundly altered in psychiatric illness, yet mechanistic links between economic preferences and dimensional psychopathology remain poorly understood [6, 7, 11, 16, 19, 21, 31, 33, 34, 55, 59]. Information-seeking has also become markedly more prominent in an era of abundant digital information, raising concerns about information quality, tolerance of uncertainty, and costs to individuals and society [76, 77]. But how reward, risk, and information together control our everyday choices, which often tend to incorporate and trade-off among all these attributes, remains poorly understood. Also, though dimensional and computational approaches propose that transdiagnostic symptom factors—such as anxious–depressive, compulsive–intrusive, and social withdrawal—arise from disturbances in core computations including motivation, uncertainty processing, and sensitivity to loss, links between theoretical accounts and multi-attribute economic decision-making remain sparse (but see [7, 9, 11, 16, 54, 55, 59]).

Here, we measured how 1,954 individuals traded off expected reward, reward uncertainty, and uncertainty-resolving information within a single task that directly compares matched gain and loss contexts. This design allows us to characterize how preferences—and their gain–loss modulation—vary across individuals, and to test whether symptom dimensions map onto broad scalings of the strength with which economic attributes influence choice, selective alterations in uncertainty or loss processing, or more complex, low-dimensional latent structure [16, 59].

We find that psychopathology is associated with distinct and context-dependent alterations in economic preferences. Across the population, participants preferred higher expected reward and advance information in both gain and loss contexts, were risk-averse for gains and relatively risk-seeking for losses, and exhibited weaker reward and information preferences under loss framing, extending classic reflection effects to a multi-attribute setting with explicit non-instrumental information [6, 19, 21, 31, 33, 34].

Importantly, individual-level analyses revealed substantial heterogeneity in both the strength and direction of these effects. While many individuals showed consistent information-seeking across contexts, a distinct subset exhibited an information-reflecting pattern—seeking information about gains but avoiding it in loss.

When related to dimensional psychopathology, anxious–depressive (AD) and social withdrawal (SW) factors were associated with increased reward sensitivity (and, for AD, stronger scaling of information value with uncertainty, most reliably in loss), whereas the compulsive–intrusive (CI) factor was associated with weaker reward valuation, reduced risk aversion in gain, and diminished information-seeking and information–uncertainty scaling in loss Critically, these effects showed a hierarchical structure. A broad common component captured substantial psychopathology-related variation in the overall strength with which economic attributes influenced choice, particularly in loss, but systematic attribute- and context-specific deviations remained. These residuals were most prominent for CI and for risk and information valuation, and were not well characterized as simple shifts along the ordinary gain-to-loss preference axis. A data-driven analysis then highlighted a latent dimension contrasting CI with AD/SW, characterized by reduced reward and information valuation—particularly in loss—which converged with an independently identified subgroup selectively enriched for compulsive–intrusive symptoms. Together, these findings show that dimensional psychopathology is linked to structured, low-dimensional variation in multi-attribute economic preferences and that task-defined phenotypes, such as information reflection, provide concrete behavioral targets for computational psychiatry [7, 9, 11, 54, 55].

### 3.1 Multi-attribute decision-making across gains and losses

Our findings of reward and risk preferences are largely consistent with prior work [6, 33, 34]. Our population-level findings replicate the well-known reflection pattern—risk aversion for gains and relative risk seeking for losses. However, whereas prospect theory approaches emphasize curvature of the value function and probability weighting, we directly measured sensitivity to expected value under matched gain and loss frames and found that reward sensitivity itself is reduced in loss, despite trials that were structurally identical across contexts except for framing. Thus, loss framing alters economic decision-making beyond a simple reversal of risk attitude, potentially by recruiting context-dependent affective and attentional processes [6, 33, 78].

We also examined the value of non-instrumental information in gain and loss. Overall, participants consistently preferred informative options even when information could not alter objective outcomes in both contexts, demonstrating that information functions as a genuine economic attribute in both gain and loss contexts [18–21]. Importantly, information was valued even when there was no outcome uncertainty to resolve, while its value increased further with reward uncertainty. Thus, the baseline Info effect and the Info × Unc[Rew] interaction capture complementary aspects of information valuation: a general value assigned to knowing and an additional uncertainty-dependent increment in that value. This uncertainty-dependent scaling was stronger in gain than in loss, indicating that the benefit of resolving uncertainty depends both on its magnitude and on outcome valence [21, 22, 43]. These findings align with theories that information about future outcomes can carry both cognitive value from resolving uncertainty and valence-dependent emotional consequences [19–22, 79].

Participants did not evaluate reward, risk, and information independently: their influences on choice interacted in systematic and context-dependent ways. This underscores the importance of multi-attribute paradigms, as attempts to characterize “risk preference” or “information-seeking” in isolation risk overlooking the structured interactions that govern behavior [30, 31, 34].

Apparent economic preferences could arise due to several mental processes (hidden states) and internal computations. Recent work has argued that some of what looks like risk attitude in humans could be reproduced by riskless objects whose “description” or sensory statistics are artificially complicated, suggesting that preferences partly reflect the difficulty of evaluating an option rather than for example only an attitude toward uncertainty itself [80]. While this work does not address whether, or how, complexity might itself induce uncertainty or risk through additional hidden states, it does highlight important considerations. For example, more generally, apparent curvature in choice can arise from imprecise internal evaluation rather than from the preference parameter it is taken to reflect [81]. Analogous concerns apply to the other attributes here: apparent curvature in reward valuation may partly reflect comparison against a remembered sample of alternatives rather than a fixed utility function [82], and the value of non-instrumental information may reflect the value of information itself [19] along with the cost of processing it, an information-demand analogue of the same complexity concern [83]. Separating these accounts will require designs that vary the processing demands of an offer independently of its economic content, for instance by presenting identical outcomes in simple and complex formats. While in our task, the risk and info attitudes are conserved when tested in non-human primates with visual fractals with fixed complexity, the details of the underlying neural mechanisms in humans with less training ought to be further investigated [23, 84, 85].

Large individual differences highlight that population averages mask substantial computational diversity. Some individuals became more risk-averse or more information-seeking in loss, whereas others showed the opposite pattern, and a distinct subset exhibited an information-reflecting phenotype—seeking information about gains but avoiding it for losses. This heterogeneity is consistent with work showing that economic preferences occupy a multi-dimensional trait space rather than a single continuum and motivates linking such variation to dimensional psychopathology and underlying computations [1, 30, 34, 59, 86].

### 3.2 Psychopathology dimensions and computational mechanisms

By integrating multi-attribute preferences with dimensional symptom scores, we could characterize how specific psychopathology factors relate to reward, risk, and information valuations and evaluate competing computational accounts. AD symptoms were associated with steeper reward sensitivity in both gain and loss, more strongly in loss, and with stronger scaling of information value with reward uncertainty, most reliably in loss. At first sight this conflicts with the blunted reward responsiveness that characterizes depression in reinforcement-learning and effort-based tasks [87–89]. However, the AD factor is a broad internalizing dimension, loading on trait anxiety, depressive symptoms, apathy, and impulsivity; it is not a measure of anhedonia. Indeed, anxiety is associated with heightened weighting of outcomes, particularly potential losses [31, 90], and the AD steepening was largest under loss framing. Moreover, reward sensitivity here is the weight placed on described expected value at the moment of choice, whereas the blunted-reward literature measures how received rewards shape subsequent behavior or mood. There is evidence that these are dissociable processes: valuation of described outcomes is intact in major depression [91], and in a risky-choice task requiring no learning, the neural and emotional impact of reward prediction errors is likewise intact in depression [48]. Our trial-history analysis suggests a similar dissociation, with AD showing no change on how the previous trial’s outcome influenced the next choice (Supplementary Figure 2). Thus, the association between AD and reward sensitivity may be due to heightened outcome sensitivity—greater reliance on expected value and on uncertainty-resolving information, especially when losses are at stake—rather than heightened reward-seeking in the hedonic sense. Future work is needed to investigate the association between anhedonia and reward-sensitivity in gain and loss decisions.

In contrast, the CI factor showed a profile consistent with both motivational blunting and altered uncertainty processing. Prior work has linked CI to deficits in model-based, goal-directed control and to diminished use of feedback [16, 17, 59, 92]. Here, higher CI was associated with weaker reward sensitivity, reduced risk aversion in gain, diminished information-seeking in loss, and attenuated scaling of information value with reward uncertainty (Info × Unc[Rew]), most reliably in loss. These were the most robust symptom–preference associations we observed: CI’s broad dampening of attribute weights was the largest scaling effect of any factor, the reward-sensitivity reduction was evident even in simple median splits on each of the CI-loaded instruments (OCI-R, EAT-26) in both task-order cohorts (Supplementary Figure 6), and higher CI was further associated with weaker modulation of choice speed by the value difference between offers in loss (Supplementary Figure 8). The reduced E[Rew] sensitivity need not imply reduced hedonic response to reward receipt: in our task it indexes how strongly prospective expected value controls choice. Consistent with this distinction, patients with OCD have shown attenuated nucleus-accumbens responses during reward anticipation despite relatively preserved responses during reward receipt [93]. Together with evidence for impaired goal-directed and model-based control in compulsivity [16, 59, 92], this raises the possibility that future reward consequences exert less influence over ongoing choices in individuals high in CI. At the same time, because CI broadly dampened multiple economic attribute weights, reduced reward sensitivity may be one expression of a more general weakening of task-attribute influence rather than a reward-specific deficit.

The loss-specific information effects raise an apparent paradox. Individuals with OCD often engage in compulsive checking and reassurance seeking, behaviors commonly understood as attempts to resolve uncertainty about possible harm, reduce perceived threat, or restore a sense of responsibility or control [94, 95]. Experimentally, unambiguous checking or reassurance information can transiently reduce uncertainty and estimated threat and shift perceived responsibility [96]. Such behaviors therefore need not reflect a generalized preference for information: checking is often subjectively useful because the information is treated as relevant to preventing harm, obtaining reassurance, or deciding that a threat has been adequately addressed. By contrast, the information in our task was explicitly non-instrumental. Once an option was chosen, advance information could neither change the outcome nor enable an action that prevented or mitigated the loss; it only resolved uncertainty sooner. In this setting, the value of knowing may reflect a trade-off between the discomfort of remaining uncertain and the affective cost of learning potentially bad news, a central theme in theoretical and empirical work on information avoidance [19, 20]. Higher CI was associated not only with diminished information-seeking in loss but also with weaker scaling of information value with the amount of reward uncertainty resolved. Thus, rather than reflecting a simple intolerance of uncertainty that uniformly increases information-seeking, CI may involve a contextdependent alteration in the value assigned to uncertainty resolution: information may be sought when it is experienced as useful for reducing threat or restoring control, yet avoided when it merely makes an unavoidable negative outcome known. This account remains speculative, but it provides a testable distinction between compulsive checking and the valuation of non-instrumental information [45, 97, 98].

SW was associated primarily with reward and risk preferences in this nonsocial monetary task: higher SW predicted greater reward sensitivity in both contexts and greater risk aversion in gain, alongside a smaller but significant association with increased information-seeking in loss. Importantly, higher SW was also associated with slower choices in both contexts and with stronger modulation of choice speed by the value difference between offers (Supplementary Figure 8), consistent with reports of increased and biased deliberation in social anxiety [64]. Such increased deliberation could itself contribute to the heightened reward sensitivity associated with SW: more extensive evaluation of the offers would be expected to sharpen the influence of expected value on choice. This pattern is consistent with evidence that social withdrawal is more strongly associated with altered valuation of social rewards and risks than with broad distortions in epistemic valuation, though tasks involving social uncertainty may reveal stronger effects on information-seeking in this domain, where socially anxious individuals show biased seeking and integration of self-relevant information [99–102].

Our model-based analyses suggest that at least one component of psychopathology-related differences could be related to broad changes to economic sensitivity: AD and SW were associated with a stronger influence of economic attributes on choice, whereas CI was associated with a weaker influence. This correspondence to uniform scaling was especially close in the loss context. However, our data do not identify this scaling uniquely with a particular psychological variable. Within a value-based choice model, uniformly scaling the fitted attribute weights is compatible with several related processes, including differences in the strength of subjective valuation, attention or engagement with task attributes, and the scaling factor with which value differences are translated into choice [46, 87, 103, 104]. The absence of psychopathology effects on fixation or outcome-collection responses, together with their specificity to economically relevant choice variables, argues against a trivial generalized psychomotor disengagement account, but future work using independent measures of valuation, attention, and choice consistency will be needed to identify the process underlying this common scaling component.

Against this common component, there were also key deviations in changes to specific attribute preferences. Psychopathology effects were not well described as shifts along the multivariate direction that ordinarily distinguishes gain from loss: after accounting for overlap with scaling, the gain–loss context-change axis contributed little unique explanatory power. Thus, stronger symptom expression did not simply make participants’ preferences globally more “loss-like” or “gain-like.” This constrains a simple negativity-bias or reference-point account expressed along this particular behavioral axis, rather than excluding abnormalities in reference-point setting or adaptation more generally [31, 34, 59]. Uniform scaling itself was also incomplete. Its clearest departures were concentrated in gain, where CI altered risk preference more strongly than predicted while largely sparing information preference from its otherwise broad dampening, and AD showed less enhancement of information preference than its reward sensitivity would predict. A prospect-theoretic reparameterization provided an alternative description of the broad reward-sensitivity component: variation in outcome-magnitude curvature closely paralleled the opposing reward-sensitivity effects of AD/SW and CI. Yet magnitude curvature had little leverage over uncertainty preference and, where it did, predicted changes opposite to the observed CI and SW risk effects; classical prospect theory likewise does not specify the intrinsic value assigned to non-instrumental information [6, 22, 33]. Together, these results could reflect a hybrid architecture in which psychopathology is associated with a low-dimensional common modulation of multi-attribute choice sensitivity, superimposed with selective alterations in computations governing risk and information whose expression depends on outcome context.

### 3.3 A compulsive-intrusive-linked information-reflecting phenotype and computational nosology

To complement the theory-driven accounts, we used a data-driven partial least-squares analysis to identify a latent dimension of shared variation between economic preferences and psychopathology. This latent dimension contrasted CI against AD and SW and loaded on weaker reward sensitivity in both contexts, reduced information-seeking in loss, and diminished Info × Unc[Rew] scaling in loss, suggesting that these aspects of psychopathology may at least, in part, be distinguished by underlying computations related to these economic preferences. The “information-reflecting” subgroup—individuals who were significantly information-seeking in gain and significantly information-averse in loss— had been identified directly from the distribution of Info weights, independently of this component; its profile nevertheless converged with the component, providing mutual validation. This subgroup displayed relatively typical risk preferences but a sharp inversion in information valuation across contexts and was selectively enriched for CI symptoms and a sparse set of OCD-like items from the questionnaire battery [16, 20, 21, 59].

Computational and dimensional frameworks such as the Research Domain Criteria (RDoC) and related initiatives support understanding the underlying constructs and computational mechanisms of psychiatric disorders [7, 9, 11, 54, 56, 105]. Our results link dimensions of economic behavior and psychopathology and potentially identify a task-defined endophenotype—information reflection across gains and losses—that aligns with a specific symptom profile. Such a phenotype and approach could provide well-defined behavioral targets for studies of neural circuitry, genetics, and treatment, and illustrate how dimensional approaches to psychopathology can mutually inform computational models of decision-making [11, 54, 105].

### 3.4 Future directions, limitations, and translational implications

The observation that multi-attribute preference parameters—and especially context-dependent information valuation— carry associations with dimensional symptoms suggests that they can be developed into computational markers of compulsive–intrusive liability and related traits—pending validation in clinical samples and over time—complementing self-report and standard cognitive measures [7, 9, 54, 55]. In principle, brief versions of this task, combined with focused questionnaires, could help identify individuals whose decision-making is dominated by avoidance of loss-related information, guiding targeted interventions aimed at improving uncertainty tolerance, affect regulation around negative information, and beliefs about the value of knowing [20, 45, 79].

An especially informative next step will be to determine why CI is associated with reduced demand for passive information about losses despite the prominent information-gathering and checking behaviors associated with compulsivity. In the present task, information is explicitly non-instrumental: learning the outcome cannot change what happens next. Future experiments could orthogonally manipulate outcome valence, uncertainty, and the actionability or controllability of information—for example, comparing cues that merely reveal an unavoidable outcome with cues that can guide a subsequent choice. Such designs could distinguish an altered intrinsic value of uncertainty reduction from aversion to negative foreknowledge or an abnormal dependence on the opportunity to act on newly acquired information [20, 45, 97, 98].

Future work should test this task and simplified variants in clinically identified populations, examine how gain loss economic behavior co-evolves with the disease course and treatment, and link these behavioral phenotypes to underlying neural circuits [106]. Current diagnostic frameworks remain coarse—largely questionnaire-based across both clinical and online settings, a limitation our own symptom measures share—and often fail to capture individual-level behavioral and computational variation, particularly in non-stationary conditions such as mental illness [107]. We therefore propose that combining multi-attribute decision tasks, which reveal low-dimensional structure in individual decision-making, with continuous measures collected during daily behavior [106] can enable identification of individual-specific computational profiles that track symptom dimensions over time, supporting a more precise and dynamic approach to diagnosis and intervention.

More broadly, our results show how large-scale, multi-attribute decision tasks can be combined with dimensional psychopathology measures to reveal latent computational phenotypes—such as the CI-linked information-reflecting profile—that cut across diagnostic categories and can inform dimensional, task-based approaches to psychiatric classification [7, 9, 11, 54, 56].

For decision science, these findings support valuation of information as a core, context-sensitive dimension of economic choice that can exhibit reflection-like patterns and interact with reward and risk. For computational psychiatry, they provide a proof of principle that multi-attribute tasks can uncover low-dimensional latent structure linking behavior and symptoms and define interpretable subgroups that help bridge computational models and clinical translation.

Information reflection itself also warrants direct study as a behavioral phenomenon. The striking directional asymmetry—information-seeking for gains coupled with information avoidance for losses—suggests a context-dependent phenotype that is not captured by a single continuum of general information-seeking. Replication with different tasks will be important to establish whether this pattern generalizes across forms of uncertainty and information and to determine the stability of subgroup membership over time. Its selective association with CI and with a specific symptom profile raises the possibility that information reflection marks a narrower computational subtype within compulsive–intrusive psychopathology. Establishing that possibility will require replication in clinical samples, prospective characterization, and tests of whether the phenotype predicts clinically meaningful outcomes or treatment response before it could be considered a target for precision therapeutics.

In conclusion, tasks that enable precise characterization of decision-making processes—and of where those processes break down in mental illness—should ultimately allow behavior to be linked to underlying circuit-level mechanisms. As phenotyping of psychiatric dimensions becomes more refined, such approaches provide a foundation for developing targeted, circuit-informed interventions tailored to individual computational profiles [106].

## 4 Methods

### 4.1 Participants

We recruited a total of *N* = 1,974 adult participants online through Prolific; they provided informed consent according to procedures approved by the Washington University in St. Louis Institutional Review Board. We paid participants a base rate plus a performance bonus. For inclusion we required fluent English and a minimum prior task-approval rate of 95%. The recruited sample had a mean age of 35.95 years (SD = 9.62, median = 35, range 18–57; age was available for 1,973 of 1,974 participants, and we excluded one implausible recorded value, so age statistics reflect *n* = 1,972) and self-identified as female (*n* = 1,223; 61.96%), male (*n* = 709; 35.92%), non-binary (*n* = 32; 1.62%), preferred not to disclose (*n* = 5; 0.25%), or did not provide gender information (*n* = 5; 0.25%). After excluding participants who did not produce convergent per-context choice models (see “Choice generalized linear models” below), we retained *N* = 1,954 participants for the analyses reported below.

### 4.2 Task design

Each participant completed two 150-trial blocks of an economic choice task adapted from [23] for gain and loss contexts, with block order counterbalanced. In the gain context, each trial could add coins to a running bonus; in the loss context, participants began with an endowment from which coins could be subtracted. Trials in the two contexts were structurally identical except for framing.

On each trial, participants chose between two simultaneously presented offers depicted as four vertical stacks of coins representing possible reward outcomes. We characterized each offer by:

- *Expected reward* (*E*[Rew]), the mean of its outcome distribution (positive in the gain context, negative in the loss context).
- *Reward uncertainty* (Unc[Rew]), the standard deviation of its outcome distribution. We used three discrete distribution types: certain (single deterministic outcome, SD = 0), intermediate (four equiprobable outcomes, SD ≈ 2.83), and risky (two equiprobable outcomes, SD = 4.0).
- *Information* (Info), a binary attribute indicating whether the offer would be displayed with an advance cue revealing which of its outcomes would be delivered. Information was explicitly non-instrumental: it did not alter the realized outcome, and Info and Noinfo choices contributed equally to total earnings.
- *Position* (Pos), the left or right screen location of the offer, which we included as a nuisance regressor to absorb side biases.

In addition to these four offer attributes, two interactions were of theoretical interest, and we included them in all regression models: Info × *E* [Rew], capturing whether the value of information depended on offer magnitude, and Info × Unc[Rew], capturing whether the value of information depended on offer uncertainty.

### 4.3 Psychopathology measures

Each participant completed nine self-report instruments (209 items total) covering symptoms across multiple domains: the Eating Attitudes Test (EAT-26), Alcohol Use Disorders Identification Test (AUDIT), State–Trait Anxiety Inventory– Trait subscale (STAI-T), Zung Self-Rating Depression Scale (SDS), Obsessive–Compulsive Inventory–Revised (OCI-R), the Short Scales for Schizotypy, Apathy Evaluation Scale (AES), Barratt Impulsiveness Scale (BIS-11), and Liebowitz Social Anxiety Scale (LSAS). We applied instrument-specific reverse-coding where required, and averaged the LSAS fear and avoidance ratings per item. (We collected a tenth instrument, the Ruminative Responses Scale, but did not include it in the factor analysis.) We *z*-scored item scores across the *N* = 1,954 retained participants, imputed missing responses by iterative multiple imputation (MICE-style, 50 iterations), and re-standardized the imputed scores.

Following [16], we re-estimated that study’s three-factor structure using the same factor-analytic approach, with the number of factors fixed a priori at three: an exploratory maximum-likelihood factor analysis on a heterogeneous (polychoric/polyserial) item correlation matrix with oblimin rotation, loadings freely estimated (`psych` package in R). The recovered factors matched the identities of the original structure — an Anxious–Depressive (AD) factor loading most heavily on the STAI-T, SDS, and AES; a Compulsive–Intrusive (CI) factor loading on the OCI-R, EAT-26, and AUDIT; and a Social Withdrawal (SW) factor dominated by the LSAS — and the loading pattern was qualitatively consistent with the published three-factor solution (visual comparison; the oblique factors were correlated, e.g. AD–SW *r* = 0.52). Figure 1B displays the 209 item loadings as returned by the estimation (the signs of oblique factors are not standardized). We extracted Thurstone regression scores for AD, CI, and SW and *z*-scored them across the sample for use as covariates.

### 4.4 Model-free choice curves

To directly assess how attributes traded-off to affected participant choices, we computed three model-free psychometric curves describing choice as a function of the difference in expected reward between the two offers:

1. *Reward sensitivity.* On all trials, we computed the probability of choosing the right-hand offer as a function of the signed difference in expected reward between the right and left offers (no other matching applied). This indexes overall reward sensitivity.
2. *Risk preference.* On the subset of trials in which one offer was certain (SD = 0) and the other risky (SD = 4.0, the equiprobable two-outcome distribution), we computed the probability of choosing the risky offer as a function of the difference in expected reward between the risky and certain offers.
3. *Information preference.* On the subset of trials in which one offer was Info and the other Noinfo (no further matching of reward or distribution), we computed the probability of choosing the Info offer as a function of the difference in expected reward between the Info and Noinfo offers.

For each curve, each context, and each participant, we fit a two-parameter logistic function *P* = 1*/*(1 + exp(-(*β*_0_ + *β*_1_ Δ*E*[Rew]))) without a lapse term, in which the slope *β*_1_ measures sensitivity to expected reward in that comparison and the intercept *β*_0_ measures preference for the target attribute (right offer, risky offer, or Info offer) at Δ*E* [Rew] = 0. We obtained population curves by fitting the same two-parameter logistic form to all participants’ trials stacked together — a trial-level binomial fit containing only the intercept and slope terms, distinct from the multi-attribute pooled GLM described above — with CR0 participant-clustered sandwich standard errors (indifference points computed via the delta method).

As a complementary model-free preference measure (Figure 2B), we computed each participant’s fraction of choices in favor of the offer with the higher value of each attribute (with the loss-context *E* [Rew] comparison sign-flipped, as above) and tested the population mean of each fraction against chance (0.5) with one-sample *t* tests, BH-corrected across the six tests (three attributes × two contexts); we report the accompanying gain-vs-loss difference tests uncorrected.

### 4.5 Tertile-stratified analyses

To visualize how each psychometric curve varied with psychopathology, we re-fit the model-free curves above within tertile-defined strata of each factor (top vs. bottom 33% by percentile cuts; middle tertile excluded), using per-stratum cluster-<u>robust logisti</u>c fits. We compared strata by Wald *z* tests with independent group standard errors, 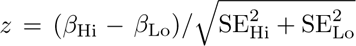. For each curve we tested a single parameter — the slope for the *E* [Rew] curves and the intercept for the Unc[Rew] and Info curves — and BH-corrected the resulting 18 tertile contrasts jointly with the 6 population-level tests as a single 24-test pool.

### 4.6 Reward-uncertainty contingent information seeking

To test directly whether information seeking depended on the level of reward uncertainty, we restricted analysis to trials on which the two offers differed in Info but had matched distribution types (both certain, both intermediate, or both risky). Within each context and for each of these three uncertainty levels we computed each participant’s proportion of choices directed to the higher-Info offer, yielding a three-point curve per participant and context. We tested the monotonic trend of information seeking with reward uncertainty, separately in each context, with Page’s *L* trend test on the per-participant values, assessing significance by within-participant permutation of the uncertainty-level labels (10,000 permutations, two-sided). A cluster-robust linear trend model gave consistent results and we treat it as a sensitivity check.

We complemented this with a trial-level cluster-robust logistic GLM in which the dependent variable was “chose Info,” the predictors were the standardized expected-reward difference between the two offers, a binary risky-vs-certain indicator, the three psychopathology factors (continuous), and the risky × factor interactions, fit separately within each context. We based statistical inference on the continuous risky × factor interaction terms. For visualization (Figure 4C), we computed model-predicted probabilities of choosing Info at Δ*E* [Rew] = 0 for the certain and risky conditions, evaluated at the mean factor score of the above- and below-median groups of each factor, normalizing each line to its own certain-condition value so that the displayed quantity represented the change in information seeking caused by introducing reward uncertainty.

### 4.7 Choice generalized linear models (GLM)

#### Individual GLM

For each participant and each context separately, we fit a Bernoulli generalized linear model with a logit link predicting the binary choice from the differences in offer attributes (offer 2 minus offer 1):

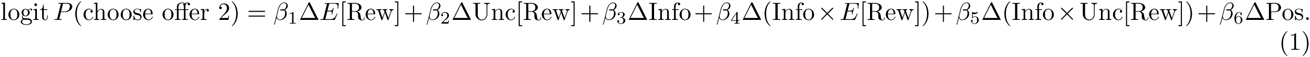

We included no intercept. Within each participant and context, we *z*-scored each attribute across the two offers (pooling both offers’ values across trials), formed interaction terms as products of the *z*-scored attributes within each offer, and then took each regressor as the offer 2 minus offer 1 difference. We entered position as a raw ±1 side indicator and did not *z*-score it. For display and for all downstream participant-level summaries, we sign-flipped the loss-context *E* [Rew] and Info × *E* [Rew] coefficients so that positive values indicate reward-seeking in both contexts; individual-level significance used each fit’s Wald *p*-values. The fitted *β*’s are individual-level economic preferences and constituted the primary input to all subsequent participant-level summaries. We excluded participants whose model failed to converge in either context (optimizer warnings). We froze the individual model fits — and therefore the exclusion itself — in a cached fit object that all downstream analyses load rather than re-estimating from raw data (see “Data and code availability”).

#### Population GLM

To estimate population-average effects we additionally fit pooled trial-level GLMs jointly to all trials and all participants, with CR0 (Liang–Zeger) participant-clustered sandwich standard errors [108]; in the Results and figure captions we refer to these pooled fixed-effects fits as population GLMs (population models). This approach directly targets population-average preferences while accounting for within-participant trial dependence and is appropriate when individual-level variance components are not the inferential focus [109, 110].

For these pooled models, we constructed regressors on a single common scale across all participants and contexts: we converted per-offer attribute values to absolute values and *z*-scored them using moments pooled across both offers, all trials, all 1,954 participants, and both contexts; formed interaction terms as products of the *z*-scored attributes; then negated *E*[Rew] and Info × *E* [Rew] on loss-context trials (so that losses carry negative value); and finally took each regressor as the offer 2 minus offer 1 difference. Unlike in the per-participant GLMs, we also absolute-value transformed and *z*-scored Pos in this pooled pipeline. We sum-coded context as +0.5 for gain and −0.5 for loss, so each regressor’s main effect represents the average of its gain and loss effects and the regressor × context interaction equals the full gainminus-loss difference. Where useful, we re-expressed coefficients as context-specific effects with *β*_gain_ = *β* + 0.5 *β*_ctx_ and *β*_loss_ = *β* - 0.5 *β*_ctx_, propagating standard errors through the cluster-robust covariance matrix using the delta method. We used the same construction to compare two coefficients of the same fit within one context — specifically, whether the pooled model weighted the Info × Unc[Rew] interaction more strongly than the Info × *E*[Rew] interaction (Supplementary Figure 1B): we tested the difference of the two context-specific effects as a linear contrast of the four underlying coefficients, with the contrast variance taken from the corresponding block of the cluster-robust covariance matrix (two-sided Wald *z* tests, BH-corrected across the two per-context contrasts as their own family). Because all attribute regressors share the single common *z*-scale described above, the two interaction coefficients are directly comparable; the loss-context comparison pairs the unflipped Info × Unc[Rew] coefficient with the sign-flipped (reward-seeking-positive) Info × *E* [Rew] coefficient, and as both context-specific estimates are positive the contrast also equals the difference in magnitude.

The main pooled choice model contained the six attribute regressors, each crossed with {1, context} and {1, AD, CI, SW} — that is, attribute main effects, attribute × context, attribute × factor, and attribute × context × factor interactions (6 × 2 × 4 = 48 coefficients) — with no intercept and no context or factor main effects. We used two analogous variants of the same pooled specification for control analyses (see “Trial-history and reaction-time models” below). As a check for block-order effects, we additionally re-fit the pooled model separately for participants who experienced the gain block first and those who experienced the loss block first, and contrasted the two groups’ coefficients with independent-variance Wald *z* tests (unadjusted; inset of Figure 3C and boxed panel of Supplementary Figure 5).

#### Significance testing and multiple-comparisons correction

We assessed the significance of fixed effects by Wald *z* tests (normal approximation; also used for the linear reactiontime models below). Our default correction for multiple comparisons was Benjamini–Hochberg FDR applied across the context-decomposed gain and loss effects of a model — for the main choice model, the 48 effects formed by the 24 fitted terms × 2 contexts — rather than across the raw fitted coefficients. We tested context-dependence of the factor effects themselves (annotated with arrows in Figures 3C and 4D) on the twelve attribute × context × factor coefficients (4 economic attributes × 3 factors), BH-corrected as a single family; every term surviving this family correction is marked with an arrow. For figures showing per-participant population summaries, scatter plots, or context-decompositions of the same model, we applied the analogous grouping — correcting jointly across the same family of effects so that the figure-level family-wise rule matched the pooled-model family. Where deviations from this default were necessary we note them with the relevant figure. For population averages of the individual-level GLM weights, we BH-corrected one-sample *t*-tests over the 12 regressor-by-context tests (6 regressors × 2 contexts), and the corresponding tests of the per-participant gain-minus-loss differences over their own family of 6. To quantify across-individual heterogeneity in per-participant GLM estimates, we computed Cochran’s *Q* statistic and the *I*^2^ index for each regressor and context, and analogously for the per-participant gain-minus-loss differences; *Q* tests the null hypothesis that all individual estimates share a common population value, and *I*^2^ expresses the percentage of total variability attributable to genuine between-individual differences rather than sampling error. To test whether the observed proportion of individuals with significant per-participant effects exceeded the rate expected by chance (2.5% in each tail at *α* = 0.05), we used exact binomial tests. We tested per-participant gain-versus-loss differences <u>in a given coe</u>fficient with a two-sided *z* statistic combining the two independent per-context fits, 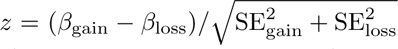, and compared the proportion of individuals significant in a given direction against the same 2.5% chance rate with exact binomial tests. For the joint gain × loss sign classifications (Supplementary Figure 3A), we tested each cell of the 3 × 3 table (significantly positive / not significant / significantly negative in each context) with a one-tailed enrichment binomial test against the cell proportion expected under independence of the two contexts’ chance rates; stars mark over-represented cells only (uncorrected *p*). These enrichment tests treat each per-participant Wald test as exactly calibrated at its nominal level; with ∼150 trials per context, modest deviations from nominal calibration would shift the expected tail rates, so the smaller enrichment counts are best read as descriptive.

#### Questionnaire-level median splits

To check whether the factor-level associations were carried by a single instrument or by the order in which the two contexts were experienced, we repeated the group-average preference display for each of the nine questionnaires separately (Supplementary Figure 6). For each instrument we formed a per-participant score as the mean of that instrument’s item-level *z*-scores, taken from the same imputed, re-standardized item matrix that entered the factor analysis, and split participants at the 50th percentile of that score across the *N* = 1,954 retained sample (above the cutoff, high; at or below it, low). We then intersected each half with the task-order cohort — gain-first (n=979) or loss-first (n=975) — giving 18 questionnaire × cohort comparisons of 455–520 participants per group. Within each group we averaged the per-participant GLM weights of each of the six attribute regressors, separately for the gain and the loss context, and plotted the two group means with their standard errors. We compared the high group against the low group on each context axis wi<u>th a two-tailed two</u>-sample *t* test computed from the two group means and their standard errors, 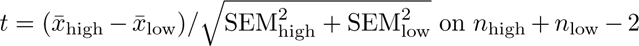 degrees of freedom. These tests are descriptive and we report them uncorrected.

### 4.8 Tests of how psychopathology reshapes the preference vector

A central inferential question is whether psychopathology factors reshape the population’s economic preference vector by uniformly scaling it within each context, or by introducing idiosyncratic per-attribute distortions. We addressed this with two complementary tests, both derived from the single pooled trial-level choice GLM described above.

#### Preference axes and factor coefficient vectors

We obtained all quantities in this analysis by exact linear reparameterization of the fitted 48-coefficient pooled choice model and its cluster-robust covariance matrix, not from any separately estimated model. Working within a single fitted model and a single covariance matrix keeps every comparison an exact contrast, rather than an approximation across separate re-fits. With the ±0.5 sum-coding of context, the population preference weights for the four well-estimated economic attributes — E[Rew], Unc[Rew], Info, and Info×Unc[Rew] — are 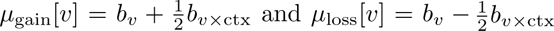. The context-change vector is *δ* [*v*] = *b*_v_**_×_**_ctx_ = *µ*_gain_ [*v*] - *µ*_loss_ [*v*]. We excluded Info × *E* [Rew] and Pos from the economic axes; we retained Pos as a nuisance regressor in all constrained re-fits. For each factor *f* ∈ {AD, CI, SW}, we obtained the unconstrained 8-dimensional factor coefficient vector 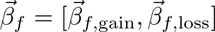, with components 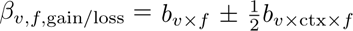, and its 8 × 8 cluster-robust covariance *V*_8_. We obtained both by the corresponding sparse linear transformation of the pooled coefficient vector and its participant-clustered (CR0) sandwich covariance.

#### Scaling test

The scaling hypothesis holds that each factor rescales each context’s own population preference axis: 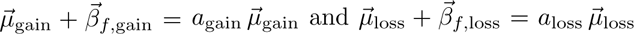. Equivalently, 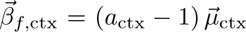, so that *a* = 1 corresponds to no factor-related change. Under this hypothesis the 8-D factor vector lies in the 2-D subspace spanned by 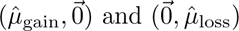. The fixed weights *μ̂*_gain/loss_ lock the relative contribution of each attribute, leaving only the overall amplitude in each context free (6 degrees of freedom per factor; 8 unconstrained → 2 constrained). We tested whether alignment was perfect by a cluster-robust Wald statistic on the 6-D orthogonal complement of the constrained subspace (orthonormal basis *R* obtained by QR decomposition; 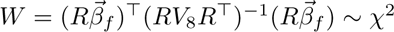(6) under *H*_0_). We retain this joint statistic for the maximum-likelihood version of the test (Supplementary Figure 12C, below).

For the version of this test we use in the main text, we do not fit the per-context scalar but pin it to the factor’s own E[Rew] interaction: *a*_ctx_ = 1 + *β*_R,f,ctx_*/µ*_R,ctx_. Here *β*_R,f,ctx_ is factor *f*’s unconstrained E[Rew]-interaction coefficient in that context and *µ*_R,ctx_ is the corresponding population-average weight. The standard error follows by the delta method, se(*a*_ctx_) = se(*β*_R,f,ctx_)*/µ*_R,ctx_, treating *µ*_R,ctx_ as fixed. We report per-scalar tests as two-sided Wald *z* tests of *H*_0_: *a*_ctx_ = 1, *z* = (*a*_ctx_ - 1)*/*se(*a*_ctx_). Pinning has two advantages. It requires no re-fitting, because the scalar is available in closed form. It also forces the constrained model to reproduce the unconstrained E[Rew] weight exactly in each context, so the test asks whether the remaining attributes follow the scaled population-average pattern rather than crediting the fit for reproducing E[Rew] itself.

Every constrained quantity is then a fixed linear function of the pooled model’s coefficient vector 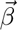. All comparisons between the unconstrained and constrained fits therefore reduce to exact linear contrasts under the pooled cluster-robust covariance *V*_8_. The per-attribute residual for attribute *i* is *d*_i_ = *β*_i_ - (*µ*_i_*/µ*_R_)*β*_R_, with Var(*d*_i_) = *V*_ii_ + *k*^2^*V*_RR_ - 2*kV*_iR_ for *k* = *µ*_i_*/µ*_R_; independence-SE formulas are invalid here because the constrained prediction shares *β*_R_’s sampling noise. The E[Rew] contrast is exact by construction and we report it as *p* = 1. As in the maximum-likelihood version, we BH-correct these per-attribute contrasts within each factor’s eight attribute-context comparisons, with the two pinned E[Rew] slots entering at *p* ≡ 1. We likewise tested the gain-versus-loss difference in the scaling scalar as an exact linear contrast using the full cross-context covariance, *z* = (*a*_gain_ - *a*_loss_)*/*se(*a*_gain_ - *a*_loss_). We computed that standard error from *V*_8_ rather than assuming independence between contexts.

We retain the maximum-likelihood version of this test in Supplementary Figure 12 (statistics unchanged from the original fit). There we instead estimated *a*_gain_ and *a*_loss_ by re-fitting a logistic GLM with two composite regressors per factor, replacing that factor’s eight interaction columns by 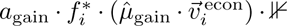[gain] and the analogous loss term, with 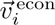 = (E[Rew], Unc[Rew], Info, Info × Unc[Rew])_i_.

#### Context-change (*Δ-axis*) test

A complementary test asked whether each factor’s effect lay along the single direction in attribute-coefficient space along which population preferences themselves changed between contexts: 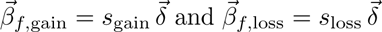. The constrained subspace is spanned by 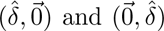, and the Wald construction is identical to that of the scaling test (*W* ∼ *χ*^2^(6) under *H*_0_). The corresponding two-scalar re-fit used the single composite direction 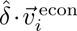 in both contexts, with free per-context amplitudes *s*_gain_ and *s*_loss_. Unlike in the scaling test, we did not pin this test’s scalars. Pinning requires a single attribute that dominates the reference axis, but 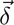 is spread across attributes (E[Rew] carries only 10.5% of 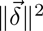 versus 61.6% for E[Rew] within 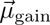). Pinning to E[Rew] here is therefore numerically unstable (scalars 4–50× the maximumlikelihood values). The per-context alignment statistics below are scale-invariant in the constrained scalar and identical under either choice.

#### Cluster-robust covariance

The Wald covariance *V*_8_ derives from the pooled model’s participant-clustered (CR0) sandwich covariance via the linear transformation above. The scalar standard errors from the constrained re-fits used a logistic Huber–White sandwich clustered by participant with a *G/*(*G* - 1) finite-cluster correction.

#### Calibrated above-chance null

A *χ*^2^ (6) test of perfect alignment is expected to reject any point null at this sample size, so the informative comparison is whether alignment was better than chance. We defined chance by a calibrated directional null that randomizes the effect direction rather than the reference axes. Each null draw generates 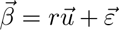, with 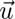 an isotropic random unit direction in ℝ^8^ and 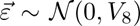 drawn from the factor’s full cluster-robust covariance. We matched the signal magnitude *r* to the observed effect size (*r*^2^ = max(0, ||*β̂*||^2^ - tr *V*_8_)). The one-tailed *p*-value is 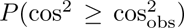 over 10^6^ draws (fixed seed). This construction places isotropy on the unaligned effect — which has no preferred direction — rather than on the estimated reference axes. We did not use the naive alternative that redraws the reference axes themselves as isotropic random unit vectors, because it is anticonservative for these strongly anisotropic axes (measured Type I error ≈ 0.12–0.13 at nominal 0.05).

#### Alignment (cosine) reporting

We report signed cosine similarities between the unconstrained factor vector and the constrained axis separately for gain and loss. For the joint (8-D) quantities retained in Supplementary Figures 12 and 14, we report the squared cosine between the unconstrained 8-D coefficient vector and the fitted constrained profile ([*a*_gain_ *µ̂*_gain_; *a*_loss_*µ̂*_loss_] for the scaling test; [*s*_gain_*δ̂*; *s*_loss_ *δ̂*] for the Δ-axis test). Because this hypothesis space is two-dimensional, the expected value of cos^2^ for a random direction is 2*/*8 = 0.25. The dashed chance line in the figures marks this expected value under chance, not a significance threshold. Joint error bars on cos^2^ are the standard deviation of a 10,000-draw parametric bootstrap perturbing the coefficients by their standard errors with the constrained fit held fixed across draws (fixed seed). Holding the fit fixed is an approximation, since refitting the model on every draw is infeasible. The displayed error bars — together with the per-attribute agreement standard errors of the maximum-likelihood version and the competitive-decomposition bootstrap of Supplementary Figure 14E — perturb the coefficients independently for simplicity, whereas the cluster-robust Wald *χ*^2^ (6) test of perfect alignment and the calibrated above-chance null both use the full *V*_8_, as do the main-text pinned contrasts of Figure 5, which are computed exactly under the pooled covariance.

For the main-text scaling test (Figure 5C) and its Δ-axis counterpart (Supplementary Figure 14C), we instead score alignment per context: within each 4-D context block, 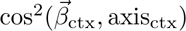 is scale-invariant in the constrained scalar. We computed two tests per context bar. The first is a deviation-from-perfect cluster-robust Wald test on the 3-D orthogonal complement of the axis within that 4-D block (*χ*^2^ (3) under perfect alignment). The second is a calibrated above-chance test using the same directional-null logic as the joint test (Methods above), restricted to the 4-D context block 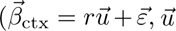 isotropic in 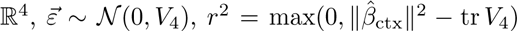), 10^6^ draws, fixed seed); chance is E[cos^2^] = 1*/*4 for a 1-D target in 4-D under isotropy. For a gain-versus-loss comparison of the per-context cos^2^ values we used a two-sided percentile *p* from a parametric bootstrap of the full joint covariance *V*_8_ (10^4^ draws, fixed seed), respecting the covariance between contexts. Error bars on the per-context cos^2^ are central 68% percentile intervals from a full-covariance parametric bootstrap, drawn as asymmetric [lo, hi] segments, as cos^2^ is bounded at 1 and left-skewed near that ceiling.

#### Subspace-projection robustness check

We also used a complementary alignment measure (Supplementary Figure 12D for the scaling test and Supplementary Figure 14D for the Δ-axis test) that evaluates how well the unconstrained model aligns with the subspace of vectors defining the plane of possible scaling hypotheses. For it we computed the squared cosine of the orthogonal projection onto the two-scalar hypothesis space, 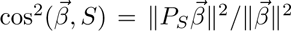 with 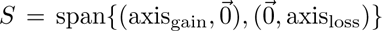. This is the same subspace whose 6-D orthogonal complement the Wald statistic tests. For this statistic the bootstrap recomputes the projection on every draw, and both the ceiling (exactly 1) and the chance expected value (exactly 2*/*8) are analytic. 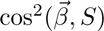 necessarily bounds the constrained-fit statistic from above, since the projection is the angle-optimal point in the subspace while the constrained model is the likelihood-optimal one.

#### Per-attribute and scalar-difference tests (maximum-likelihood version, Supplementary Figure 12)

For the maximum-likelihood scaling fit, we furth<u>er tested agree</u>ment between the unconstrained and constrained models per attribute by two-sample *z* tests with 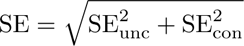. We BH-corrected these within each facto<u>r (three pools o</u>f eight). To test whether the scaling coefficient differed between contexts we computed 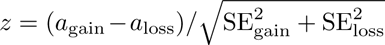, treating the two scalars as independent (Supplementary Figure 12). Under the pinned scaling test used in the main text, the analogous per-attribute and gain-versus-loss tests instead use the exact cross-context covariance, as described above.

#### Competitive decomposition

The two reference axes are not orthogonal (cos^2^ (*µ̂*, *δ̂*) = 0.376), so the single-axis squared cosines are not additive. To disentangle the contributions of the two candidate directions (Supplementary Figure 14E), we orthogonally projected each factor’s unit-normalized 8-D coefficient vector onto the plane spanned by the unit vectors *µ̂*_8D_ (the concatenated gain/loss population axes) and *δ̂*_8D_ (the context-change direction repeated in both context blocks). The joint projection weights disentangle the axes’ shared and unique contributions. We report the projection weights *w*_µ_ and *w*_δ_ and the squared cosines with each axis and with the plane. Confidence intervals are the 2.5th/97.5th percentiles of a 10,000-draw parametric bootstrap (fixed seed). Projection-weight p-values are two-sided normal tests on the bootstrap standard errors, uncorrected.

### 4.9 Cumulative Prospect Theory modeling

To provide a parametric account of the choice data, we fit cumulative prospect theory [33] to the trial-level choices. Each offer comprised up to four possible outcomes with known probabilities determined by its distribution type. We valued outcomes with a power-law value function, *v*(*m*) = sign(*m*) |*m*|_α_, where *α* controls the curvature of the value function (*α <* 1 indicating diminishing marginal sensitivity). We distorted probabilities by a one-parameter Prelec weighting function, *w*(*p*) = exp(-(- ln *p*)^a_w_^), where *a*_w_ controls curvature (we fixed the elevation parameter of the two-parameter Prelec form at 1). We constructed decision weights from the weighted cumulative probabilities, *π*_j_ = *w*(*P*_j_) - *w*(*P*_j_ - *p*_j_), and the prospect value of each offer was *V* = Σ_j_ *π*_j_ *v*(*m*_j_). Choice probabilities were logistic in the prospect-value difference plus a non-CPT offset, *P* (choose offer 2) = *σ*((*V*_2_ - *V*_1_) + *z*) (no temperature parameter), where *z* absorbed the trial-level influence of non-CPT offer attributes (Info, the two Info interactions, and position) on choice. In the population-level fits, we computed *z* from the corresponding fixed-effect coefficients of the pooled choice GLM (including their context and factor interactions); in the individual-level fits, we computed *z* from each participant’s own per-context GLM coefficients on the non-CPT attributes.

The CPT pipeline comprised three fits. First, a two-parameter population fit estimated a single curvature parameter *α* and weighting parameter *a*_w_ jointly from all trials and all participants, providing the population estimate of the probability-weighting parameter, *â*_w_ ≈ 0.65, which we fixed in the two subsequent fits; this allowed individual-level estimation of risk aversion to be identifiable from the relatively small number of trials per participant. Second, with *a*_w_ fixed, an eight-parameter population model allowed *α* (only) to vary as a linear function of context, the three psychopathology factor scores, and the context × factor interactions:

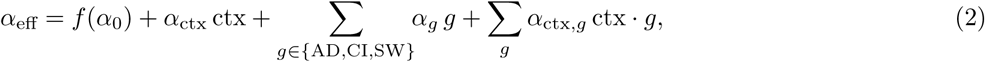

where *f*(*x*) = log(1 + exp(*x*)) is the softplus transform, applied to the baseline term only so that the baseline curvature is positive; the context and psychopathology terms then enter additively on the *α* scale. Across all participants and both contexts, *α*_eff_ remained strictly positive at the fitted estimates (range 0.52 to 1.18). This model provided the context-specific population estimates of *α* (via the context contrasts, with delta-method standard errors) and the psychopathology effects on *α* presented in the Supplementary Material. Third, in the individual-level fits, *α* was the only free parameter (*a*_w_ fixed to the population estimate), and we fit it separately for each participant and each context. The resulting per-participant, per-context *α̂* _gain_ and *α̂*_loss_ provided individual-level indices of risk sensitivity, which we then related to each participant’s context-specific choice GLM coefficients via a linear model (one per context, with the *E*[Rew] and Unc[Rew] GLM coefficients — the two attributes about which the prospect-theoretic value function makes direct predictions — as predictors and *α̂*, on its natural scale, as the outcome). We excluded participants whose individual *α̂* collapsed to exactly zero in either context (degenerate fits; *n* = 312) from this regression analysis, leaving *n* = 1,642; we report regression coefficients with uncorrected two-sided *p*-values.

#### Forward simulation of the α → GLM-weight mapping (Supplementary Figure 15A)

To derive what a change in the curvature parameter *α* predicts for the choice-GLM weights, we simulated the fitted model forward. For each *α* on a dense grid we computed CPT values for all ordered pairs of the nine offer types exactly as in the fitted model (three-outcome prospects at sign(*E* [Rew]) (|*E*[Rew]|+{-4, 0, +4}) coins, distribution probabilities as in the task, Prelec weighting with *a*_w_ fixed at 0.646, choice probability logistic in the prospect-value difference) and refit the no-intercept logistic choice model choice ∼ Δ*E* [Rew] + ΔUnc[Rew] to the deterministic model-implied choice probabilities separately per context, using the pooled regressor convention. We report derivatives as central differences at the published context-specific *α* values; we verified results with stochastic choice simulation and they were insensitive to the *α*-reporting convention. Because the small Unc[Rew] derivative reflects a near-cancellation of curvature- and probability-weighting-driven risk premia, its sign (unlike its negligible magnitude) depends on the assumed *a*_w_; an *a*_w_ sensitivity grid is stored with the figure sidecars.

We estimated parameters by minimizing the negative log-likelihood with unconstrained L-BFGS using analytic gradients computed by forward-mode automatic differentiation in Julia (`ForwardDiff.jl`). To address the multimodal likelihood surface we used a multistart strategy in which each fit was launched from multiple starting points obtained by Gaussian perturbation of a canonical start (eight starts per participant–context fit in the individual stage), retaining the solution with the best likelihood. We used parameter scaling factors derived from the diagonal of the Hessian to normalize curvature across parameters. Statistical inference for population-level parameters used cluster-robust sandwich standard errors: the Hessian *H* at the optimum supplied the model-based variance, while the “meat” 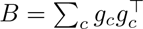 summed per-participant gradients to capture clustering, giving *V* = *H*^-1^*BH*^-1^. For softplus-transformed parameters we adjusted standard errors via the delta method, SE(*f* (*θ*)) = SE(*θ*) · *σ*(*θ*), where *σ* is the logistic function. We assessed significance by two-sided Wald *z* tests. We report no standard errors for the individual-level *α̂* estimates, which enter subsequent analyses as point estimates only.

### 4.10 Partial least squares regression of preferences on psychopathology

To characterize the joint covariation between the three psychopathology factors and the full set of context-specific economic preferences, we fit a partial least squares (PLS) regression. The predictors were the three *z*-scored factor scores (AD, CI, SW). The outcomes were the eight context-specific per-participant choice coefficients: four economic attributes — expected reward, reward uncertainty, information, and information × reward uncertainty — crossed with the two contexts. For this analysis, we took the per-participant coefficients from a five-attribute variant of the choice GLM that omitted the Info×*E* [Rew] interaction (retaining E[Rew], Unc[Rew], Info, Info×Unc[Rew], and position). We then excluded the position bias coefficient from the PLS outcome set, yielding four economic attributes per context (eight outcome variables total). We standardized both the predictor and outcome blocks prior to fitting.

We assessed the number of components by several complementary criteria. We computed held-out mean-squared error by repeated 5-fold cross-validation (10 random shuffles); a 10-fold variant, shown in Supplementary Figure 16, gave consistent results. The remaining criteria were elbow-point detection (Kneedle algorithm), a permutation test on per-component improvement (1,000 permutations of participant labels), a bootstrap stability index of loading sign-agreement, and held-out latent-score diagnostics for the second component (below). Held-out mean squared error (MSE) was minimized at two components, and a one-standard-error rule applied to the same curve also preferred two. The margin over the one-component model was negligible (ΔMSE ≈ 0.0008), however, and the elbow criterion favored a single component.

Our primary model therefore used one component, on grounds of parsimony: the second component added negligible held-out generalization. Because PLS components are estimated sequentially, the first component’s weights and loadings are identical whether one or two components are extracted. The displayed loadings, confidence intervals and bootstrap *p*-values are therefore unaffected by this choice. Component-selection diagnostics, including the second component’s held-out generalization, are shown in Supplementary Figure 16. We obtained confidence intervals on the predictor and outcome loadings from 2,000 nonparametric bootstrap resamples of participants, sign-aligned to the reference solution by Procrustes-style matching. We computed bias-corrected and accelerated (BCa) intervals using a leave-one-out jackknife for the acceleration parameter, and we report 95% BCa intervals as error bars. We assessed significance for coloring by exact two-sided bootstrap *p*-values, *p* = 2 · min(*P* (*b >* 0)*, P* (*b <* 0)) computed from the raw draws of the same 2,000 resamples (granularity 1*/*2,000, so *p* = 0 in the resampling is reported as *p <* 0.001). We BH-corrected these within each block (eight outcome loadings; three predictor loadings).

We tested gain-versus-loss differences between the two context-specific loadings of the same regressor (the identity-line arrows of Figure 6A) with a paired within-replicate bootstrap contrast. We formed the gain-minus-loss difference inside each of the same 2,000 resamples, because the two loadings are correlated across draws and combining independent standard errors would be invalid. We summarized the contrast by the observed difference, its 95% percentile interval, and an exact two-sided *p* = max(2 · min(*P* (*d* ≤ 0)*, P* (*d* ≥ 0)), 1*/*2,000); the floor is needed because a zero proportion is unrepresentable at 2,000 draws. We BH-corrected these contrasts over the four regressor contrasts as their own family, separately from the against-zero tests above. We assessed the omnibus significance of the predictor-to-outcome mapping by a 1,000-iteration permutation test on the participant labels of the predictor block, using the one-component model’s in-sample *R*^2^ (= 0.019). That value is quoted as the equal-weighted mean of the per-outcome-variable *R*^2^ values, 0.0187, while the permutation statistic itself is the variance-pooled form, 0.0190; both round to 0.019.

We calculated latent-score correlations from the X-side and Y-side scores of the one-component model: participants’ standardized predictor and outcome blocks projected onto the model’s weight vectors (unit-norm outcome weights). For a single component these are the scores the NIPALS algorithm estimates, and they coincide with component 1 of any higher-order fit. We computed the cross-validated latent-score correlation *r*_CV_ within the same repeated crossvalidation used for component assessment, so that no participant contributed to the weights that scored them. In each fold, we estimated standardization and all PLS weights from training participants only and then froze them. We projected held-out participants into the training-derived latent spaces, yielding exactly one out-of-fold score pair per participant per repeat (*N* = 1,954). We computed one Pearson correlation per repeat and combined the ten repeat estimates by Fisher-*z* averaging; we report their spread descriptively, because repeats reuse the same participants and are not independent. We fixed component signs by a deterministic training-only rule (CI loading positive). We assessed significance of *r*_CV_ by permuting the participant correspondence between the predictor and outcome blocks and re-running the entire cross-validation pipeline, with identical fold assignments, for each of 1,000 permutations. We report the twosided *p* = (#{|*r*_null_| ≥ |*r*_obs_ |} + 1)*/*(*B* + 1). (An alternative rotation-based score convention is returned by scikit-learn’s `transform`, which mixes components on the outcome side in higher-order fits. Under that convention the same quantities are *r* = 0.257 in-sample and *r*_CV_ = 0.241, two-sided permutation *p <* 0.001; conclusions are unchanged.) We quantified sensitivity of the cross-validated *R*^2^ to component-count selection by nested cross-validation, re-selecting the count by inner 5-fold cross-validation (CV) within every outer training fold (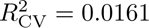, vs. 0.0156 with one component fixed).

### 4.11 Information-reflecting subgroup

We identified an extreme subgroup of participants whose information preferences flipped sign between gain and loss contexts. From the per-participant choice GLM, we defined the information-reflecting subgroup as the set of participants whose Information coefficient was both significant (*p <* 0.05) and positive in the gain context and both significant (*p <* 0.05) and negative in the loss context. To test the directionality of the sign reversal, we restricted to the participants with individually significant Information coefficients in both contexts (*n* = 1,374). We compared the counts of the two opposite-sign patterns (gain-positive/loss-negative vs. gain-negative/loss-positive) with an exact two-sided binomial sign test against an even split.

We then characterized this subgroup along several axes:

- The joint distribution of the six per-participant choice coefficients in the (gain, loss) plane. We highlighted subgroup members against the rest of the population, displayed as a kernel-density background, and contrasted each context-specific coefficient between subgroup and remainder by Welch’s *t* tests, BH-corrected across the 16 tests in the figure.
- The psychopathology factor scores, contrasted between subgroup and remainder by Welch’s ***`t`*** tests on the frozen full-sample factor scores, BH-corrected across the three factors.
- The distribution of total scores on each of the nine self-report questionnaires, contrasted by two-tailed rank-sum tests, uncorrected.
- Per-event task reaction times (choice, fixation, and outcome collection), tested between subgroup and remainder by Mann–Whitney *U* and within subgroup-and-context by Wilcoxon signed-rank tests.
- The time taken to complete the questionnaire battery.
- Two checks against the hypothesis that the subgroup reflected pure noise responding: the per-participant prediction accuracy and per-trial mean log-likelihood of the per-context GLM. We contrasted these between subgroup and remainder by two-sided rank-sum tests, and assessed group × context interactions by rank-sum tests on the per-participant gain-minus-loss differences.

#### Demographic comparisons

Within each group (subgroup and remainder) separately, we quantified associations between demographic variables and the three factor scores by Spearman correlations (continuous variables) and rank-biserial correlations (binary variables), assessing significance by permutation (5,000 permutations, fixed seed). We tested subgroup-vs-remainder differences in these associations by the analogous permutation difference tests. We additionally tested multivariate group differences by permutation Hotelling *T*^2^ and Pillai trace tests (5,000 permutations each) (Supplementary Table 1). For between-group demographic comparisons we used a two-sided Mann–Whitney *U* test for age and a *χ*^2^ (1) test for gender, the latter over the *n* = 1,913 participants reporting binary gender.

#### Item-level prediction

To test whether subgroup membership could be predicted directly from raw questionnaire items, we trained an L1-regularized (lasso) logistic regression classifier (`scikit-learn liblinear` solver, with balanced class weights) on the 209 item scores. We used those scores as pre-processed for the factor analysis: imputation and *z*-scoring performed once on the full sample, with features additionally re-standardized within each cross-validation training fold. We evaluated the classifier by nested cross-validation, with an outer stratified 10-fold loop for out-of-fold performance estimation and an inner stratified 5-fold loop selecting the regularization strength *C* over a seven-value grid, maximizing ROC-AUC. The outer folds took no part in selecting *C*, so the resulting performance estimate is not inflated by that selection. We compute all reported performance from the pooled out-of-fold predicted probabilities, so no participant contributed to the model that scored them. We computed a permutation null distribution of held-out performance by shuffling participant labels and re-running the full nested cross-validation, including the inner search over *C*, on every permutation (1,000 permutations). Re-running the inner search on every permutation lets the null carry the same selection optimism as the observed estimate. We evaluated the permutation null separately for each reported metric (AUC and balanced accuracy). The null AUC had mean 0.499 and standard deviation 0.032, and its maximum over the 1,000 permutations (0.623) fell below the observed value. We then refit the classifier on the full sample to obtain feature importances for the item-level visualization.

ROC-AUC is insensitive to class prevalence, and the positive class comprised only 6.7% of the sample (130 of 1,954). We therefore also report prevalence-sensitive summaries of the same out-of-fold predictions. The area under the precision– recall curve was 0.128 against a chance baseline equal to the prevalence (0.067), a lift of 1.9×. The positive predictive value at the 0.5 decision threshold was 9.9% (71 of 717 flagged participants). These figures indicate that the discrimination, although reliably above chance, is far too weak to identify individuals.

To assess how much the single reported estimate depends on the particular fold assignment, we repeated the entire nested cross-validation under 20 different outer-fold random seeds. Held-out AUC ranged from 0.633 to 0.687 (mean 0.665, SD 0.014). The reported split falls at the low end of this distribution, so the headline estimate is conservative with respect to fold assignment rather than the product of a favorable split.

We assessed statistical significance of individual item coefficients by a separate 1,000-permutation test on the fullsample fit. The magnitude of an L1 coefficient is expressed in units of the penalty. This test therefore holds the regularization strength fixed at the operating value (*C* = 0.01) for the observed fit and for every permutation, so that observed and null coefficients are on the same scale. Re-selecting *C* within each permutation is the correct procedure for the performance statistics above, which are scale-free, and is what we do for them. Applied to a coefficient magnitude it instead compares a sparse, heavily penalized solution against the dense, weakly penalized solutions that an unconstrained search returns when there is no signal for it to lock onto. We report maximum-statistic family-wise corrected *p*-values, which account for the tested coefficients having been selected as the largest among 209 items. We also compute Benjamini–Hochberg values across the retained set, but they are anticonservative in this setting, because each individual item’s null is almost entirely zero. This test is conditional on *C*, which was itself chosen using the observed labels. *C* is a single scalar shared by all items and was stable across folds, so the dependence is minor. The resulting *p*-values should nonetheless be read as conditional on the selected penalty rather than as fully nested.

### 4.12 Trial-history and reaction-time models

To verify that psychopathology-related effects on choice reflected economic preferences rather than non-economic biases, we fit two auxiliary families of pooled trial-level models with cluster-robust standard errors clustered by participant.

The trial-history choice models predicted the current binary choice from (i) a participant-specific value-difference regressor — the dot product of the five attribute differences and their context interactions with the mixed model’s fixed effects plus that participant’s random effects (not *z*-scored) — and (ii) the current-trial position regressor interacted with the previous-trial position × previous-trial choice and with the previous-trial reward prediction error (realized outcome minus expected reward), including the corresponding four-way term; all terms were crossed with context and the three psychopathology factors (40 coefficients). Previous-trial outcome did not enter the model as a separate regressor. The reaction-time (RT) models predicted reciprocal (inverse) reaction times for three trial events with linear models estimated with CR0 cluster-robust standard errors, with a different regressor set per event: choice-commitment RT — the absolute value difference, the signed value sum (the summed subjective values of the two offers, *v*(offer 1) + *v*(offer 2), computed from the same participant-specific weights as the value difference; negative in the loss context), the previous-trial outcome, and the previous-trial reward prediction error; outcome-collection RT — the absolute value difference, the value sum, and the current-trial outcome and reward prediction error; initial-fixation RT — the previous-trial outcome and reward prediction error only. We crossed each set with context and the three psychopathology factors. We excluded trials with missing or non-finite reaction times. For the trial-history and reaction-time models, we applied BH–FDR correction over each model’s own family of context-decomposed effects, following the same rule as the main choice model.

### 4.13 Mixed-model sensitivity check

For all five model families above (main choice, trial-history choice, choice RT, fixation RT, and collection RT), we additionally re-fit each model as a (generalized) linear mixed model, as a check on the pooled cluster-robust fits. The choice models included correlated participant random slopes on all attribute and attribute × context terms but no random intercept. The psychopathology factors are between-participant covariates and received no random slopes. The reaction-time models additionally included random intercepts. As expected, fixed-effect estimates were largely consistent with the pooled cluster-robust analyses. For several high-dimensional specifications, however — especially the reactiontime models and models with extensive random-slope structure — the random-effects covariance was weakly identified (singular fits, near-perfect random-slope correlations). We therefore retained the cluster-robust pooled models as the primary analyses: the mixed models did not alter population-level conclusions and introduced additional assumptions about covariance structure. We report the model-by-model Pearson and Spearman correlations between pooled and mixed fixed-effect estimates as the sensitivity check (Supplementary Figure 4; parametric correlation tests).

### 4.14 Questionnaire engagement controls

To ask whether the psychopathology associations could be produced by careless or disengaged questionnaire responding, we computed two engagement proxies for all *N* = 1,954 retained participants. The two proxies capture complementary signatures of careless responding: how long a participant spent on the battery, and how consistently they answered near-identical items. The first was questionnaire completion time, the time taken to complete the self-report battery. The second was a cross-item reliability score, the psychometric-synonym index [111]. We assembled item responses to 247 questionnaire items — the 209 factor-analysis items plus 38 items from three further instruments (Ruminative Responses Scale, PHQ-9, GAD-7) — on a common *z*-scored scale. We took the item pairs correlating at *r >* 0.7 across participants as psychometric synonyms (41 pairs). We then computed each participant’s index as the Pearson correlation between their responses to the first and to the second member of each pair, so that a participant who answers semantically near-identical items inconsistently scores low. We mean-imputed missing responses in the 38 additional items on the *z*-scored scale, which cannot inflate the item–item correlations that the pair-finding step thresholds. The 209 factor-analysis items carried the imputed values already produced for the factor analysis.

We *z*-scored both proxies across participants. We first regressed each psychopathology factor score on both proxies simultaneously in a linear model and tested the partial coefficients with two-tailed *t* tests, BH–FDR corrected over the six proxy × factor tests. We report the variance explained by each proxy on its own per cell as the *R*^2^ of the corresponding single-proxy regression (Supplementary Figure 7A). We then refit the main pooled trial-level choice model with the two proxies added to the moderator block, on the same trials and with participant-clustered standard errors as before. The moderator block (1 + AD + CI + SW) thereby became (1 + AD + CI + SW + completion time + cross-item reliability). We compared the two fits term by term over the 30 psychopathology coefficients — the five economic attribute regressors × three factors × {main, × context} (Supplementary Figure 7B). We summarized agreement between the original and the adjusted coefficient vectors by their Pearson correlation, by the mean absolute fractional change per coefficient, and by the through-origin slope of adjusted on original coefficients.

### 4.15 Task-comprehension adjustment

Participants answered a task-comprehension question (Q6) in each context. We scored a participant as passing only if the answer was correct in both contexts, counting a missing response as incorrect; of the 1,954 retained participants, 1,526 passed and 428 failed. This is the same variable that defines the comprehension filter of Supplementary Figure 17G. To ask whether the reported associations depend on task comprehension, we refit the pooled trial-level GLM — the analysis behind Figure 3C and Figure 4D — with this variable adjusted out: we added a *z*-scored pass indicator to the moderator block, so that (1 + AD + CI + SW) became (1 + AD + CI + SW + Q6), on the full n=1954 with cluster-robust standard errors and the identical 48-term global BH family as the published model.

We performed this analysis after the primary results and report it as a post-hoc, exploratory robustness check (Supplementary Figure 17H). We compare the adjusted and published coefficients over the 30 psychopathology-interaction terms — the five economic regressors crossed with the three factors, each entered as a factor interaction and as a context × factor interaction — on an identity plot, quantified by their Pearson correlation, the mean absolute fractional change in coefficient, and the least-squares slope through the origin. The comparison is drawn by the same routine as Supplementary Figure 7B, on which it is modeled.

We use the same variable once more, as a filter rather than as an adjustment. Supplementary Figure 17G redraws the individual Info-weight density scatter of Figure 6C on the 1,526 participants who passed, with subgroup membership again held frozen at its published definition, so that the 73 of the 130 members who passed are drawn over the density of the passing sample; the crosshair marks that sample’s mean and standard error in each context. No test is attached to this panel. It asks only whether the information-reflecting pattern is still present once the sample is restricted to participants who demonstrably understood the task, and it is read against the panel it repeats.

### 4.16 Visualization and shared statistical procedures

We computed two-dimensional density backgrounds (Figures 2C, 6C and Supplementary Figure 17G) with a product-Gaussian kernel density estimate on a 300 × 300 grid, with per-axis bandwidth set to 1.5× Silverman’s rule (1.06 *σ n*^-1^_/_^5^) and log-transformed for display. These densities are descriptive only; in subgroup panels, the maximum density attained over subgroup members set a display threshold determining which non-subgroup points were drawn individually. For correlation analyses (Supplementary Figures 1D, 3B, and 4) we used parametric Pearson or Spearman tests throughout. We used permutation tests only where stated above — the item-level classifier, the demographic comparisons, the information-seeking trend test, and the PLS component and omnibus tests; no correlation inference was permutation-based. Benjamini–Hochberg FDR corrections throughout used the standard step-up procedure (equivalent implementations in MATLAB, `scipy`, and `statsmodels`); we state the correction pool for each analysis with that analysis.

### 4.17 Software

We implemented pooled trial-level GLMs, mixed models, cluster-robust covariance estimation, the scaling and contextchange-axis analyses, and the cumulative prospect theory pipeline in Julia 1.11.4 with pinned package versions (`GLM.jl`, `MixedModels.jl`, `CovarianceMatrices.jl`, `ForwardDiff.jl`, `Optim.jl`). We implemented per-participant choice GLMs, model-free psychometric fits, the reward-uncertainty contingent information-seeking analyses, and the tertile-stratified analyses in MATLAB. We implemented the psychopathology factor analysis in R (`psych`, `GPArotation`, `polycor`, `glmnet`, `softImpute`). We implemented partial least squares regression, the L1-regularized item-level classifier, and the bootstrap and permutation procedures in Python 3.10 (`scikit-learn`, `statsmodels`, `scipy`, `numpy`).

### 4.18 Data and code availability

We will make all data and analysis code publicly available upon publication (see Declarations for repository details).

## Supplementary information

The Supplementary Information comprises Supplementary Figures 1–17 and Supplementary Table 1.

## Acknowledgements

The authors thank their affiliated institutions for support. Y.-Y.F. was supported by the Medical Scientist Training Program at Washington University in St. Louis. This work was supported by the National Institute of Mental Health under award numbers R01MH128344, R01MH110594, and R01MH116937; and by the Conte Center on the Neurocircuitry of OCD MH10643 (to I.E.M.). The decision-making paradigm used in this study evolved from a gain-only version of the task developed with Dr. Ethan S. Bromberg-Martin (see Bromberg-Martin and Feng, et al., 2024; Nature Neurosci). We would like to acknowledge Christopher Teng for his help with task code changes. We are grateful to members of the Laboratory of Adaptive and Maladaptive Intelligence for helpful discussions that improved this manuscript.

## Declarations

- Funding: NIH/NIMH R01MH128344, R01MH110594, R01MH116937, Conte Center MH10643, Wellcome Trust Discovery Award 554452/Z/25/Z (to I.E.M.); Medical Scientist Training Program (to Y.-Y.F.).
- Conflict of interest: The authors declare no competing interests.
- Ethics approval and consent to participate: All procedures were approved by the Washington University in St. Louis Institutional Review Board; participants provided informed consent.
- Consent for publication: Not applicable.
- Data availability: Data from this study are available upon reasonable request to the corresponding author and will be made publicly available upon publication at https://github.com/yang-yangfeng/gainlosspaper.
- Materials availability: Not applicable.
- Code availability: Code from this study is available upon reasonable request to the corresponding author and will be made publicly available upon publication at https://github.com/yang-yangfeng/gainlosspaper.
- Author contribution: Y.-Y.F. led conceptualization, methodology, data collection, analysis, and writing of the manuscript. I.E.M. oversaw and contributed to conceptualization, analysis, funding acquisition, and writing of the manuscript.
- AI usage: AI tools were used for feedback on manuscript drafts and refactoring code. All intellectual contributions are made by the authors who take full responsibility for the statements made herein.

---

**Supplementary Figure 1.**
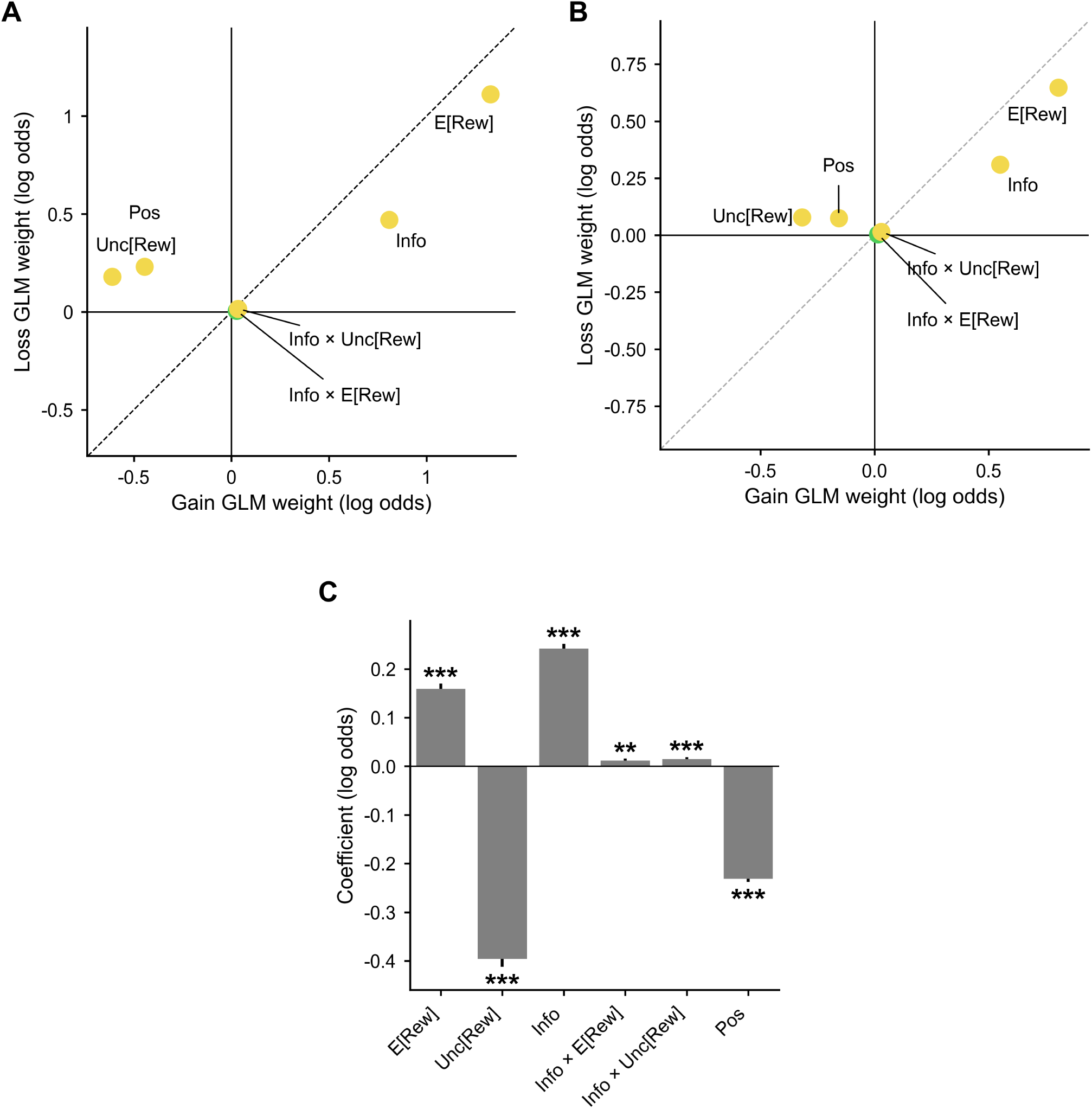
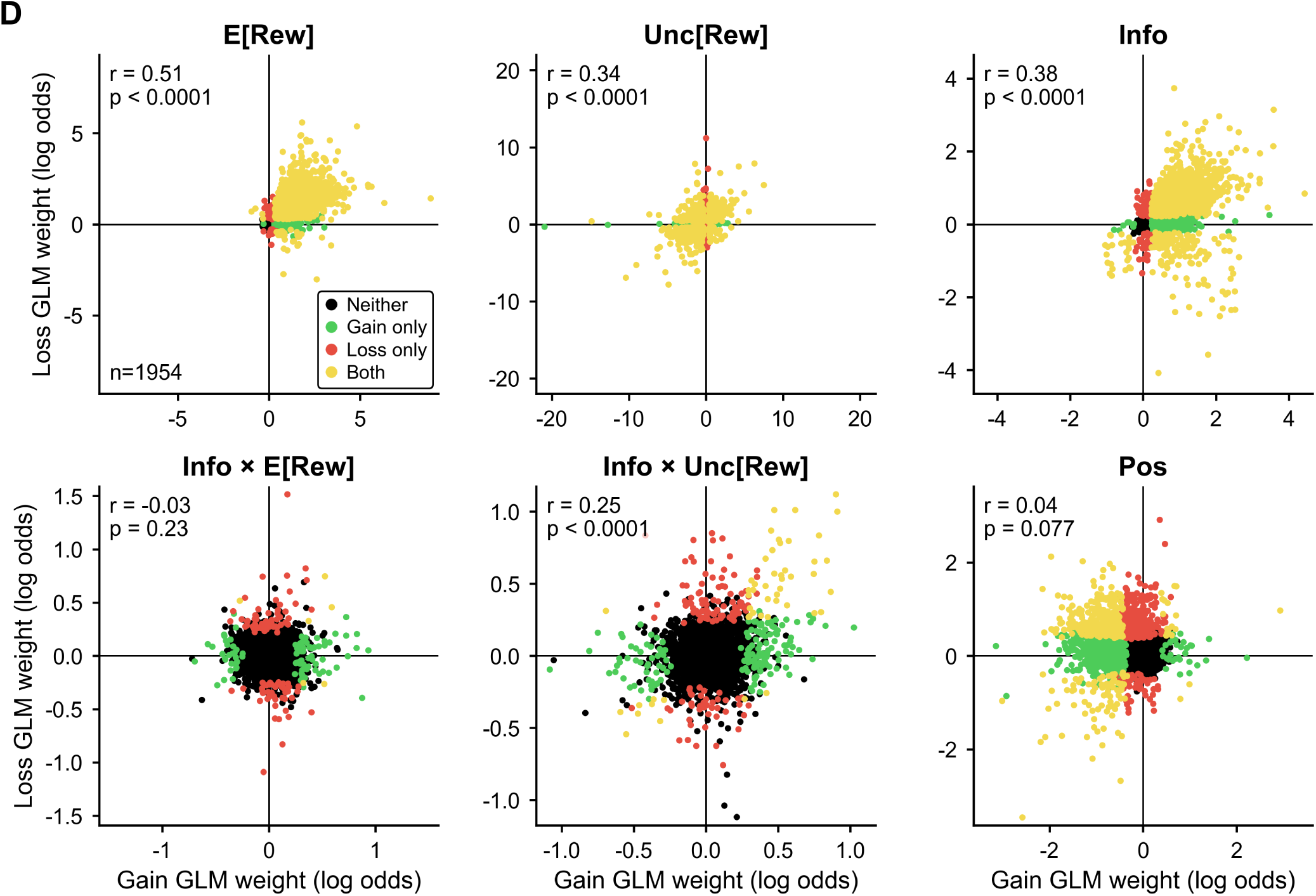
Population and individual economic preferences in gain and loss, across all six choice-GLM regressors. The choice model of **Figure 2** carries six regressors—expected reward (E[Rew]), reward uncertainty (Unc[Rew]), information (Info), the two interactions Info × E[Rew] and Info × Unc[Rew], and the side-bias term Pos (Methods). **Figure 2** displays three of them; this figure displays all six, at both levels of analysis—averaged over the n=1954 individual fits **(A)** and from one population fit **(B)**, with their gain–loss differences **(C)** and the full individual weight distributions **(D)**. Throughout, the loss-context E[Rew] and Info × E[Rew] measures are sign-flipped (scored on the higher-reward, i.e. less negative, offer) so that a positive weight means reward-seeking in both contexts; Unc[Rew], Info, Info × Unc[Rew] and Pos are not flipped. *, **, *** indicate *p <* 0.05, 0.01, 0.001. **(A)** Population average of the individual GLM weights: one labeled point per regressor at the mean across individuals in gain (x-axis *Gain GLM weight (log odds)*) vs. loss (y-axis *Loss GLM weight (log odds)*), with SE bars (smaller than the markers) and a black dashed unity line; solid lines mark zero. Marker color is a one-sample *t*-test of that weight against zero, significant in gain (green), loss (red), both (yellow) or neither (black), BH–FDR corrected over the panel’s 12 within-context tests (6 regressors × 2 contexts); marker fill encodes the separate gain-minus-loss test, BH-corrected over its own 6, so an unfilled marker would denote a non-significant context difference. All six markers are filled and five are yellow: E[Rew] +1.327 ± 0.019 in gain and +1.111 ± 0.019 in loss; Info +0.808 ± 0.014 and +0.470 ± 0.015; Info × Unc[Rew] +0.034 ± 0.005 and +0.016 ± 0.004; Unc[Rew] reversing from −0.610 ± 0.032 to +0.180 ± 0.028; Pos reversing from −0.444 ± 0.012 to +0.231 ± 0.012 (all *p <* 0.0001 except Info × Unc[Rew] in loss, *p*_BH_ = 3.4 × 10^-4^). Info × E[Rew] is the exception and is drawn green: positive in gain (+0.027 ± 0.004, *p <* 0.0001) but indistinguishable from zero in loss (+0.005 ± 0.004, *p* = 0.17). **(B)** The same six regressors from the population (cluster-robust) choice GLM, decomposed into context-specific coefficients (*β*_gain_ = *β*_main_ + 0.5 · *β*_context_, *β*_loss_ = *β*_main_ - 0.5 · *β*_context_) and plotted on the same axes with cluster-robust SE arms and a gray dashed unity line. Point color is a two-tailed Wald test against zero on the same four-class scheme, BH– FDR corrected over this model’s 48 context-decomposed effects (24 model terms × 2 contexts; Methods). The pattern matches **(A)**: E[Rew] +0.806 ± 0.011 in gain and +0.647 ± 0.011 in loss; Info +0.551 ± 0.009 and +0.309 ± 0.010; Info × Unc[Rew] +0.0299 ± 0.0034 and +0.0151 ± 0.0030; Unc[Rew] reversing from −0.318 ± 0.014 to +0.078 ± 0.013; Pos reversing from −0.157 ± 0.004 to +0.074 ± 0.004; all of these *p*_BH_ *<* 0.0001. Info × E[Rew] is again green—positive in gain (+0.0152 ± 0.0028, *p*_BH_ *<* 0.0001) and not in loss (+0.0036 ± 0.0026, *p*_BH_ = 0.26). Directly contrasting the two interactions within each context (difference of the decomposed coefficients, cluster-robust Wald test on the linear contrast; Methods), the Info × Unc[Rew] weight exceeds the Info × E[Rew] weight in both contexts: gain Δ = +0.0147 ± 0.0045, *z* = 3.24, *p*_BH_ = 2.4 × 10^-3^; loss Δ = +0.0115 ± 0.0039, *z* = 2.92, *p*_BH_ = 3.5 × 10^-3^ (BH over the two contrasts). **(C)** Gain-minus-loss differences in the population GLM, estimated as each regressor’s interaction with context under the ±0.5 context coding: gray bars with black SE bars on the y axis *Coefficient (log odds)*, with the six regressors as rotated x tick labels. Stars are two-tailed cluster-robust Wald tests from the same 48-effect BH family. All six context interactions are significant: E[Rew] +0.159 ± 0.011 (*z* = 14.66), Unc[Rew] −0.396 ± 0.016 (*z* = −24.76), Info +0.241 ± 0.010 (*z* = 23.49) and Pos −0.232 ± 0.006 (*z* = −38.57), all *p*_BH_ *<* 0.0001; Info × E[Rew] +0.0116 ± 0.0039 (*z* = 2.97, *p*_BH_ = 7.2 × 10^-3^) and Info × Unc[Rew] +0.0148 ± 0.0039 (*z* = 3.75, *p*_BH_ = 5.2 × 10^-4^). **(D)** The individual variability behind the averages in **(A)**: one tile per regressor showing all n=1954 individual GLM weights in gain (x) vs. loss (y), each tile on its own scale with its own tick labels and with solid lines at zero. Each participant is colored by the significance of their own two weights (*p <* 0.05 per context, from the individual fits, uncorrected): significant in gain only (green), loss only (red), both (yellow) or neither (black). The legend is drawn once, in the E[Rew] tile. The E[Rew] tile is annotated with the sample size (n=1954), and every tile with the Pearson correlation between the two contexts’ weights and its parametric two-tailed p-value: E[Rew] *r* = 0.51, Unc[Rew] *r* = 0.34, Info *r* = 0.38 and Info × Unc[Rew] *r* = 0.25 (all *p <* 0.0001), whereas Info × E[Rew] (*r* = −0.03, *p* = 0.23) and Pos (*r* = 0.04, *p* = 0.077) are not correlated across contexts. As in **Figure 2C**, individual variation in the Info × Unc[Rew] weight exceeded estimation noise in both contexts (SD_obs_, SD_noise_, *I*^2^, Cochran’s *Q* as defined there; Methods): gain 0.21, 0.17, 34%, *Q*(1953) = 2978 and loss 0.20, 0.15, 31%, *Q* = 2822, both *p <* 0.0001; the per-participant gain-loss difference in this weight was likewise heterogeneous, as in **Figure 2E** (0.25, 0.23, 12%, *Q*(1953) = 2228, *p* = 1.2 × 10^-5^). The joint sign structure of these clouds is tabulated in **Supplementary Figure 3A** and their cross-regressor correlations in **Supplementary Figure 3B**.

**Supplementary Figure 2.**
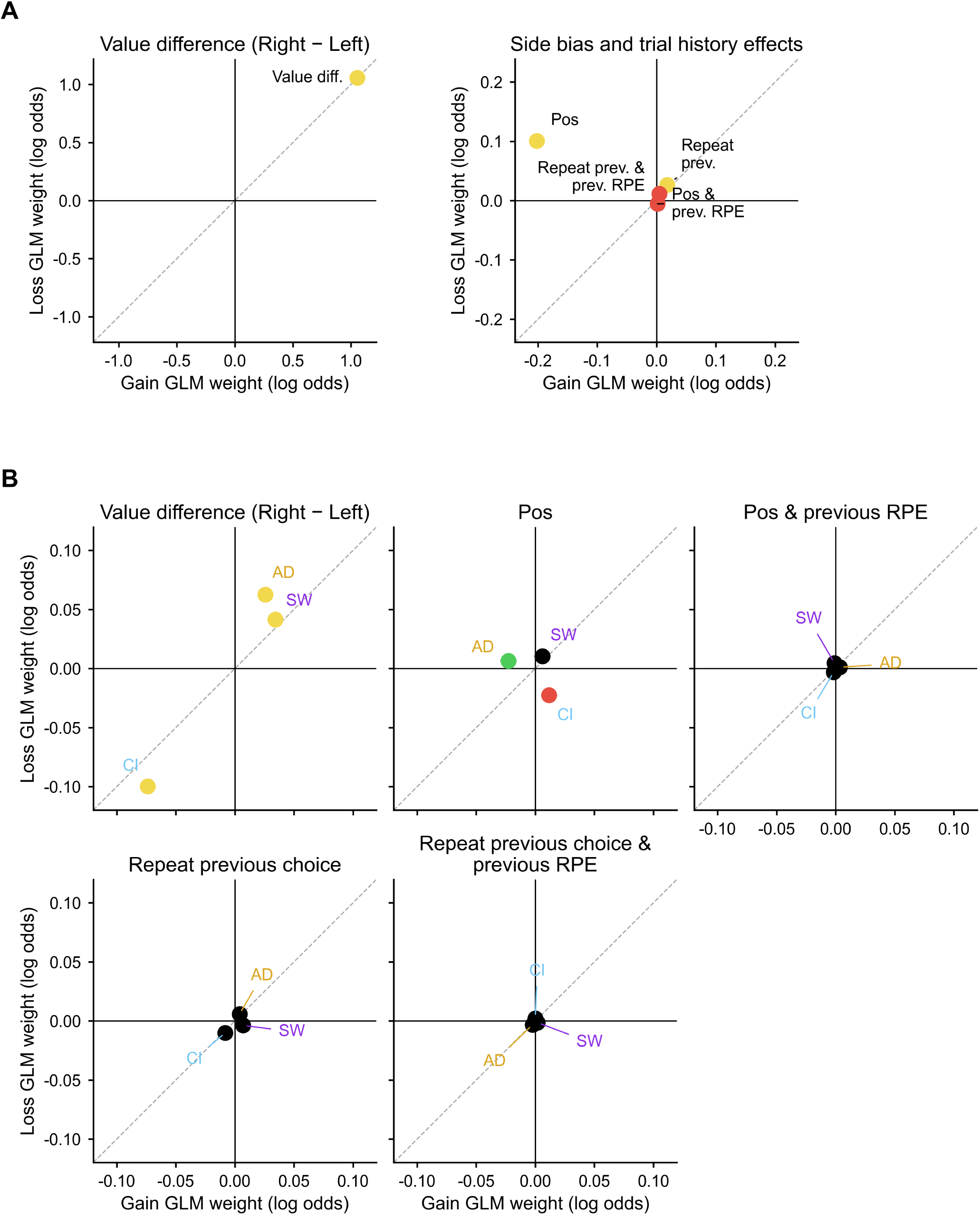
Choices are driven by economic value, not by trial history. A population (cluster-robust) logistic GLM of choice on the value difference between the two offers (computed from each participant’s own economic preference weights), the Pos side bias, the modulation of that side bias by the previous trial’s reward prediction error (RPE), the tendency to repeat the previous choice, and the modulation of that tendency by the previous trial’s RPE; fitted across n=1954 participants and decomposed into gain- and loss-context coefficients (Methods). Both panels plot coefficients in gain (x-axis *Gain GLM weight (log odds)*) vs. loss (y-axis *Loss GLM weight (log odds)*) with cluster-robust SE arms (smaller than the markers for the largest terms), a gray dashed unity line and solid lines at zero; point color is a two-tailed Wald test against zero that is significant in gain (green), loss (red), both (yellow) or neither (black), BH–FDR corrected over this model’s 40 context-decomposed effects (20 model terms × 2 contexts). **(A)** Population effects, split across two tiles because of their scale. Left (*Value difference (Right* - *Left)*): the value difference dominates choice and is essentially identical across contexts (+1.054 ± 0.003 in gain, +1.057 ± 0.003 in loss; both *p <* 0.0001). Right (*Side bias and trial history effects*), on a several-fold finer axis: the Pos side bias reverses between contexts (−0.202 ± 0.005 vs. +0.101 ± 0.005; both *p <* 0.0001); the tendency to repeat the previous choice is small but significant in both (+0.018 ± 0.004, *p*_BH_ *<* 0.0001; +0.026 ± 0.004, *p*_BH_ *<* 0.0001); and the two RPE terms are smaller still and reach significance in the loss context only, so their points are red—the RPE modulation of the side bias (+0.002 ± 0.003, n.s.; −0.006 ± 0.002, *p*_BH_ = 0.045) and of the repeat tendency (+0.004 ± 0.003, n.s.; +0.012 ± 0.002, *p*_BH_ *<* 0.0001). Choice therefore depends overwhelmingly on the economic attributes, with only a weak tendency to repeat the previous choice and weak dependence on the previous trial’s RPE. **(B)** Interactions of the same five effects with AD, CI and SW, one tile per effect (*Value difference (Right* - *Left)*, *Pos*, *Pos & previous RPE*, *Repeat previous choice*, *Repeat previous choice & previous RPE*), with the three points labeled by factor. Sensitivity to the overall value difference mirrors the E[Rew] pattern of **Figure 3C**: AD and SW interact positively in both contexts (AD +0.026 gain, +0.063 loss; SW +0.034, +0.041) and CI negatively in both (−0.074, −0.100), all six *p*_BH_ *<* 0.0001. For the side bias, AD interacts negatively in gain only (−0.023, *p*_BH_ = 3.2 × 10^-3^) and CI negatively in loss only (−0.023, *p*_BH_ = 1.0 × 10^-3^), while SW does not differ in either context. No factor interacts significantly with any of the three trial-history terms—all nine of those points are black, the nearest being CI with the repeat tendency in loss (*p*_BH_ = 0.12)—so the psychopathology-related differences in this task act on economic value rather than on history-driven choice.

**Supplementary Figure 3.**
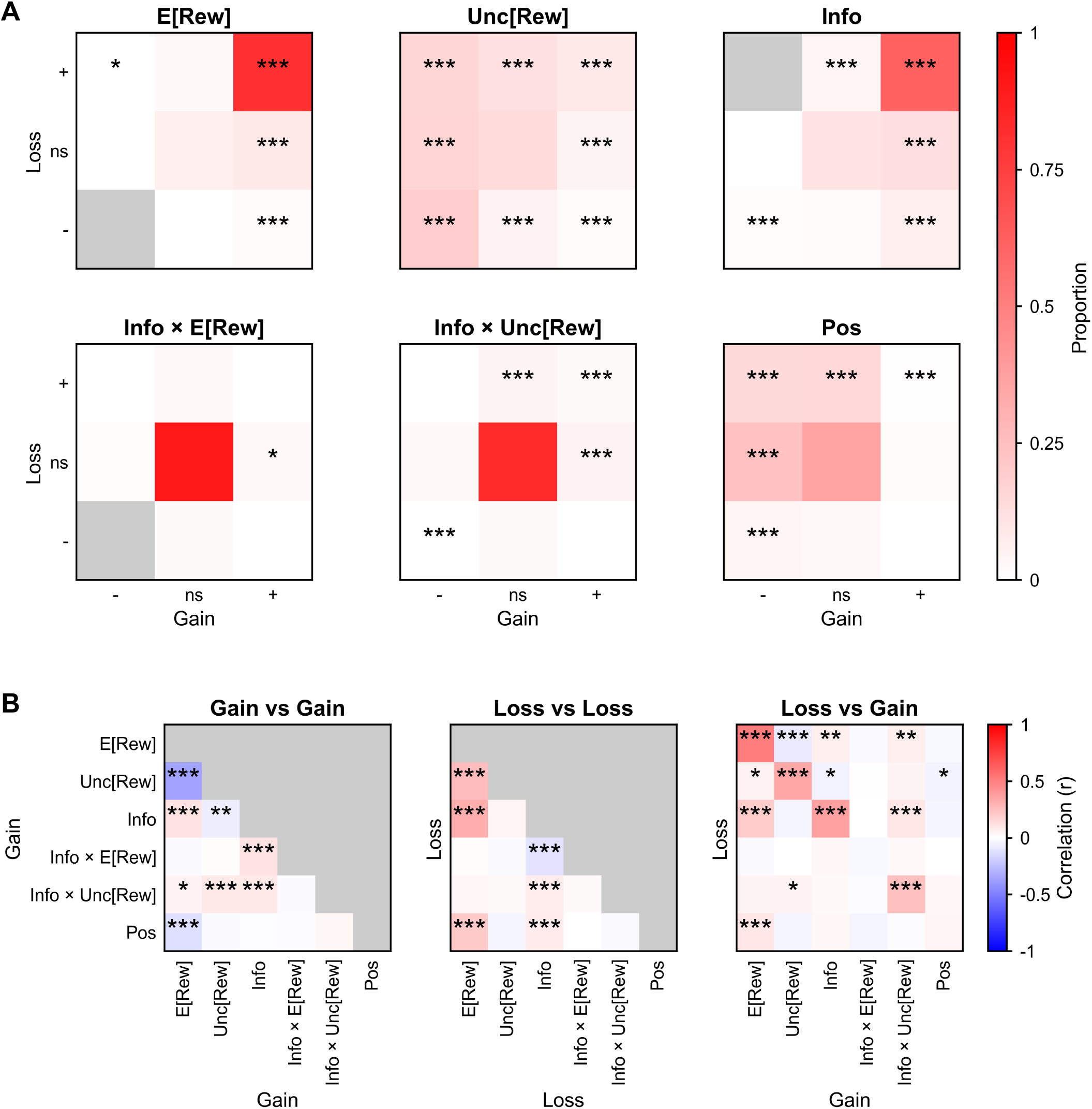
Structure of variability in economic preferences across gain and loss. Two views of how the n=1954 individual GLM weights plotted in Supplementary Figure 1D are jointly organized: their sign structure across contexts **(A)** and their correlation structure across regressors **(B)**. In both panels the loss-context E[Rew] and Info × E[Rew] weights are sign-flipped as in Supplementary Figure 1, and *, **, *** indicate *p <* 0.05, 0.01, 0.001. **(A)** Contingency of gain- and loss-context weight signs, one 3 × 3 heat map per regressor (E[Rew], Unc[Rew], Info, Info × E[Rew], Info × Unc[Rew], Pos). Every participant is classified by the sign and significance of their gain weight (columns, left to right: -, ns, +) and of their loss weight (rows, top to bottom: +, ns, -), taking *p <* 0.05 per weight, and each cell is shaded by the proportion of the 1954 participants it holds, on the shared color bar *Proportion* (white to red, 0 to 1). Three cells across the six maps hold no participants at all—the doubly negative cell of E[Rew] and of Info × E[Rew], and the gain-negative/loss-positive cell of Info—and are drawn gray rather than white. Stars are a one-tailed binomial enrichment test of each cell against the null that the two contexts’ signs are independent (the cell’s null probability is the product of the two marginal chance rates, *α/*2 for each significant sign and 1 - *α* for ns), with raw uncorrected p-values; stars therefore mark over-represented cells only, and an unstarred cell is simply one that is not more abundant than chance rather than one tested for under-representation. Context agreement dominates for E[Rew] (80.7% of participants significantly positive in both contexts) and for Info (62.4% positive in both; both *p <* 0.0001), whereas Unc[Rew] spreads across the whole table with all eight non-central cells enriched (all *p <* 0.0001) and the two Info interactions place most participants in the doubly non-significant center cell (90.1% for Info × E[Rew], 83.0% for Info × Unc[Rew]). The Info map also contains the asymmetry the subpopulation of **Figure 6C** is built on: 130 participants (6.7%) are significantly positive in gain and significantly negative in loss (*p <* 0.0001), while the mirror-image cell is empty. For Info × Unc[Rew], the cell of participants significantly negative in both contexts holds 8 participants (*p <* 0.0001). **(B)** Pearson correlations among the six individual GLM weights, as three 6 × 6 heat maps on the shared color bar *Correlation (r)* (blue–white–red, −1 to +1): *Gain vs Gain* and *Loss vs Loss*, each within one context and drawn as a lower triangle with the diagonal and upper triangle masked gray, and *Loss vs Gain*, all 36 across-context pairs with rows Loss and columns Gain. The six regressors label both axes in the same order — row labels drawn once, on the lefthand map, column labels beneath each map — and each map’s own axis words carry the context. Stars are parametric two-tailed tests of each correlation, raw and uncorrected. Within gain, the E[Rew] and Unc[Rew] weights are strongly negatively correlated (*r* = −0.36) whereas within loss the same pair is positively correlated (*r* = +0.26), mirroring the reversal of risk preference between contexts, and Info correlates positively with E[Rew] in both contexts (*r* = +0.12 in gain, +0.32 in loss; all four *p <* 0.0001). The across-context map is dominated by its diagonal—each regressor’s own gain-versus-loss correlation, the values annotated on the tiles of **Supplementary Figure 1D**—with the largest off-diagonal entry linking loss Info to gain E[Rew] (*r* = +0.20) and a small negative entry linking loss E[Rew] to gain Unc[Rew] (*r* = −0.077; both *p <* 0.001).

**Supplementary Figure 4.**
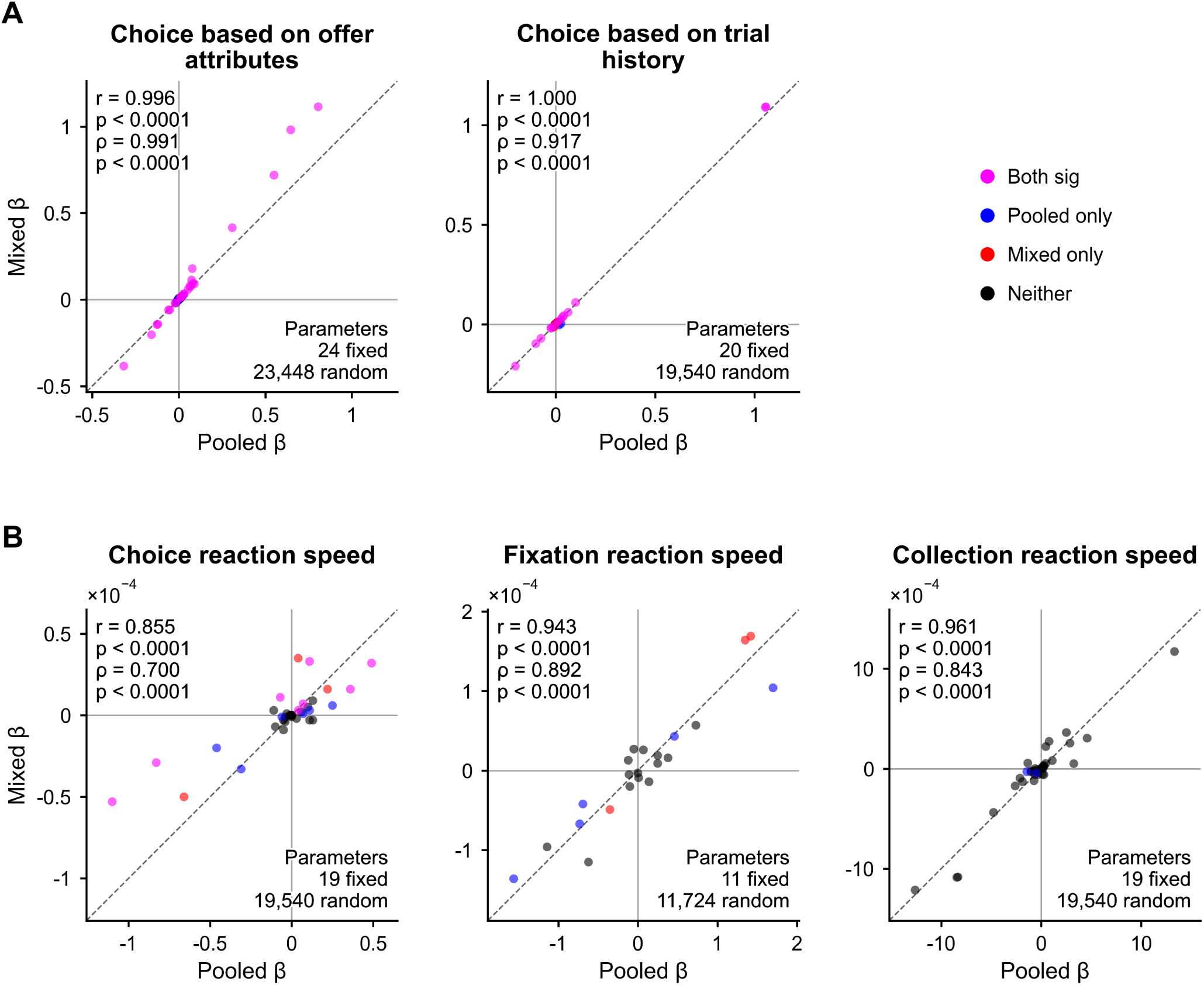
Fixed-effect estimates agree between the pooled population models and mixed models with individual random effects. Every population model in this paper is fit as a pooled GLM with cluster-robust standard errors. This figure checks each of those models against the corresponding mixed model, in which every participant additionally receives random effects. Each cell plots one model’s context-decomposed fixed effects — the pooled estimate (x-axis *Pooled β*) against the mixed-model estimate (y-axis *Mixed β*), one point per model term per context — with a gray dashed identity line and a faint origin crosshair. The annotation at the upper left of each cell gives the Pearson *r* and the Spearman *ρ* between the two sets of estimates with their parametric two-tailed p-values, all of which are drawn at the panel’s floor of *p <* 0.0001; the annotation at the lower right gives how many fixed and random parameters the mixed model carries. Point color records whether the term was significant in both models (magenta), in the pooled model only (blue), in the mixed model only (red), or in neither (black); the legend is drawn once, beside **(A)**. **(A)** The two choice models. Left, *Choice based on offer attributes* — the model of **Figure 2** and **Supplementary Figure 1**, 48 context-decomposed effects from 24 fixed parameters against 23,448 random effects — with *r* = 0.996 and *ρ* = 0.991. Right, *Choice based on trial history* — the model of **Supplementary Figure 2**, 40 effects from 20 fixed parameters against 19,540 random effects — with *r* = 1.000 and *ρ* = 0.917. Both clouds sit on the identity line and are almost entirely magenta, so no choice-based conclusion in this paper depends on the pooled specification. **(B)** The three reaction-speed models, same conventions, with a common ×10^-4^ factor taken out of the axes so that no cell needs sub-decimal ticks. *Choice reaction speed* (38 effects; 19 fixed, 19,540 random) is the loosest of the five, *r* = 0.855, *ρ* = 0.700, and carries most of the figure’s disagreement-colored points; because it is also the most heavily parameterized of the three reaction-speed models, part of that scatter is plausibly fitting unreliability on the mixed side rather than a substantive discrepancy. *Fixation reaction speed* (22 effects; 11 fixed, 11,724 random) gives *r* = 0.943, *ρ* = 0.892, and *Collection reaction speed* (38 effects; 19 fixed, 19,540 random) gives *r* = 0.961, *ρ* = 0.843 — both close agreements. All five Pearson and Spearman correlations are positive and highly significant.

**Supplementary Figure 5.**
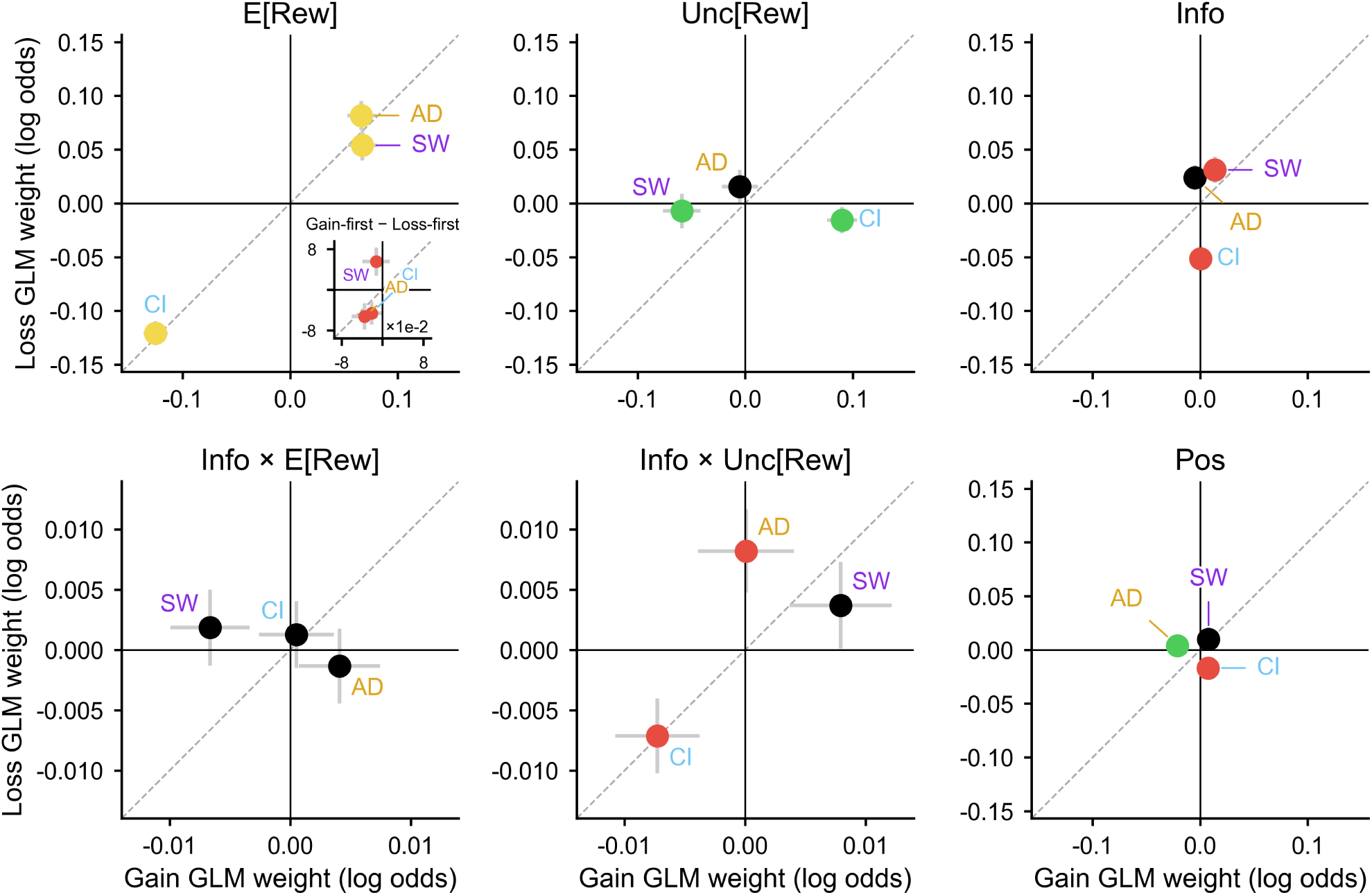
Population-GLM psychopathology-factor interactions, organized by economic attribute. The population (cluster-robust) factor interactions of **Figure 3C** and **Figure 4D**, from the same model, re-organized as one tile per economic regressor rather than one panel per psychopathology factor, and extended to all six regressors (E[Rew], Unc[Rew], Info, Info × E[Rew], Info × Unc[Rew] and the side-bias term Pos; as in **Supplementary Figure 1**). Each tile plots that regressor’s interaction with AD (gold), CI (light blue) and SW (violet) in gain (x-axis *Gain GLM weight (log odds)*) vs. loss (y-axis *Loss GLM weight (log odds)*), with gray cluster-robust SE arms, a gray dashed unity line and solid lines at zero; the three points are labeled by factor. Point color is a two-tailed Wald test against zero, significant in gain (green), loss (red), both (yellow) or neither (black), BH–FDR corrected over the model’s 48 context-decomposed effects (24 model terms × 2 contexts)—a broader family than the 12-test attribute × context × factor family that governs the arrows in **Figure 3C**. Because the main-effect and Info-interaction regressors differ by roughly two orders of magnitude in coefficient scale, the two groups of tiles use separate axis ranges. The E[Rew] tile draws the three E[Rew] × factor interactions of **Figure 3C** themselves (all three points yellow) and carries the task-order check described below as a miniature inset in its lower-right. Of the remaining five, Unc[Rew] and Info recapitulate **Figure 3C**: for Unc[Rew], CI interacts positively in gain (+0.090, *p*_BH_ *<* 0.0001) and not in loss, SW negatively in gain (−0.059, *p*_BH_ = 1.5 × 10^-3^) and not in loss (both points green), and AD in neither (black); for Info, CI interacts negatively in loss (−0.051, *p*_BH_ *<* 0.0001) and SW positively in loss (+0.031, *p*_BH_ = 0.023), both points red, while AD is black in both contexts (nearest *p*_BH_ = 0.073, loss). Among the three regressors not shown in the main figures, Info × E[Rew] has no significant factor association in either context (all three points black; nearest SW in gain, *p*_BH_ = 0.073); Info × Unc[Rew] is significantly positive for AD and negative for CI in loss (+0.0082, *p*_BH_ = 0.034; −0.0071, *p*_BH_ = 0.041)—the same two effects displayed in **Figure 4D**—with SW black; and the Pos side bias is significantly negative for AD in gain and for CI in loss (−0.021, *p*_BH_ = 5.7 × 10^-4^; −0.017, *p*_BH_ = 2.0 × 10^-3^), with SW black. *Inset (lower-right of the E[Rew] tile, titled Gain-first* - *Loss-first, drawn on* ×10^-2^ *axes):* task-order robustness check for the E[Rew] × factor interactions. The gain-first and loss-first cohorts were fitted as separate submodels and their E[Rew] × factor coefficients differenced within the gain context (x-axis) and within the loss context (y-axis), one point per factor with independent-variance SE arms and a gray dashed unity line; p-values are raw and uncorrected (Methods). This is the same check drawn as an inset in **Figure 3C**. All three differences are small relative to the interactions themselves (|Δ*β*| ≤ 0.055): none differs from zero in the gain context (AD −0.035, *p* = 0.17; CI −0.021, *p* = 0.34; SW −0.012, *p* = 0.64), whereas all three are nominally significant in the loss context, which is why all three points are red (AD −0.053, *p* = 0.044; CI −0.046, *p* = 0.038; SW +0.055, *p* = 0.047).

**Supplementary Figure 6.**
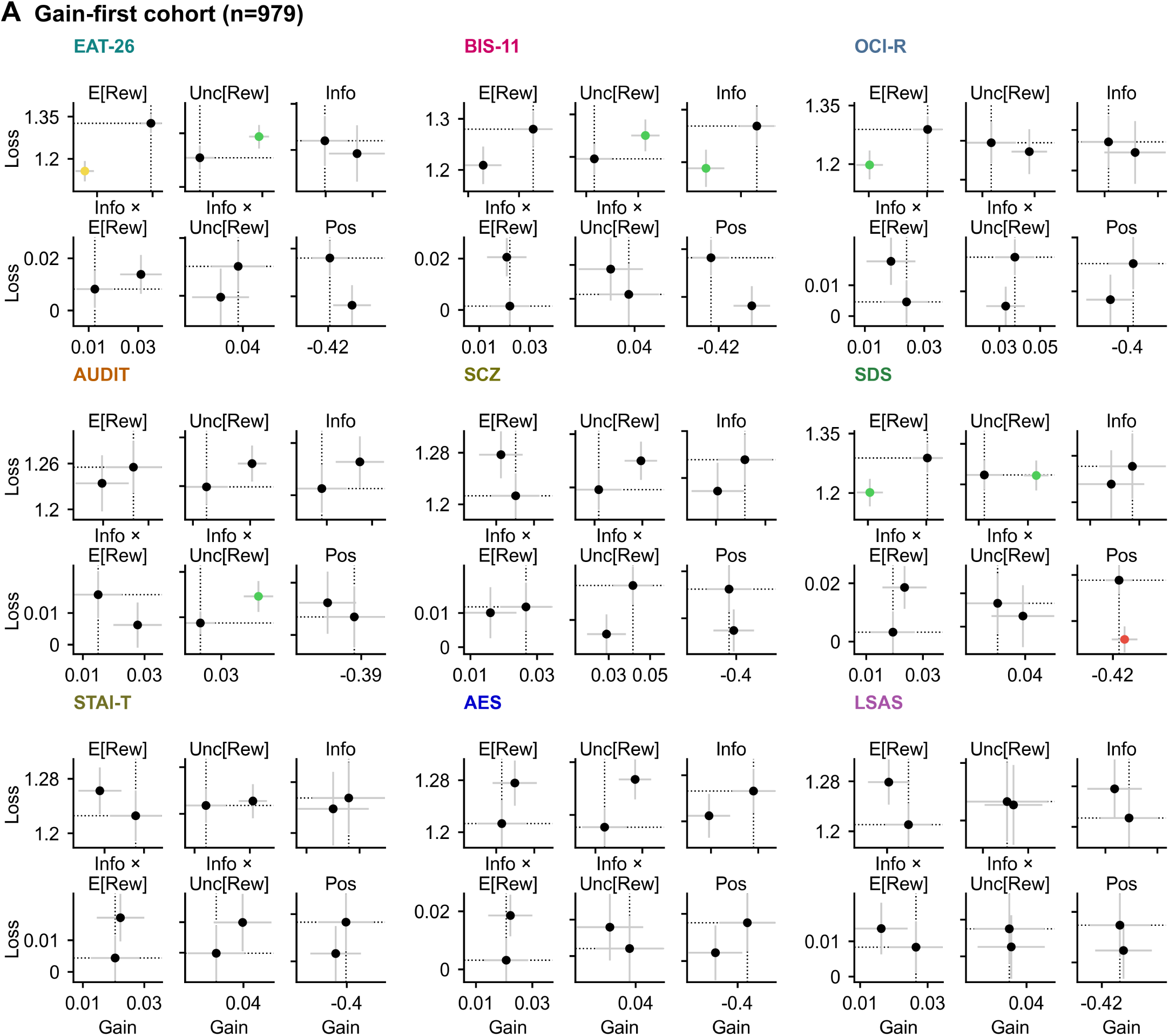
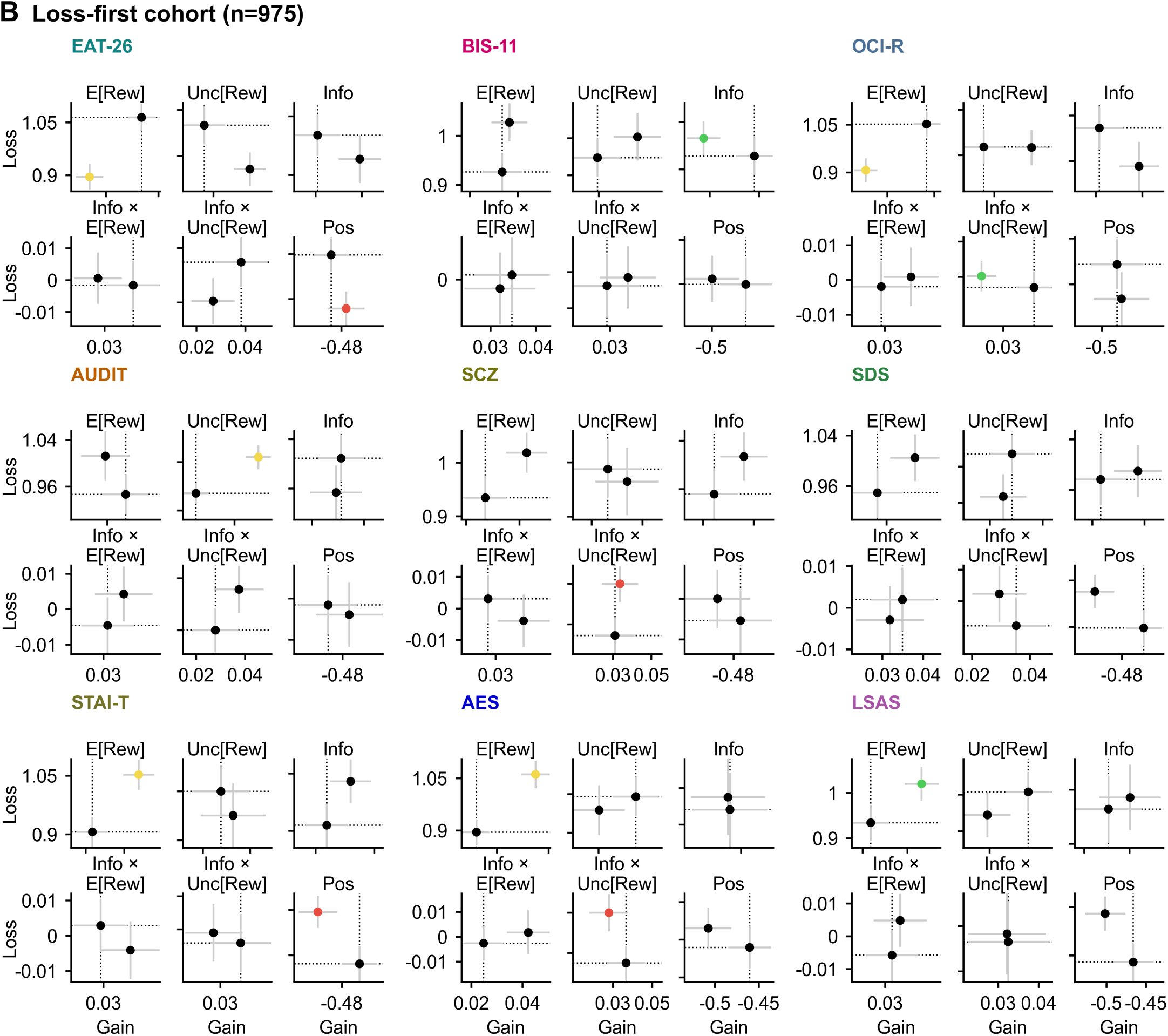
Dissociable preference alterations replicate across individual psychiatric symptom questionnaires and task order. Average economic preferences, measured by individual GLM weights, for participants scoring high vs. low on each of the nine psychiatric symptom questionnaires, shown separately for the two task-order cohorts: **(A)** the gain-first cohort (n=979) and **(B)** the loss-first cohort (n=975). Groups were defined by a median split on the questionnaire’s mean item *z*-score across the full n=1954 sample, then intersected with the task-order cohort (455–520 participants per group; Methods). Each panel is a 3 × 3 arrangement of nine questionnaire blocks, titled by instrument abbreviation in that instrument’s **Figure 1B** color (EAT-26, BIS-11, OCI-R, AUDIT, SCZ, SDS, STAI-T, AES, LSAS), and each block is a compact 2 × 3 grid of per-regressor tiles titled above each tile (E[Rew], Unc[Rew], Info, Info × E[Rew], Info × Unc[Rew], Pos). A questionnaire occupies the same grid position in **(A)** and **(B)**, so its two cohorts are compared by position across the two pages. Within a tile the two points are that questionnaire’s high- and low-scoring groups, plotted as the group’s average individual GLM weight in gain (x-axis) vs. loss (y-axis) in log odds, with SE arms; the dotted reference lines pass through the low group’s point, which is how the two groups are told apart in this compact format (point labels and group n’s are omitted for space). Every tile has its own axis limits, so the tick labels describe their own tile only, and the axis words Gain and Loss are drawn on each panel’s edge blocks. Color of the high group’s point marks whether it differs from the low group in gain (green), loss (red), both (yellow) or neither (black), by a two-tailed two-sample *t*-test on the two group means and their standard errors (uncorrected; Methods); the low group is the reference of that test and is therefore always black. The high-versus-low displacements broadly follow the factor-level associations of **Supplementary Figure 5** and **Figure 3**: the CI-loaded instruments (OCI-R, EAT-26) show the reward-sensitivity reduction in both cohorts, while the AD-loaded instruments (AES, STAI-T) show the corresponding elevation most clearly in the loss-first cohort, with SDS and BIS-11 mixed—consistent with raw questionnaire scores mixing variance from factors with opposing associations. The overall pattern indicates that the factor associations reported in the main text are carried by no single questionnaire and are not an artifact of which context participants encountered first.

**Supplementary Figure 7.**
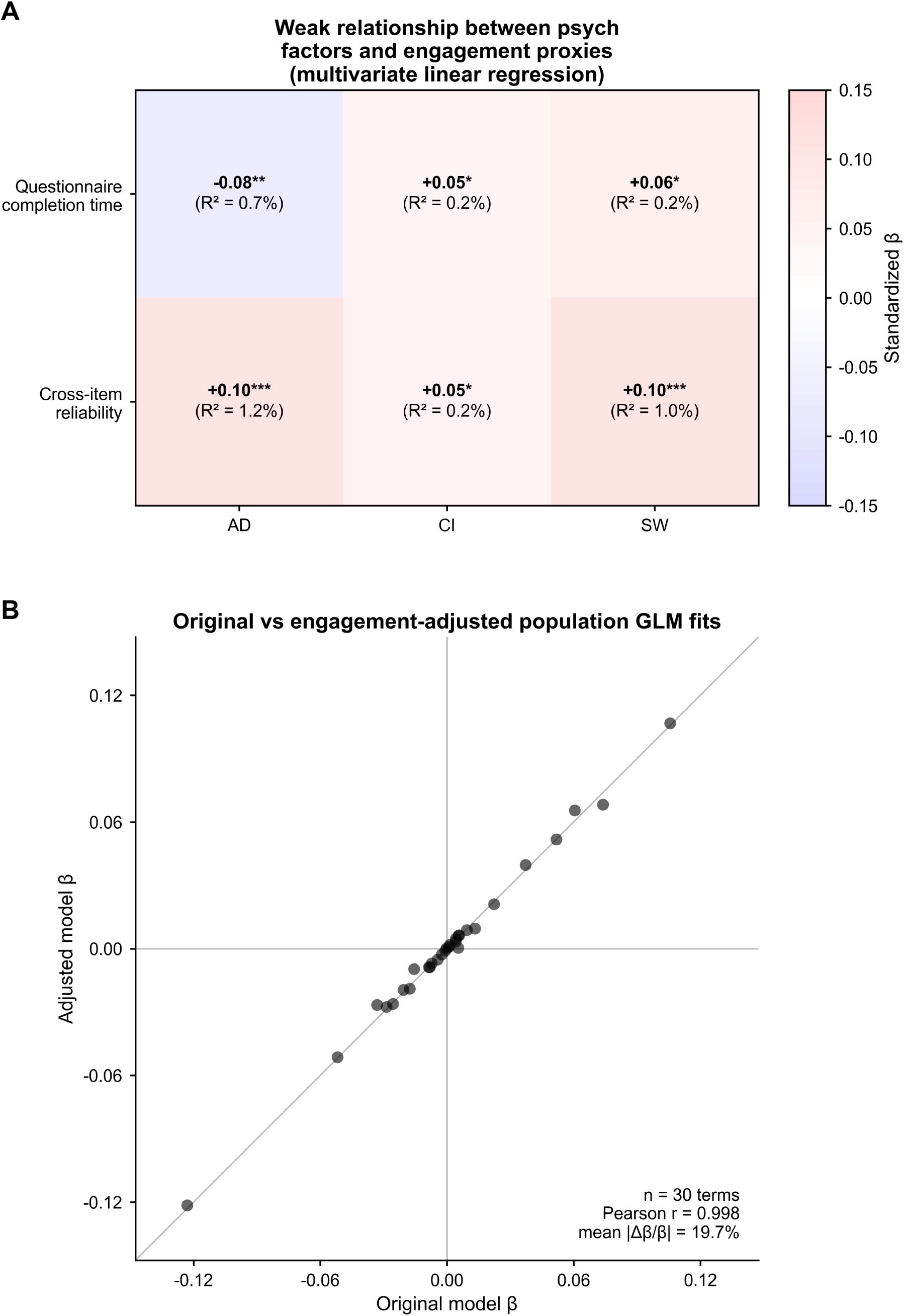
Economic preference associations with psychopathology are robust to controlling for questionnaire engagement. Control analyses against careless or disengaged questionnaire responding, using two engagement proxies computed for the full n=1954 sample: total questionnaire completion time, and a cross-item reliability score built from 41 psychometric-synonym item pairs, which flags respondents who answer semantically near-identical items inconsistently (the psychometric-synonym index of Meade & Craig, 2012; Methods). **(A)** Weak relationship between the psychopathology factors and the engagement proxies (panel subtitle *Weak relationship between psych factors and engagement proxies (multivariate linear regression)*): a 2 × 3 heat map with the two proxies as rows (*Questionnaire completion time*, *Cross-item reliability*) and AD, CI and SW as columns, shaded by the standardized partial regression coefficient on the color bar *Standardized β* (blue–white–red, ±0.15). Each factor score was regressed on both z-scored proxies simultaneously; stars are two-tailed *t*-tests of the partial coefficients, BH–FDR corrected over the 6 proxy × factor tests. Each cell prints its own coefficient with its star tier and, beneath it, the *R*^2^ of the simple regression of that factor on that proxy alone. Questionnaire completion time was weakly negatively associated with AD (−0.08, *p*_BH_ = 1.2 × 10^-3^) and weakly positively associated with CI and SW (+0.05, *p*_BH_ = 0.039; +0.06, *p*_BH_ = 0.019); cross-item reliability was positively associated with all three, most strongly with AD and SW (+0.10, *p*_BH_ *<* 0.0001; +0.05, *p*_BH_ = 0.026; +0.10, *p*_BH_ *<* 0.0001). The in-cell *R*^2^ values show that neither proxy accounts for more than 1.2% of the variance in any factor. **(B)** Original vs. engagement-adjusted population GLM fits (panel subtitle as drawn): each point is one of the 30 psychopathology terms of the population economic-choice GLM—the E[Rew], Unc[Rew], Info, Info × E[Rew] and Info × Unc[Rew] interactions with AD, CI and SW, and each of those terms’ own interaction with context—plotted as its coefficient in the original model (x-axis *Original model β*) against its coefficient in the engagement-adjusted model, which adds both proxies as covariates (y-axis *Adjusted model β*); the gray line is unity and solid lines mark zero. The in-panel annotation gives *n* = 30 terms, Pearson *r* = 0.998 and mean |Δ*β/β*| = 19.7%; the through-origin slope of adjusted on original coefficients, 0.988, is not drawn on the figure. The two coefficient vectors are thus almost perfectly collinear, and 29 of the 30 terms keep their sign—the exception, the Info × Unc[Rew] × context × CI term, moves from −0.0001 to +0.0001 and is indistinguishable from zero in both models (*p >* 0.97)—while the largest fractional shifts fall on the smallest coefficients. The psychopathology-factor associations with economic preferences reported throughout the paper are therefore not attributable to careless or disengaged questionnaire responding.

**Supplementary Figure 8.**
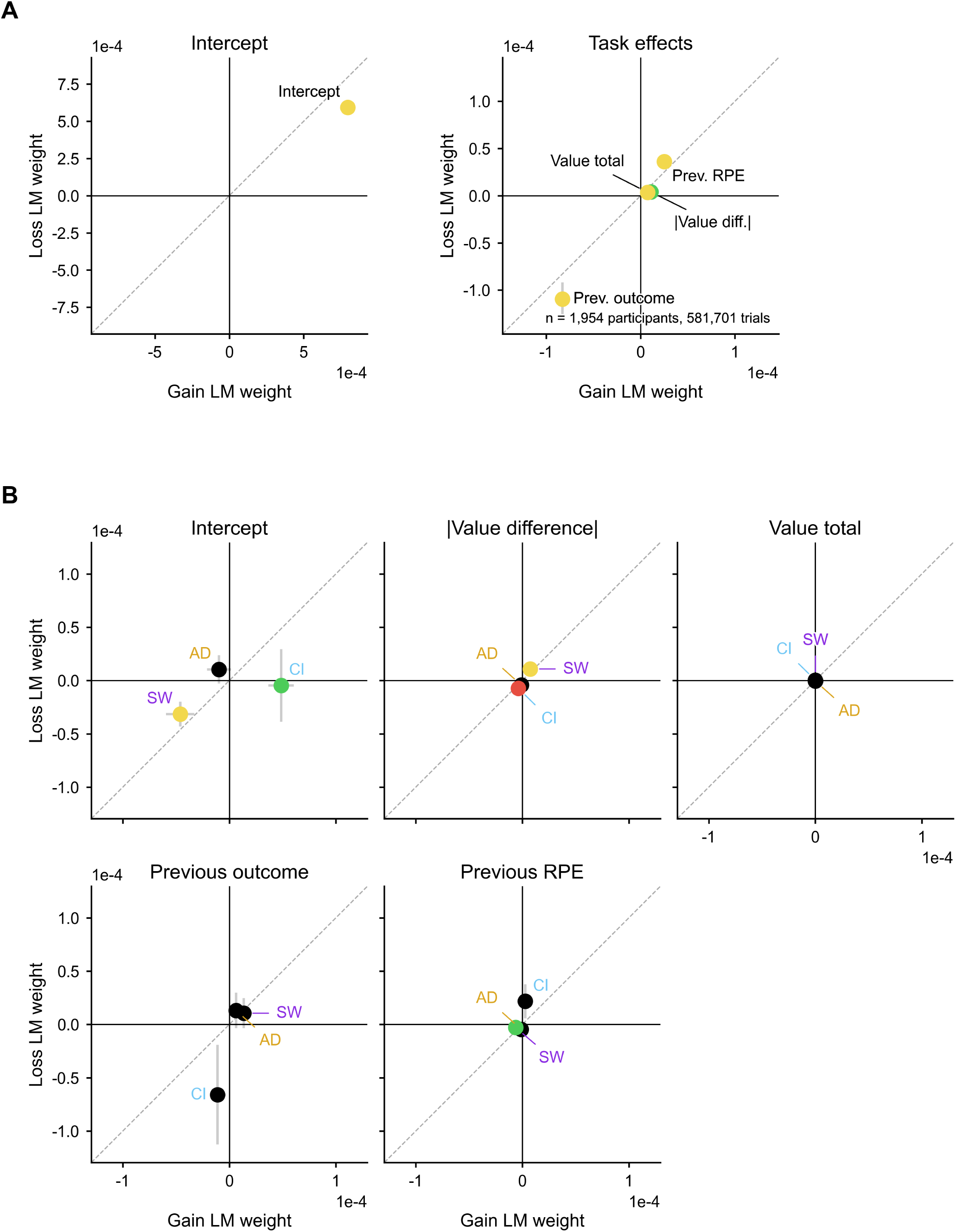
Choice reaction speed is driven by economic choice variables and recent reward prediction errors. A population (cluster-robust) linear model of trial-by-trial choice reaction speed—the reciprocal of choice reaction time, so that a positive coefficient means a faster response—on the absolute value difference between the two offers, the total economic value across them, the previous trial’s outcome and the previous trial’s reward prediction error (RPE); fitted across n=1954 participants and decomposed into gain- and loss-context coefficients (Methods). Layout, SE arms, unity line and the four-class point coloring are as in **Supplementary Figure 2**, with BH–FDR over this model’s 40 context-decomposed effects (20 model terms × 2 contexts); axes are drawn with a ×10^-4^ multiplier. **(A)** Population effects, split across two tiles. Left (*Intercept*): baseline choice speed was higher in gain than in loss (+7.96 × 10^-4^ ± 1.1 × 10^-5^ vs. +5.92 × 10^-4^ ± 1.3 × 10^-5^; both *p <* 0.0001), i.e. choices were slower under the loss framing. Right (*Task effects*): total value was associated with faster choices in both contexts (+7.28 × 10^-6^, *p*_BH_ *<* 0.0001; +3.56 × 10^-6^, *p*_BH_ *<* 0.0001); the absolute value difference was associated with faster choices in gain (+1.09 × 10^-5^, *p*_BH_ *<* 0.0001) but did not reach significance in loss (+3.89 × 10^-6^, *p*_BH_ = 0.16), so its point is green; a larger previous-trial RPE was associated with faster choices in both contexts (+2.51 × 10^-5^ and +3.61 × 10^-5^) whereas a larger previous-trial outcome was associated with slower ones (−8.29 × 10^-5^ and −1.10 × 10^-4^; all four *p*_BH_ *<* 0.0001). **(B)** Interactions of the same effects with AD, CI and SW, one tile per effect (*Intercept*, |*Value difference*|, *Value total*, *Previous outcome*, *Previous RPE*), with the three points labeled by factor. CI interacted positively with the intercept in gain only (+4.86 × 10^-5^, *p*_BH_ = 1.9 × 10^-4^; green) and SW negatively in both contexts (−4.61 × 10^-5^, *p*_BH_ = 1.8 × 10^-3^; −3.14 × 10^-5^, *p*_BH_ = 0.017; yellow), indicating that CI was associated with overall slightly faster choices in gain while SW was associated with slower choices across contexts. For the absolute value difference, SW interacted positively in both contexts (+7.44 × 10^-6^, *p*_BH_ = 7.0 × 10^-3^; +1.08 × 10^-5^, *p*_BH_ = 8.7 × 10^-4^) and CI negatively in loss (−7.30 × 10^-6^, *p*_BH_ = 0.015), so SW psychopathology is associated with stronger and CI psychopathology with weaker modulation of choice reaction speed by economic value; the corresponding AD effects run in the same direction as CI’s but do not survive correction, and AD’s point is black (loss −4.26 × 10^-6^, *p*_BH_ = 0.16). SW also interacted negatively with the previous-trial RPE in gain (−6.27 × 10^-6^, *p*_BH_ = 0.010). No factor interacted significantly with total value or with the previous-trial outcome in either context.

**Supplementary Figure 9.**
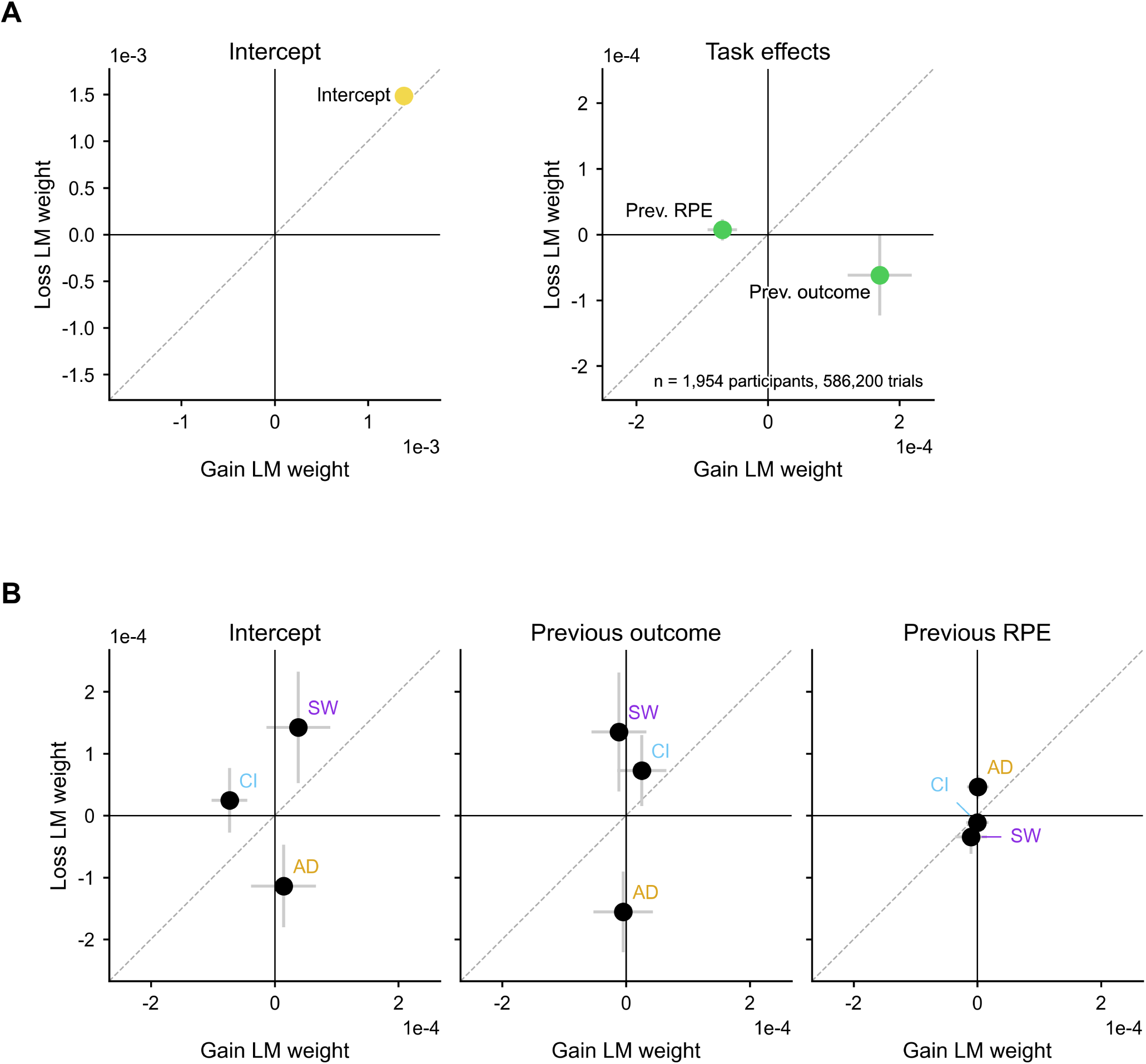
Fixation reaction speed is minimally related to task variables and unrelated to psychopathology. The same modeling approach as **Supplementary Figure 8**, applied to the speed of the trial-initiating fixation response (the reciprocal of fixation reaction time; a positive coefficient means faster) and carrying only the previous trial’s outcome and reward prediction error (RPE) as task regressors; fitted across n=1954 participants, cluster-robust, decomposed by context, with BH–FDR over this model’s 24 context-decomposed effects (12 model terms × 2 contexts). Layout, SE arms, unity line and point coloring are as in **Supplementary Figure 8. (A)** Population effects. Left (*Intercept*, axes ×10^-3^): baseline fixation speed was similar in the two contexts (+1.38 × 10^-3^ ±4.7 × 10^-5^ in gain, +1.49 × 10^-3^ ±5.6 × 10^-5^ in loss; both *p <* 0.0001). Right (*Task effects*, axes ×10^-4^): both history terms reach significance in the gain context only, so both points are green—a larger previous-trial outcome was followed by a faster fixation (+1.70 × 10^-4^, *p*_BH_ = 4.0 × 10^-3^; loss −6.17 × 10^-5^, *p*_BH_ = 0.54) and a larger previous-trial RPE by a slower one (−6.93 × 10^-5^, *p*_BH_ = 0.012; loss +7.10 × 10^-6^, *p*_BH_ = 0.87). **(B)** Interactions of the same effects with AD, CI and SW, one tile per effect (*Intercept*, *Previous outcome*, *Previous RPE*), with the three points labeled by factor. Every point is black: no psychopathology factor interacted with baseline fixation speed or with either history term in either context after correction, the nearest being the CI intercept in gain (*p*_BH_ = 0.056).

**Supplementary Figure 10.**
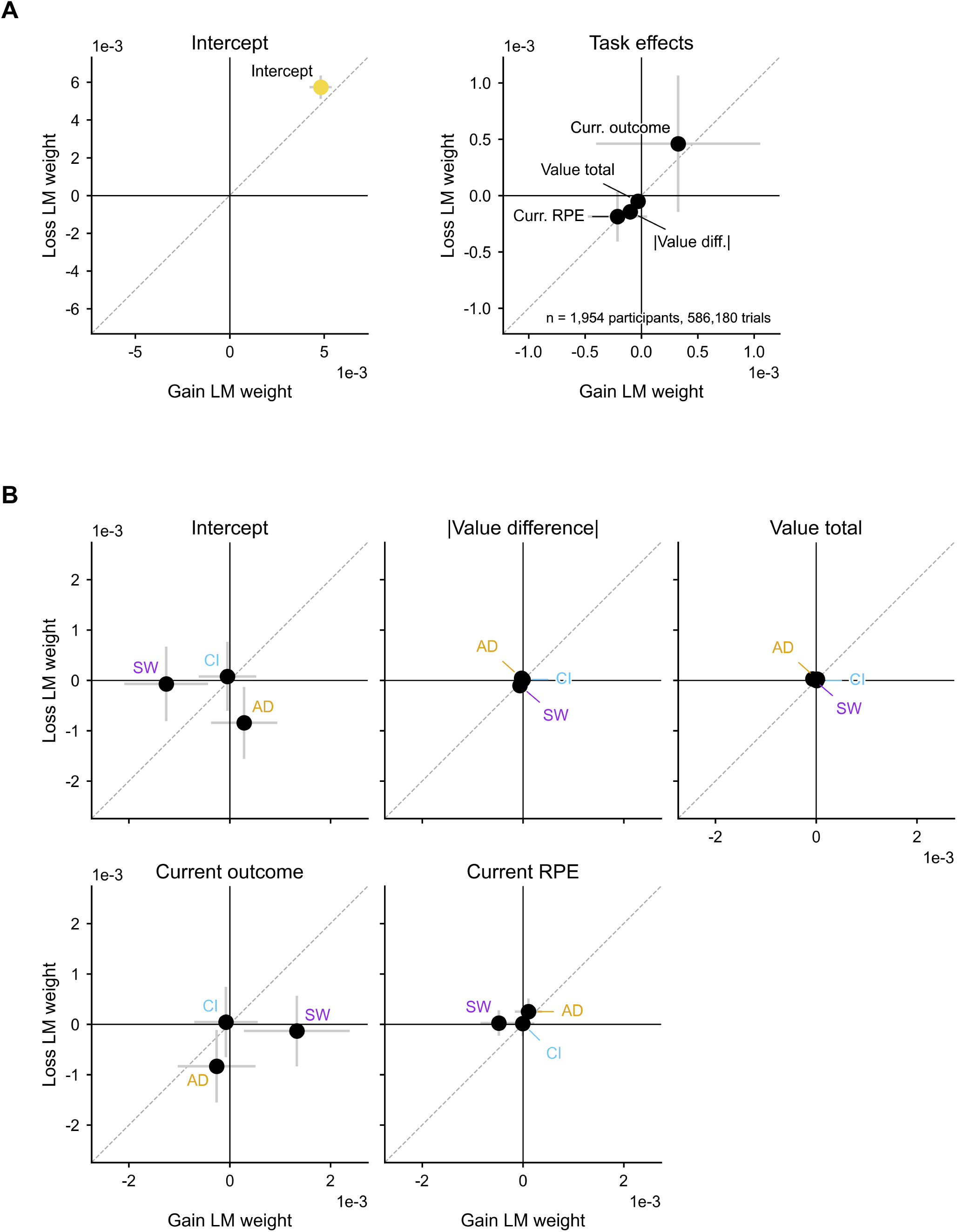
Collection reaction speed is unrelated to task variables and to psychopathology. The same modeling approach as **Supplementary Figure 8**, applied to the speed of the post-outcome collection response (the reciprocal of collection reaction time; a positive coefficient means faster) and carrying the absolute value difference between the two offers, the total economic value across them, and the current trial’s outcome and reward prediction error (RPE) as task regressors; fitted across n=1954 participants, cluster-robust, decomposed by context, with BH–FDR over this model’s 40 context-decomposed effects (20 model terms × 2 contexts). Layout, SE arms, unity line and point coloring are as in **Supplementary Figure 8**; axes are drawn with a ×10^-3^ multiplier. **(A)** Population effects. Left (*Intercept*): baseline collection speed was well above zero in both contexts and slightly higher in loss (+4.82 × 10^-3^ ±5.9 × 10^-4^ in gain, *p*_BH_ *<* 0.0001; +5.74 × 10^-3^ ± 6.2 × 10^-4^ in loss, *p*_BH_ *<* 0.0001). Right (*Task effects*): none of the four task regressors— the absolute value difference, the total value, the current outcome and the current RPE—reached significance in either context after correction, so all four points are black (smallest *p*_BH_ = 0.42). **(B)** Interactions of the same effects with AD, CI and SW, one tile per effect (*Intercept*, |*Value difference*|, *Value total*, *Current outcome*, *Current RPE*), with the three points labeled by factor. All fifteen interactions are non-significant in both contexts and every point is black (smallest *p*_BH_ = 0.77). Collection speed, measured after the choice has been made and the outcome revealed, therefore provides a null comparison for the value- and psychopathology-related modulations of choice reaction speed in **Supplementary Figure 8**.

**Supplementary Figure 11.**
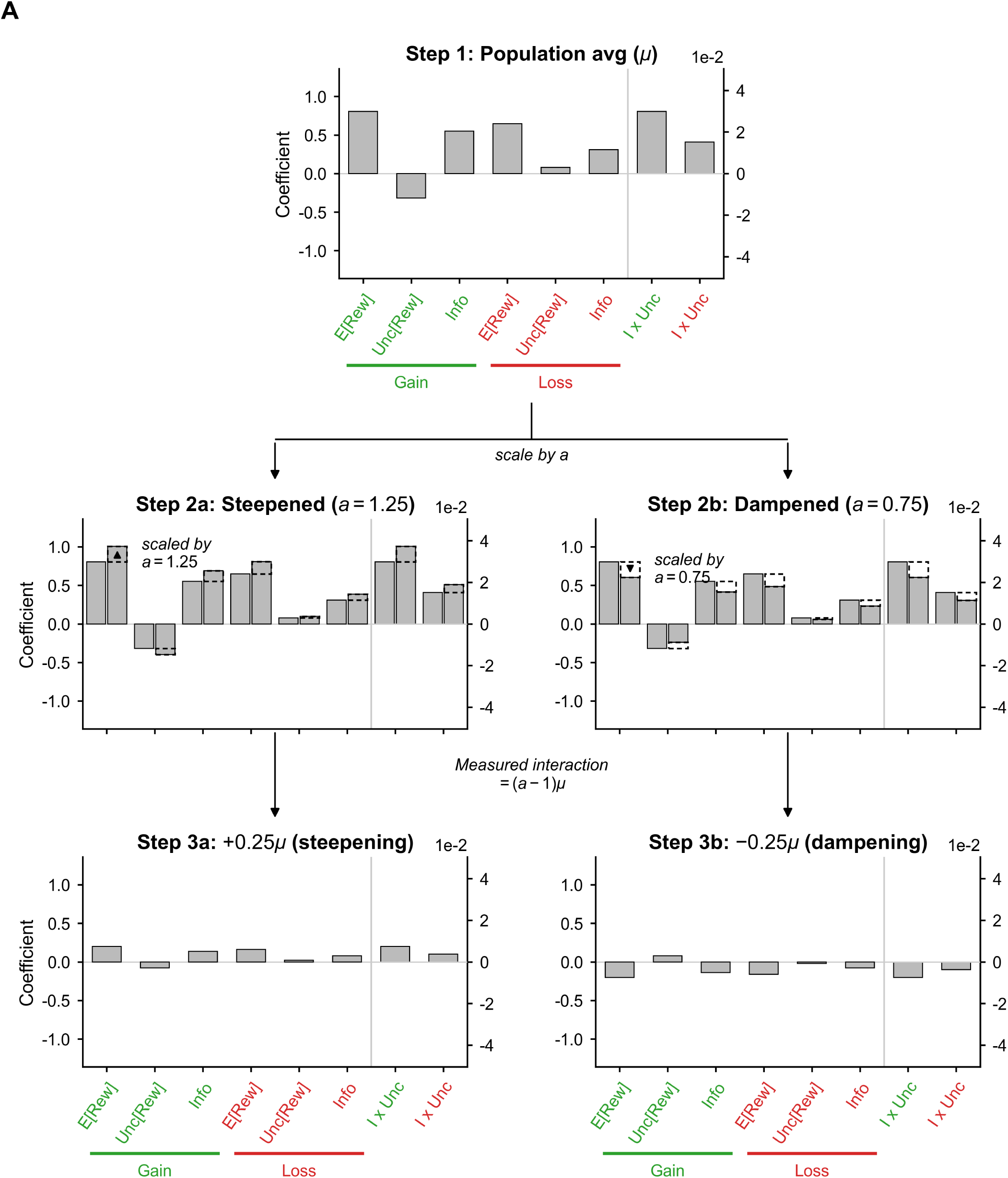
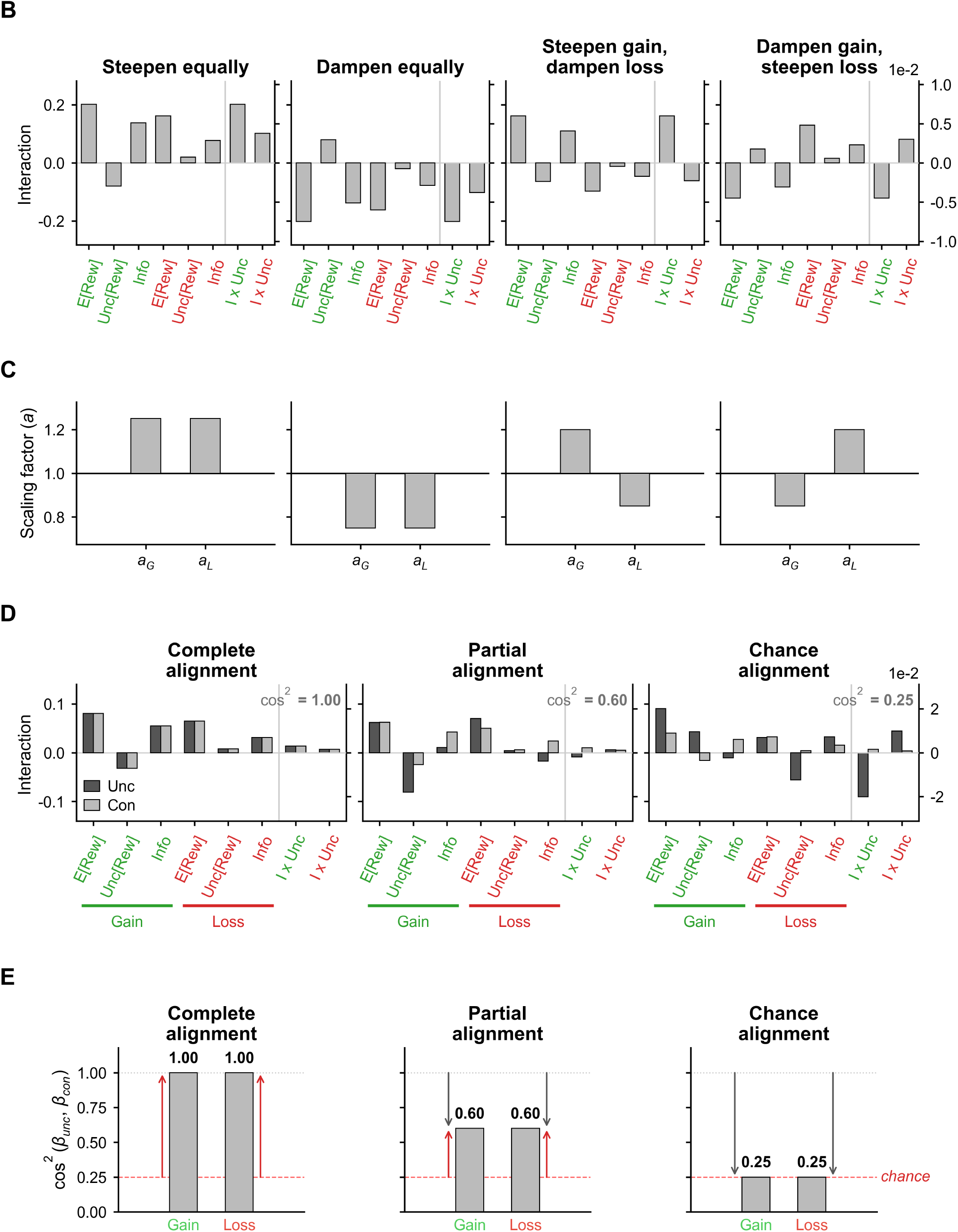
Schematic walkthrough of the scaling account tested in Figure 5. Graphical statement of the “scaling” account — each psychopathology factor multiplies a context’s population-average attribute weights by a single scalar — and of the squared-cosine statistic used to score how well the measured effects match it. The two low-dimensional accounts are presented identically; the context-change counterpart is **Supplementary Figure 13**, and the fitted results are in **Figure 5** and **Supplementary Figure 12**. Every panel here is schematic: apart from the fitted population-average baseline pattern, the bars illustrate hypothetical patterns, not fitted data. The figure runs over two pages under one caption, **(A)** on the first and **(B)**–**(E)** on the second, with the figure tag set at the top right of each. Rows **(A)**, **(B)** and **(D)** use the same eight-slot x-axis — E[Rew], Unc[Rew] and Info in gain (tick labels green), then the same three in loss (red), then, past a vertical divider and drawn against a right-hand axis an order of magnitude finer, the two Info × Unc[Rew] slots (drawn *I x Unc*) — and rows **(A)** and **(D)** add a colored underline and an italic *Gain* or *Loss* label beneath each context block, while row **(B)** carries the context in its colored tick labels alone. **(A)** The hypothesis in three steps, read left to right. *Step 1: Population avg (µ)* is the population-average preference vector *∼µ* over the four attributes in both contexts (y-axis *Coefficient*). An arrow labeled *scale by a* leads to *Step 2a: Steepened (a* = 1.25) and *Step 2b: Dampened (a* = 0.75), which draw the rescaled vector (dashed outline, with a rule marking the original level) beside the original bars and annotate the shift with a triangle and an in-panel note. A second arrow, labeled *measured interaction* = (*a* - 1)*µ*, leads to *Step 3a:* +0.25*µ (steepening)* and *Step 3b:* −0.25*µ (dampening)*: what a factor × attribute interaction would look like under each case — same-signed as *∼µ* for steepening, opposite-signed for dampening, and proportional to *∼µ* slot by slot in both. **(B)** Four example landmarks in the continuous hypothesis space obtained by freeing the two per-context scalars independently: predicted interaction patterns (y-axis *Interaction*) for *Steepen equally*, *Dampen equally*, *Steepen gain, dampen loss* and *Dampen gain, steepen loss*. **(C)** The scaling-factor pairs that generate each landmark in **(B)**: two bars per tile on the y-axis *Scaling factor (a)*, drawn from a baseline of 1, the gain scalar first and the loss scalar second (the ticks print the two scalars in their abbreviated symbol form). **(D)** What imperfect alignment looks like on the bars: three examples — *Complete alignment*, *Partial alignment* and *Chance alignment* — each drawing a hypothetical unconstrained interaction profile (dark, legend *Unc*) against the constrained scaling prediction (light, *Con*) on the y-axis *Interaction*, with the resulting squared cosine printed in the cell (cos^2^ = 1.00, 0.60 and 0.25). **(E)** The same three scenarios reduced to the alignment statistic, y-axis cos^2^(*β*_unc_*, β*_con_), one bar for gain and one for loss with its value printed above. The gray dotted line at 1.00 marks perfect alignment and the red dashed line, labeled *chance*, marks the expected value of cos^2^ for a random direction relative to the two-scalar hypothesis space (2*/*8 = 0.25) — an expected value under chance, not a significance threshold. The arrow encoding used on the fitted figures is illustrated here: a gray downward arrow measures the gap below perfect alignment and a red upward arrow the height above chance, so the complete case carries only red arrows, the chance case only gray ones, and the partial case both. The arrows are drawn without significance marks in this schematic.

**Supplementary Figure 12.**
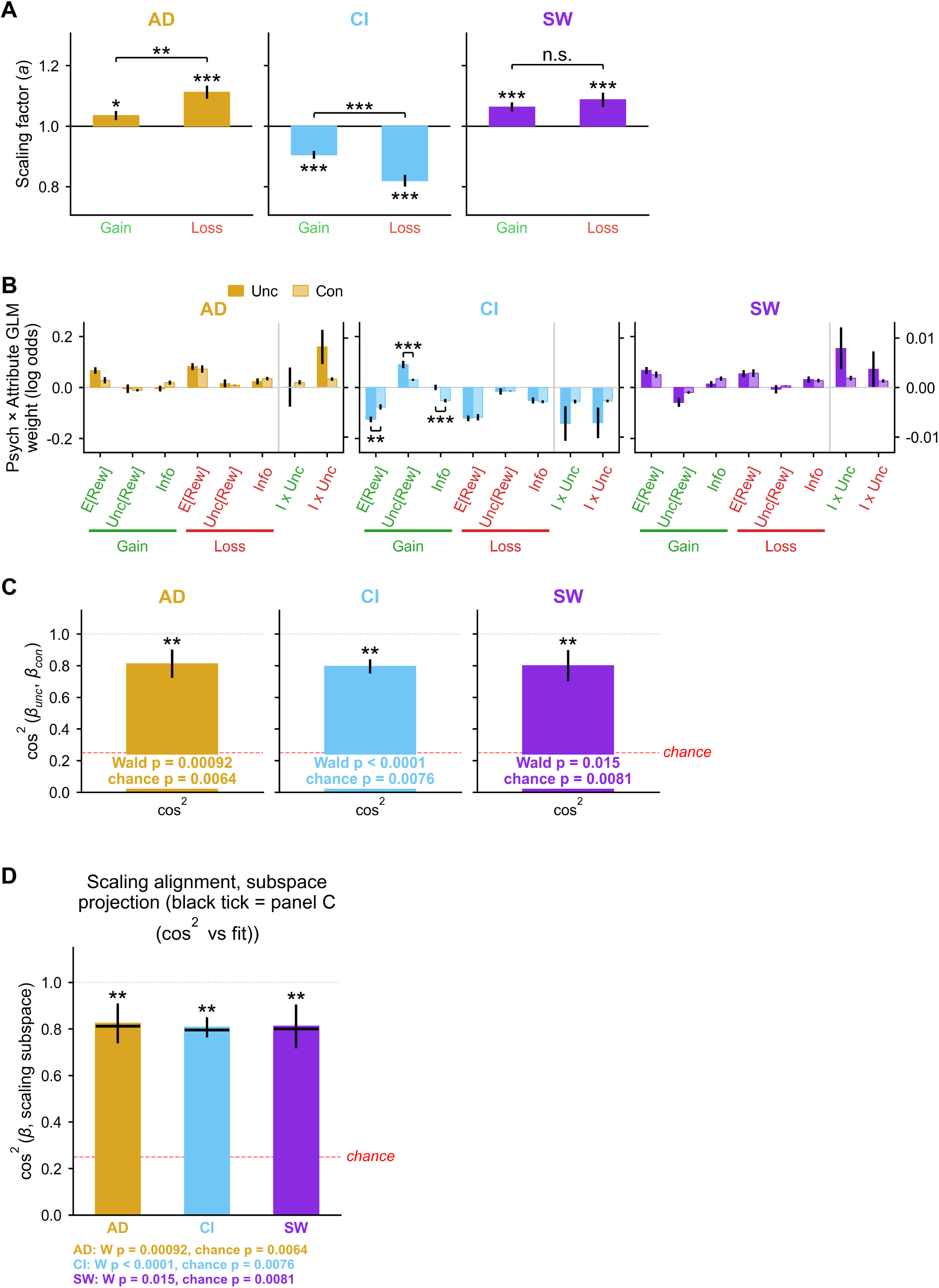
Maximum-likelihood version of the scaling test in Figure 5. **Figure 5** pins each context’s scaling factor to that factor’s own fitted E[Rew] interaction. This figure reports the original maximum-likelihood version of the same test, in which the two per-context scalars were instead estimated by re-fitting the logistic GLM under the scaling constraint (Methods): the fitted scalars **(A)**, the attribute-level comparison against the unconstrained model **(B)**, the joint eight-dimensional alignment statistic **(C)**, and a refit-free robustness check **(D)**. Rows **(A)**–**(C)** are read across three cells, one per psychopathology factor, titled and colored AD (goldenrod), CI (baby blue) and SW (blue violet). Error bars show cluster-robust SE unless stated otherwise; *, **, *** indicate *p <* 0.05, 0.01, 0.001, and *n.s.* is printed where a drawn comparison does not clear 0.05. **(A)** Fitted scaling factor in each context: one bar per context drawn from a baseline of 1 on the y-axis *Scaling factor (a)*, with x-axis ticks *Gain* (green) and *Loss* (red). Per-bar stars are two-sided *z* tests of *a* = 1, and the bracket above each cell tests the gain-minus-loss difference by an independent-SE Wald *z*. AD and SW steepened the population-average attribute weights in both contexts (AD *a*_gain_ = 1.035, *p* = 1.8 × 10^-2^, *a*_loss_ = 1.112, *p <* 0.0001; SW *a*_gain_ = 1.063, *p <* 0.0001, *a*_loss_ = 1.087, *p* = 1.9 × 10^-4^), whereas CI dampened them in both (*a*_gain_ = 0.905, *p <* 0.0001; *a*_loss_ = 0.819, *p <* 0.0001). Scaling departs further from 1 in loss for AD (difference −0.077, *z* = −2.93, *p* = 3.4 × 10^-3^) and for CI (+0.086, *z* = +3.67, *p* = 2.4 × 10^-4^) but not for SW (−0.024, *z* = −0.87, *p* = 0.38). Direction, ordering and the AD/CI gain–loss asymmetry all reproduce the pinned scalars of **Figure 5B**. **(B)** Unconstrained vs. constrained interaction profiles: for each factor, the eight attribute-context interaction coefficients drawn as pairs of bars — unconstrained (dark, legend *Unc*) against the maximum-likelihood scaling prediction (light, *Con*) — on the y-axis *Psych* × *Attribute GLM weight (log odds)* and the eight-slot x-axis described for **Supplementary Figure 11**. Unlike **Figure 5A** no slot is pinned here, so all eight comparisons carry a test. Brackets and stars mark attribute-level departures from the scaling prediction (two-sample *z* with independent SEs, BH-corrected within factor over that factor’s 8 attribute-context effects; a departure whose *p*_BH_ fell in [0.05, 0.1) would be drawn with its literal p-value instead of stars). Only three departures are significant, all for CI and all in the gain context: E[Rew] (*p*_BH_ = 4.0 × 10^-3^), Unc[Rew] (*p*_BH_ = 1.6 × 10^-4^) and Info (*p*_BH_ = 1.6 × 10^-4^). AD and SW carry no bracket at all (nearest *p*_BH_ = 0.22 and 0.24). **(C)** Joint alignment: the squared cosine between each factor’s unconstrained 8-dimensional coefficient vector and its fitted scaling prediction, on the y-axis cos^2^ (*β*_unc_ *, β*_con_), one bar per factor over a single cos^2^ tick. The gray dotted line at 1.00 marks perfect alignment and the red dashed line, labeled *chance*, the expected value of cos^2^ under the two-scalar chance model (2*/*8 = 0.25) — an expected value, not a threshold; error bars are the SD of a parametric bootstrap that perturbs the coefficients by their SEs with the constrained fit held fixed across draws. All three factors align at about 0.80 (AD = 0.81; CI = 0.80; SW = 0.80). The two-line annotation inside each cell reports both tests: the cluster-robust Wald test of exact scaling, which rejects for every factor (AD *W* (6) = 22.6, *p* = 9.2 × 10^-4^; CI *W*(6) = 72.2, *p <* 0.0001; SW *W*(6) = 15.9, *p* = 1.5 × 10^-2^), and the calibrated above-chance test — a null in which a factor effect of matched magnitude points in a random direction (Methods) — which is significant for every factor (AD *p* = 0.0064; CI *p* = 0.0076; SW *p* = 0.0081). The stars above the bars are that above-chance test. Each factor’s effect pattern is therefore far closer to uniform scaling than to a random direction while still being distinguishable from exact scaling. **(D)** Refit-free robustness check for **(C)**, titled *Scaling alignment, subspace projection*. Alignment is scored instead as the orthogonal projection of each factor’s coefficient vector onto the two-scalar scaling hypothesis space, 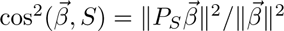, on the y-axis cos^2^ (*β*, scaling subspace) with x-axis ticks AD, CI and SW in the factor colors; the short black tick across each bar is that factor’s constrained-fit value from **(C)**, and the footer beneath the axis repeats each factor’s Wald and above-chance p-value. Because this statistic requires no model refit, its bootstrap recomputes the projection on every draw, and both its ceiling (exactly 1) and its chance expected value (exactly 2*/*8) are analytic. It reproduces **(C)** closely (AD = 0.82; CI = 0.81; SW = 0.81; every factor within 0.02 of its constrained-fit value).

**Supplementary Figure 13.**
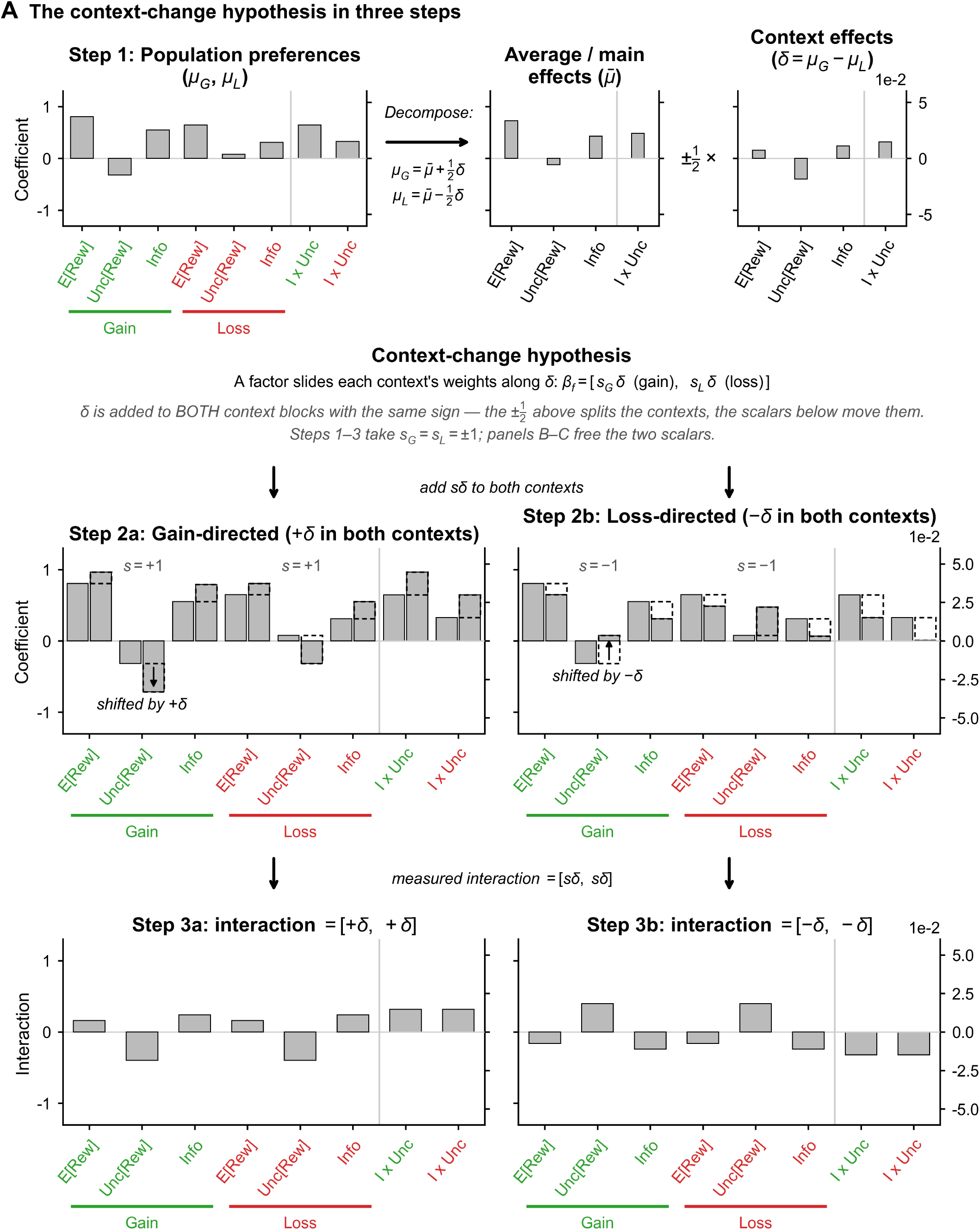
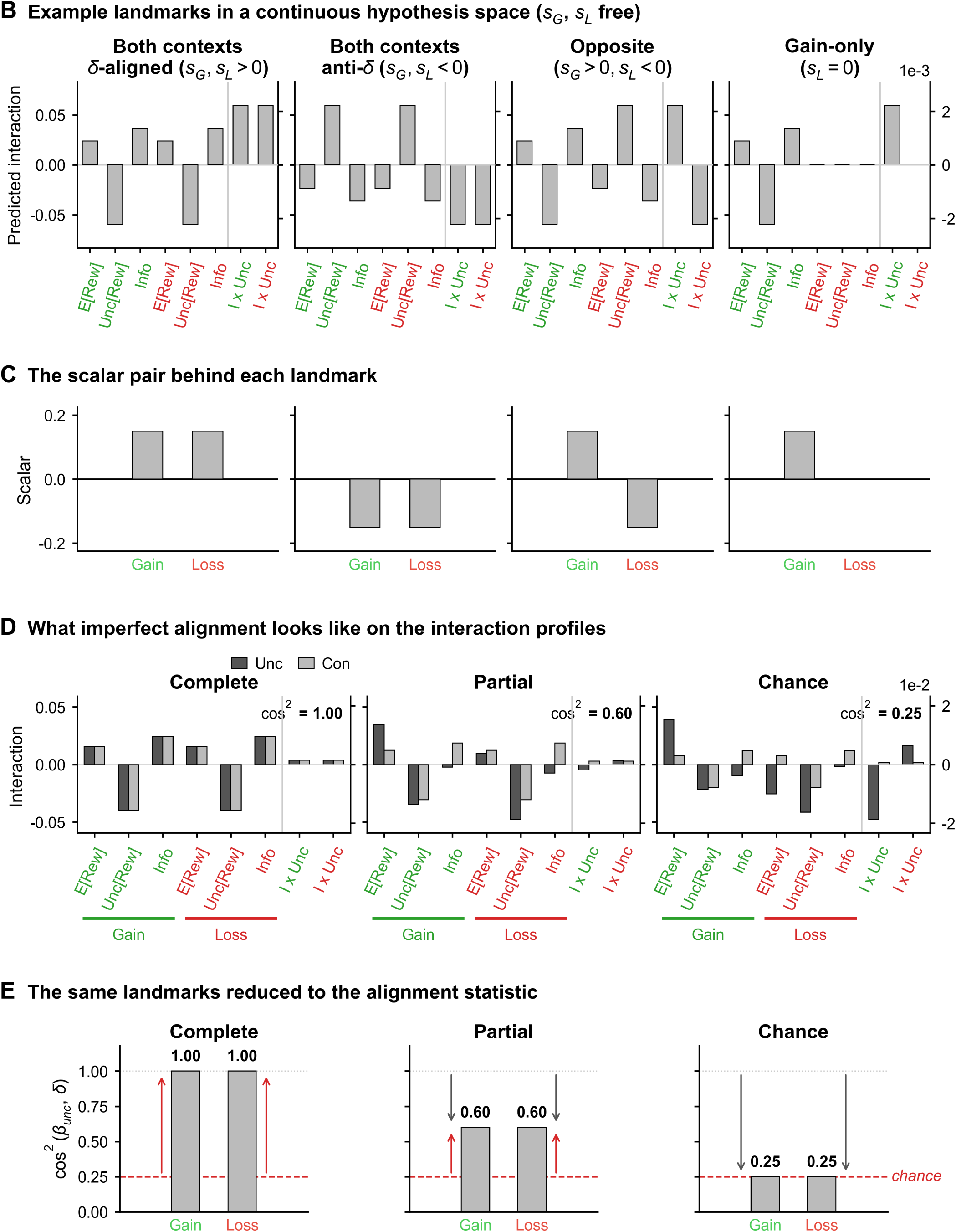
Schematic walkthrough of the context-change account. Graphical statement of the “context-change” account tested in Supplementary Figure 14, in which a psychopathology factor slides each context’s preference weights along 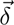, the direction along which the population’s own gain and loss preferences differ. It is the counterpart of **Supplementary Figure 11**, which walks through the scaling account tested in **Figure 5**. Every panel here is schematic: apart from the fitted population-average baseline pattern, the bars illustrate hypothetical patterns rather than fitted data. Rows **(A)**, **(B)** and **(D)** use the same eight-slot x-axis as **Supplementary Figure 11** (E[Rew], Unc[Rew], Info in gain, then in loss, then the two Info × Unc[Rew] slots past a divider on a finer right-hand axis, with green and red tick labels); as there, rows **(A)** and **(D)** add the *Gain* / *Loss* block underlines, while row **(B)** carries the context in its colored tick labels alone. **(A)** The account in one row. *Step 1: Population preferences (µ*_G_*, µ*_L_) is split, via an arrow labeled *Decompose:* and the identity 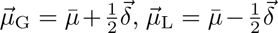, into *Average / main effects (µ̄)* and, past a 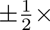 step, *Context effects (δ* = *µ*_G_ - *µ*_L_); these two cells are drawn on a four-slot attribute axis with black tick labels, because 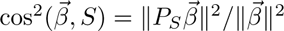 are single vectors over the four attributes rather than context-split ones. A statement beneath the decomposition gives the hypothesis — a factor slides each context’s weights along 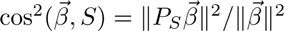, with its own scalar multiplying 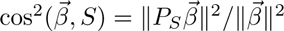 in each context — and warns in italics that 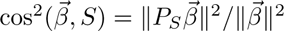 is added to *both* context blocks with the *same* sign: the 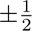 above splits the contexts, whereas the scalars below move them; steps 1–3 fix both scalars at ±1 and rows **(B)**–**(C)** free them. The walkthrough then proceeds from the same cell via an arrow labeled *add sδ to both contexts*, *Step 2a: Gain-directed (*+*δ in both contexts)* and *Step 2b: Loss-directed (*-*δ in both contexts)*, each drawing the shifted vector (dashed outline, with a rule marking the original level) beside the original bars and tagged with its scalar value; a final arrow labeled *measured interaction* = [*sδ, sδ*] leads to *Step 3a* and *Step 3b*, the interaction patterns [+*δ,* +*δ*] and [−*δ,* -*δ*] that would then be measured. **(B)** Four example landmarks in the continuous hypothesis space obtained by freeing the two scalars: predicted interaction patterns (y-axis *Predicted interaction*) for *δ*-aligned effects in both contexts, anti-*δ* effects in both contexts, opposite-signed effects, and a gain-only effect whose loss scalar is zero; each tile’s title states the sign condition it illustrates. **(C)** The scalar pairs generating each landmark in **(B)**: two bars per tile on the y-axis *Scalar*, drawn from a baseline of 0 — unlike the scaling factors of **Supplementary Figure 11C**, whose no-effect value is 1 — with the gain scalar first and the loss scalar second (ticks *Gain* and *Loss*). **(D)** Three schematic unconstrained (dark, *Unc*) vs. constrained (light, *Con*) profiles on the y-axis *Interaction*, under *Complete alignment* (cos^2^ = 1.00), *Partial alignment* (0.60) and *Chance alignment* (0.25) against the fitted context-change prediction. **(E)** The corresponding squared-cosine summaries, y-axis cos^2^ (*β*_unc_*, δ*), one bar for gain and one for loss with the value printed above; the gray dotted line at 1.00 marks perfect alignment and the red dashed line, labeled *chance*, the expected value of cos^2^ under chance (2*/*8 = 0.25), not a significance threshold. The gray downward and red upward teaching arrows use the same encoding as **Supplementary Figure 11E** and carry no significance marks here.

**Supplementary Figure 14.**
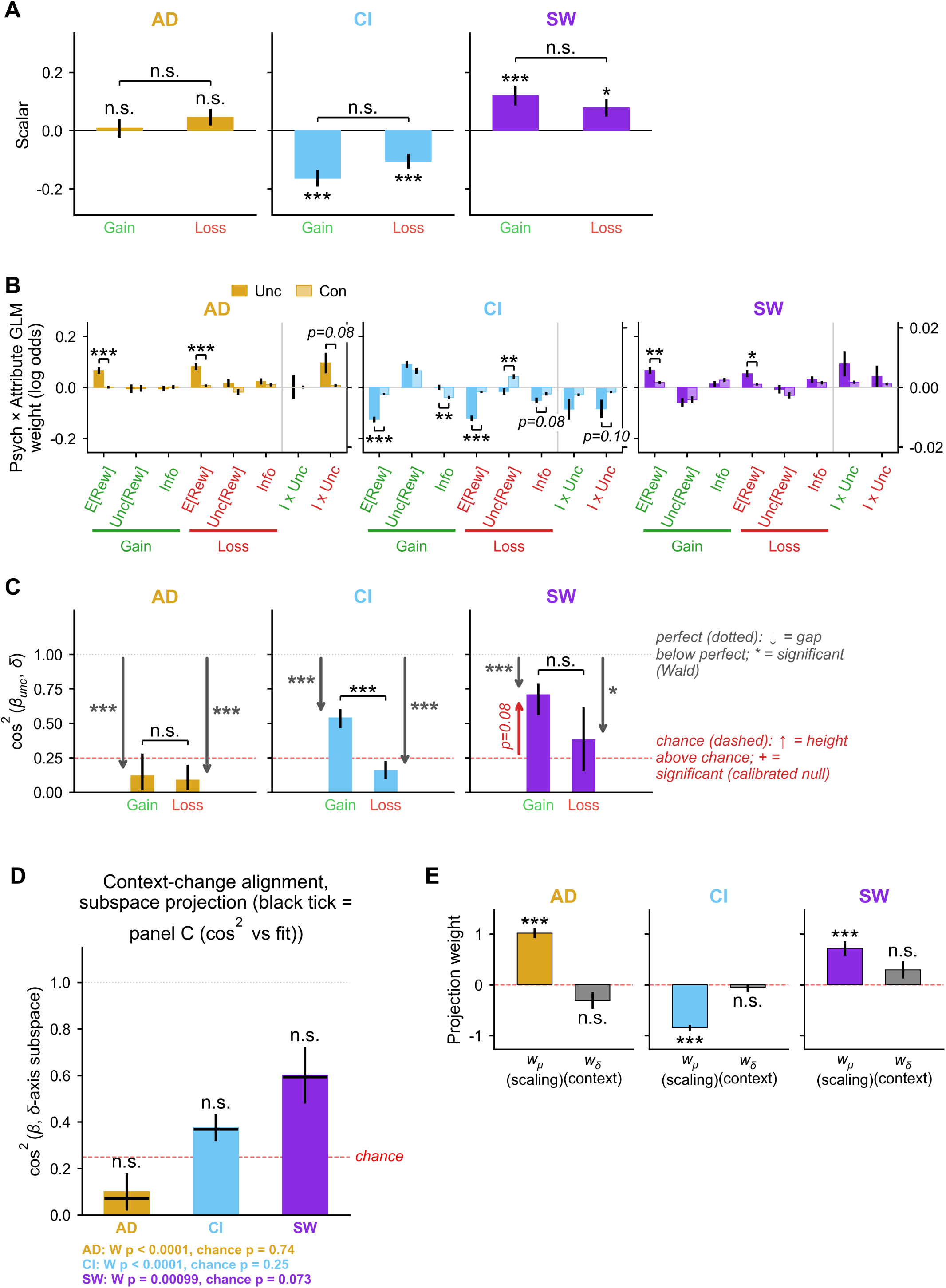
Testing a context-change account of psychopathology-related changes in economic preferences. Complementary to the scaling test of **Figure 5**, we tested the “context-change” account laid out schematically in **Supplementary Figure 13**. The unconstrained comparison model is the one **Figure 5** uses (8 free factor × attribute interaction parameters per factor); the constrained model replaces them with scalar interactions along the contextmodulation direction 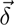, carrying separate gain and loss coefficients (*s*_gain_, *s*_loss_) so that gain-directed or loss-directed effects are permitted in either context (Methods). Both models here are fit by maximum likelihood, because the pinning device of **Figure 5** has no stable counterpart along 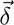: 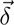 is dominated by Unc[Rew] rather than E[Rew] (the E[Rew] component carries 10.5% of ||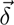||^2^, vs. 61.6% of ||*µ̂*_gain_ ||^2^), so pinning to E[Rew] would inflate the scalars 4–51-fold relative to the maximum-likelihood fit; the alignment statistics of **(C)**–**(D)** are scale-invariant and identical under either choice. The matched scaling comparison for these panels is therefore the maximum-likelihood version in **Supplementary Figure 12**, not **Figure 5** itself. Except for **(D)**, each row is read across three cells, one per factor, titled and colored AD (goldenrod), CI (baby blue) and SW (blue violet). Panels **(A)**–**(C)** report the context-change test, **(D)** is its robustness check and **(E)** the competitive decomposition. Error bars show cluster-robust SE unless stated otherwise; *, **, *** indicate *p <* 0.05, 0.01, 0.001, and *n.s.* is printed where a drawn comparison does not clear 0.05. **(A)** Fitted context-change scalars: one bar per context drawn from a baseline of 0 on the y-axis *Scalar*, with x-axis ticks *Gain* (green) and *Loss* (red), per-bar two-sided *z* tests against zero, and a bracket for the gain-minus-loss difference. CI loaded significantly negatively on 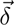 in both contexts (*s*_gain_ = −0.164, *p <* 0.0001; *s*_loss_ = −0.105, *p <* 0.0001) and SW significantly positively (*s*_gain_ = +0.120, *p* = 4.4 × 10^-4^; *s*_loss_ = +0.078, *p* = 1.0 × 10^-2^), whereas neither AD scalar reached significance (+0.008, *p* = 0.80; +0.046, *p* = 0.11). No gain–loss scalar difference was significant (all three brackets *n.s.*; smallest *p* = 0.13). **(B)** Unconstrained (dark, *Unc*) vs. constrained (light, *Con*) interaction profiles on the y-axis *Psych* × *Attribute GLM weight (log odds)* and the eight-slot x-axis of **Supplementary Figure 11**. Brackets and stars mark per-attribute departures from the context-change prediction (two-sample *z* with independent SEs, BH-corrected within factor over that factor’s 8 attribute-context effects), with the literal p-value drawn in place of stars where *p*_BH_ falls in [0.05, 0.1]. In contrast to the scaling test, significant residuals remain for all three factors, and they are concentrated on E[Rew] in both contexts (AD gain and loss both *p*_BH_ *<* 0.0001; CI gain and loss both *p*_BH_ *<* 0.0001; SW gain 7.5 × 10^-3^, loss 2.1 × 10^-2^). CI carries two further significant departures — Info in gain (*p*_BH_ = 3.3 × 10^-3^) and Unc[Rew] in loss (*p*_BH_ = 1.1× 10^-3^) — and two marginal ones drawn with their literal values (Info in loss, *p* = 0.08; Info × Unc[Rew] in loss, *p* = 0.10); AD’s Info × Unc[Rew] slot in loss is likewise marginal (*p* = 0.08). **(C)** Per-context alignment: the squared cosine between each context’s unconstrained coefficient vector and the fitted context-change prediction for that context, on the y-axis cos^2^ (*β*_unc_ *, δ*) with x-axis ticks *Gain* (green) and *Loss* (red), a gray dotted line at 1.00 for perfect alignment and a red dashed line at 0.25 for chance. The italic key in the right margin states the encoding, which is the one used in **Figure 5C**: a gray downward arrow measures the gap below perfect alignment and carries asterisks from a cluster-robust Wald test (*χ*^2^ (3)), while a red upward arrow measures the height above chance and carries crosses from the calibrated effect-direction null, a marginal chance p-value being drawn as its literal value instead. Confidence intervals are drawn as the interval itself rather than anchored on the bar. Alignment departed significantly from perfect for every context and factor (AD gain = 0.120, *χ*^2^(3) = 31.3; AD loss = 0.088, *χ*^2^ = 36.9; CI gain = 0.539, *χ*^2^ = 112.3; CI loss = 0.154, *χ*^2^ = 98.9; SW gain = 0.706, *χ*^2^ = 20.7; SW loss = 0.380, *χ*^2^ = 10.0; all six *p* ≤ 0.019) and did not exceed chance in any cell, the closest being SW in gain (*p* = 0.080, drawn as its literal value). The gain-versus-loss brackets show no asymmetry for AD (*p* = 0.82) or SW (*p* = 0.19); CI’s alignment, while not itself above chance in either context, was significantly higher in gain (cos^2^ = 0.54) than in loss (cos^2^ = 0.15; bootstrap *p <* 0.0001). **(D)** Refit-free robustness check for **(C)**, titled *Context-change alignment, subspace projection*: the same orthogonal subspace projection as **Supplementary Figure 12D**, computed for the two-scalar context-change hypothesis space, on the y-axis cos^2^ (*β*, *δ*-axis subspace), with the short black tick on each bar giving that factor’s joint constrained-fit value and the footer beneath the axis repeating each factor’s Wald and above-chance p-value. It reproduces the joint statistic closely (projection AD = 0.10, CI = 0.38, SW = 0.60 against constrained-fit AD = 0.07, CI = 0.37, SW = 0.59; every factor within 0.03), and all three bars are marked *n.s.* because none exceeds the calibrated chance null (AD *p* = 0.74; CI *p* = 0.25; SW *p* = 0.073). Note that the dashed line is the null mean: the null distribution of cos^2^ is strongly right-skewed, so a bar sitting above the line — SW’s, here — is not by itself significant. **(E)** Competitive decomposition. Each factor’s unit-normalized 8-dimensional coefficient vector is jointly projected onto the plane spanned by the scaling axis *µ̂*_8D_ and the context-change axis *δ̂*_8D_, which are themselves correlated (cos^2^ (*µ̂*, *δ̂*) = 0.38). Bars give the two projection weights on the y-axis *Projection weight*, with x-axis ticks *w*_µ_ *(scaling)*, drawn in the factor color, and *w*_δ_ *(context)*, drawn gray, 95% intervals from a 10,000-draw parametric bootstrap, and a red dashed line at zero. The scaling weight is highly significant for all three factors and carries the sign of that factor’s effect (AD *w*_µ_ = +1.02, *p <* 0.0001; CI *w*_µ_ = −0.85, *p <* 0.0001; SW *w*_µ_ = +0.72, *p <* 0.0001), whereas the context-change weight is significant for none (AD *w*_δ_ = −0.31, *p* = 0.062; CI *w*_δ_ = −0.06, *p* = 0.47; SW *w*_δ_ = +0.30, *p* = 0.081); these are two-sided normal p-values from the bootstrap SE, uncorrected. Adding the context-change axis raises the captured variance by at most 0.06 (cos^2^ of the plane vs. cos^2^ along the single fixed axis *µ̂*_8D_ alone: AD 0.75 vs. 0.69; CI 0.78 vs. 0.78; SW 0.86 vs. 0.81). This panel’s single-axis cos^2^ values are 1-D quantities by design — the decomposition competes two specific fixed axes — and are therefore not comparable to the two-scalar alignment statistics of **Supplementary Figure 12C** and of panel **(C)**. Takeaway: context-change alignment is not above chance for any factor and adds no reliable explanatory power beyond scaling — constraining the interpretation of psychopathology effects as reference-point or negativity-bias shifts — while the residual structure that neither low-dimensional account explains (Figure 5) remains open.

**Supplementary Figure 15.**
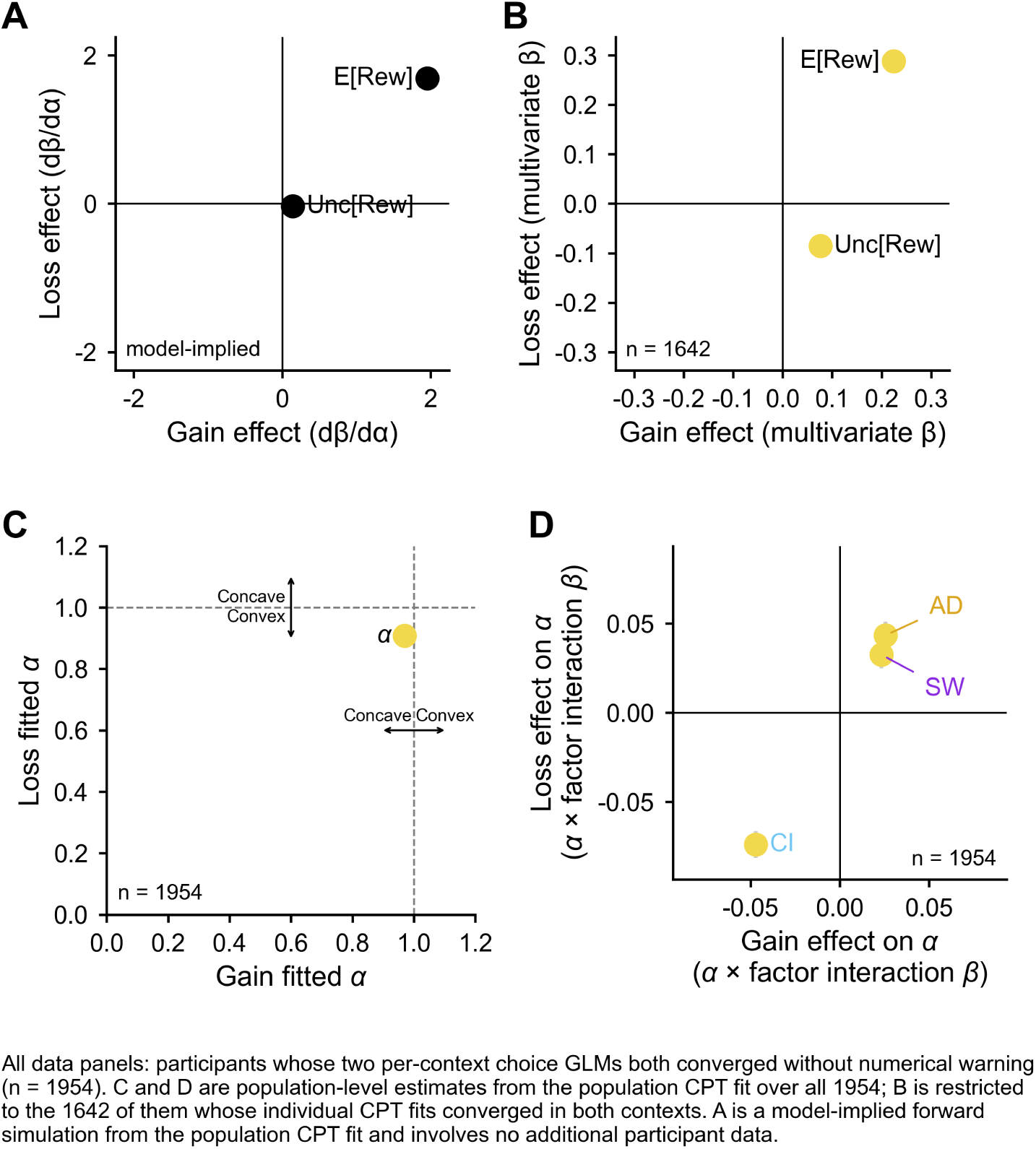
Prospect-theory account of economic preferences for reward expectation and risk. The choice data were additionally fit with a cumulative-prospect-theory (CPT) value function whose only free curvature parameter is the magnitude-distortion exponent *α* (Methods). This figure gives the fitted model’s own implied mapping from *α* to the choice-GLM weights **(A)**, relates *α* to the model-free choice-GLM weights at the individual level **(B)**, gives its population value in each context **(C)**, and gives its association with psychopathology **(D)**. All four panels plot a gain-context quantity on the x-axis against the matched loss-context quantity on the y-axis; in **(B)**–**(D)**, point color, marker labeling and error-bar conventions are as in **Figure 2D**: color marks a two-tailed test against the panel’s reference value that is significant in gain (green), loss (red), both (yellow) or neither (black). Every p-value in this figure is twotailed and uncorrected (Methods). **(A)** Model-implied prediction from the fitted CPT model itself. Using the population CPT fit (Prelec probability weighting fixed at the population estimate *a*_w_ = 0.646, outcomes in task coin units, choice probability logistic in the prospect-value difference), choices were simulated deterministically for the task’s offer pairs across a grid of *α* values, and the manuscript’s logistic choice model (choice ∼ ΔE[Rew] + ΔUnc[Rew], no intercept) was refit to the model-implied choice probabilities in each context (Methods). Plotted are the local derivatives *dβ/dα* of the two GLM weights at the published context-specific *α* (gain 0.970, loss 0.908), gain on x vs. loss on y. Increasing *α* strongly increases the E[Rew] weight in both contexts (*dβ/dα* = +1.95 in gain, +1.69 in loss) while leaving the Unc[Rew] weight nearly unchanged (+0.14, −0.03), so *α* functions largely as an E[Rew]-sensitivity dial. Points are exact model-implied quantities, drawn in black; error bars and significance tests do not apply. **(B)** Individual *α* against individual preference weights: one labeled point per predictor giving the coefficient of a linear model that regresses each participant’s own fitted *α* on their own E[Rew] and Unc[Rew] choice-GLM weights, fit separately within each context, plotted as the gain coefficient (x-axis *Gain effect (multivariate β)*) against the loss coefficient (y-axis *Loss effect (multivariate β)*) on axes made symmetric about zero, with solid lines at zero. The model contains these two preference weights only, and only the *n* = 1642 participants whose CPT model converged in both contexts contribute; SE bars are smaller than the markers. Both points are yellow. A stronger E[Rew] weight went with a higher *α* in both contexts (+0.225 ± 0.004 in gain, *t*(1639) = +51.9; +0.288 ± 0.005 in loss, *t* = +58.8; both *p <* 0.0001), consistent with the monetary amounts used here sitting in a range where the power distortion steepens the utility difference between offers. A stronger Unc[Rew] weight went with a higher *α* in gain (+0.076 ± 0.002, *t* = +30.9) but a lower *α* in loss (−0.085 ± 0.003, *t* = −29.5; both *p <* 0.0001), so this relationship reverses across contexts. **(C)** Population *α* in gain vs. loss: the single population-level *α* of the CPT fit, on fixed axes running 0 to 1.2 (x *Gain fitted α*, y *Loss fitted α*). This is the one panel of the family whose dashed reference crosshair sits at *α* = 1 rather than at zero, and double-headed arrows on the crosshair label the curvature implied on each side of it (concave vs. convex over gains along the x-axis; concave vs. convex over losses along the y-axis). Standard errors are cluster-robust (sandwich) and smaller than the marker. *α* was below 1 in both contexts (gain *α* = 0.970 ± 0.005; loss *α* = 0.908 ± 0.007; both *p <* 0.0001), placing the point in the quadrant marked concave over gains and convex over losses. **(D)** Psychopathology effects on *α*: the conditional effect of each psychopathology factor on *α* in gain (x-axis *Gain effect on alpha*) vs. loss (y-axis *Loss effect on alpha*), one point per factor labeled in that factor’s color (AD goldenrod, CI baby blue, SW blue violet), with cluster-robust SE arms drawn over rather than behind the markers. All three points are yellow. AD and SW were associated with a higher *α* in both contexts (AD gain +0.026 ± 0.005, *p <* 0.0001, loss +0.043 ± 0.008, *p <* 0.0001; SW gain +0.023 ± 0.005, *p <* 0.0001, loss +0.033 ± 0.008, *p <* 0.0001) and CI with a lower *α* in both (gain −0.047 ± 0.005; loss −0.074 ± 0.008; both *p <* 0.0001) — the same AD-and-SW versus CI split that the reward-sensitivity interactions of **Figure 3C** show in the model-free GLM.

**Supplementary Figure 16.**
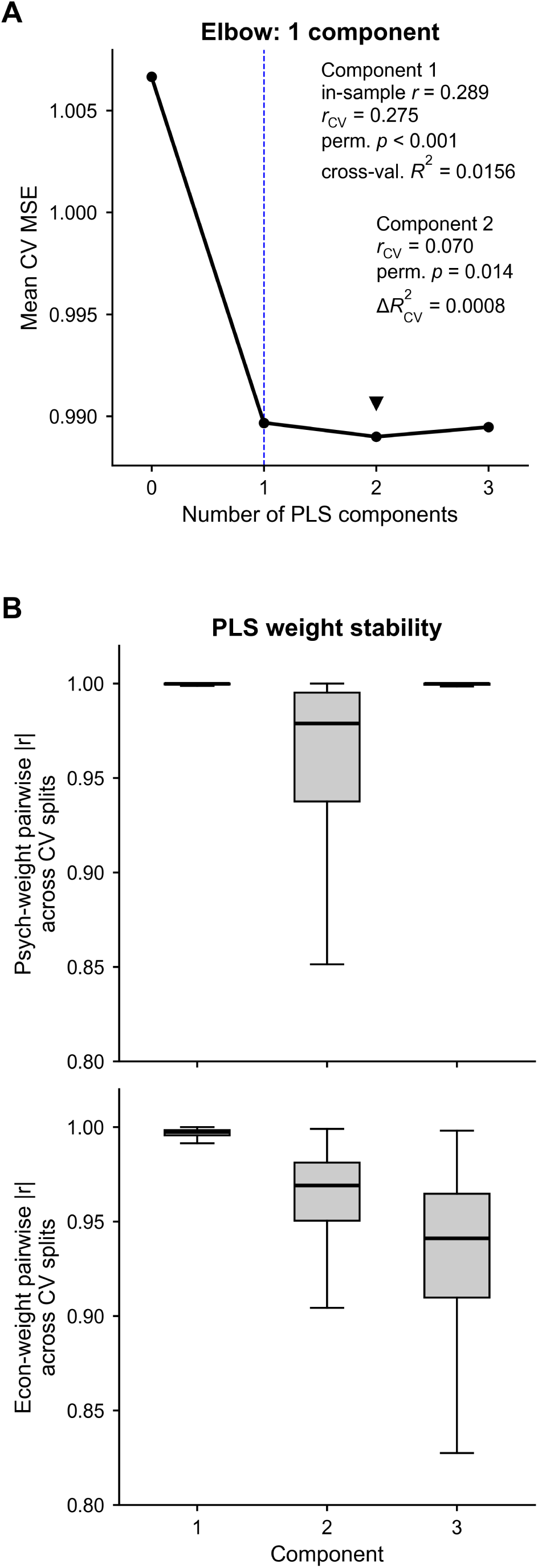
Component selection for the latent dimension of shared economic–psychopathological variation. Component-selection diagnostics for the partial least squares (PLS) regression of **Figure 6**, which predicts the individual economic preference GLM weights (E[Rew], Unc[Rew], Info and Info × Unc[Rew] in gain and loss; 8 responses) from psychopathology factor scores (AD, CI, SW; 3 predictors) for n=1954 participants. The primary model uses one component (**Figure 6A–B**); the second component of the two-component variant is not displayed — it generalizes only weakly out of sample (panel **(A)** and Methods). The overall one-component regression was significant (in-sample *R*^2^ = 0.019, equal-weighted across the eight responses; 1000-permutation *p* = 0.001). **(A)** How many components: mean cross-validated mean squared error on standardized data (y-axis *Mean CV MSE*) against the number of PLS components (x-axis *Number of PLS components*, ticks 0–3), with a blue dashed vertical line at the elbow point (elbow criterion: one dominant component; Methods). The curve shown is the 10-fold robustness variant of the canonical 5-fold × 10-shuffle crossvalidation, which returns the same verdicts: MSE falls from 1.0066 with no components to 0.9897 at one component, is minimized at 0.9890 at two and rises to 0.9895 at three, with SEs of about 0.0003 (smaller than the markers). The minimum at two components is marginal and the criteria disagree on it: a one-standard-error rule prefers two components and a per-component permutation test detects the second (*p* = 0.040 in this 10-fold variant; *p* = 0.010 in the canonical 5-fold selection), while the elbow criterion favors one dominant component and the second component generalizes only weakly to held-out participants — the basis of the one-component choice. The two-group stats block beside the curve reports the canonical 5-fold × 10-shuffle model statistics, not values read off this 10-fold curve: for Component 1 (the primary model), the in-sample latent-score correlation (*r* = 0.289), its held-out counterpart (*r*_CV_ = 0.275, two-sided permutation *p <* 0.001; see **Figure 6A**) and the one-component cross-validated *R*^2^ (0.0156, equal-weighted across the eight responses — the prediction counterpart of the in-sample 0.019 above); for Component 2, its held-out latent-score correlation (*r*_CV_ = 0.070), its two-sided permutation *p* = 0.014 and the incremental cross-validated *R*^2^ it adds (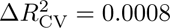; Methods). **(B)** How reproducible the weights are: boxplots of the pairwise absolute Pearson correlation between the weights recovered on different cross-validation splits (shared y scale, each cell carrying its own y label and tick labels; y-axes *Psych-weight* / *Econ-weight pairwise* |*r* | *across CV splits*, x-axis *Component*, ticks 1–3, on the lower cell). The upper cell, psychopathology-side weight stability, is the psychopathology (predictor) block and the lower cell, economic-side weight stability, the economic-preference (response) block. Psychopathology weights are near-perfectly reproducible at every component (median |*r*| = 0.9998, 0.979 and 0.9998 for components 1–3; with only three predictors the third component is fixed up to sign, which is why its box collapses back onto 1), whereas the economic weights decline monotonically (0.997, 0.969, 0.941) — consistent with a dominant, highly reproducible first component and progressively weaker later ones.

**Supplementary Figure 17.**
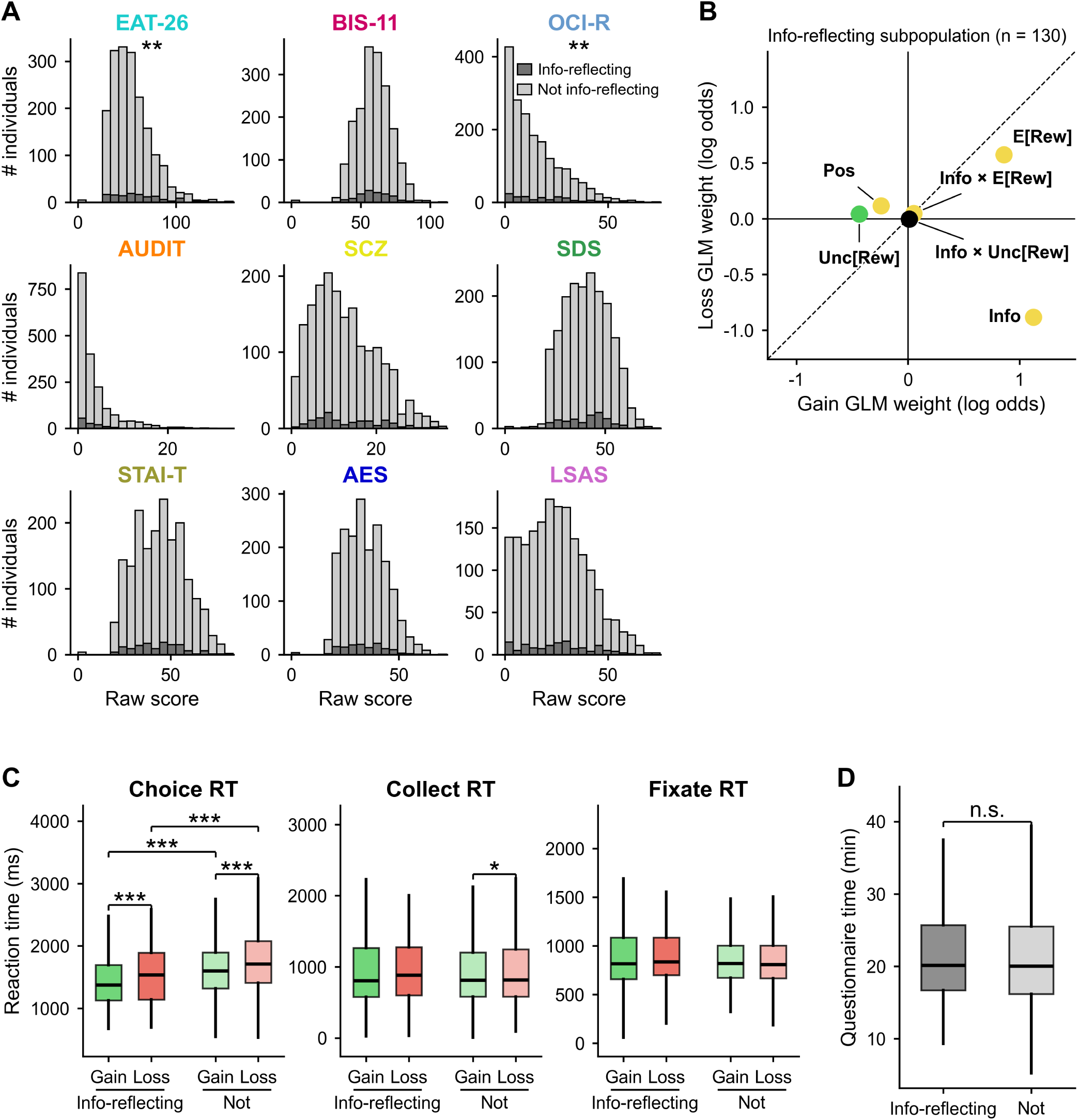
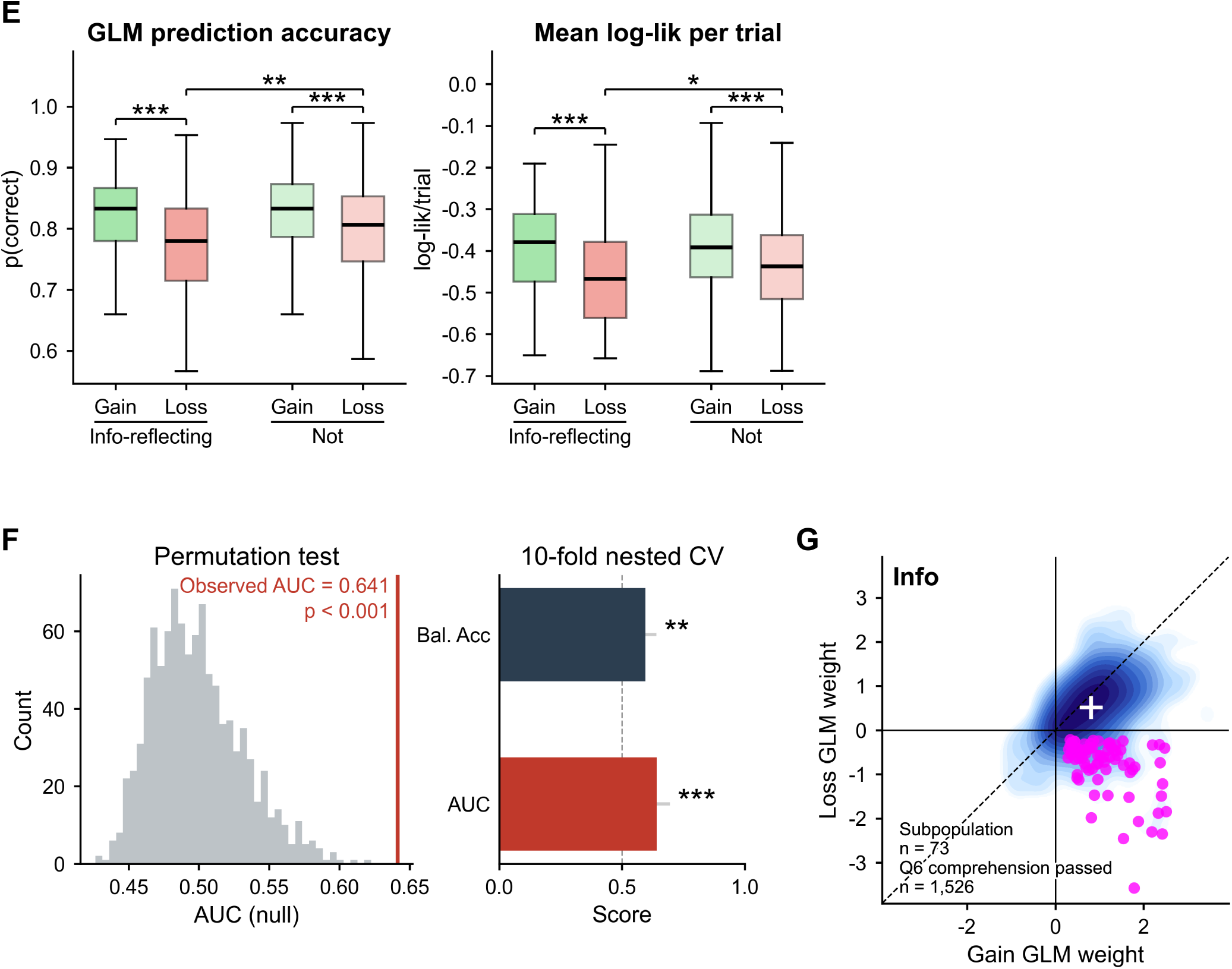
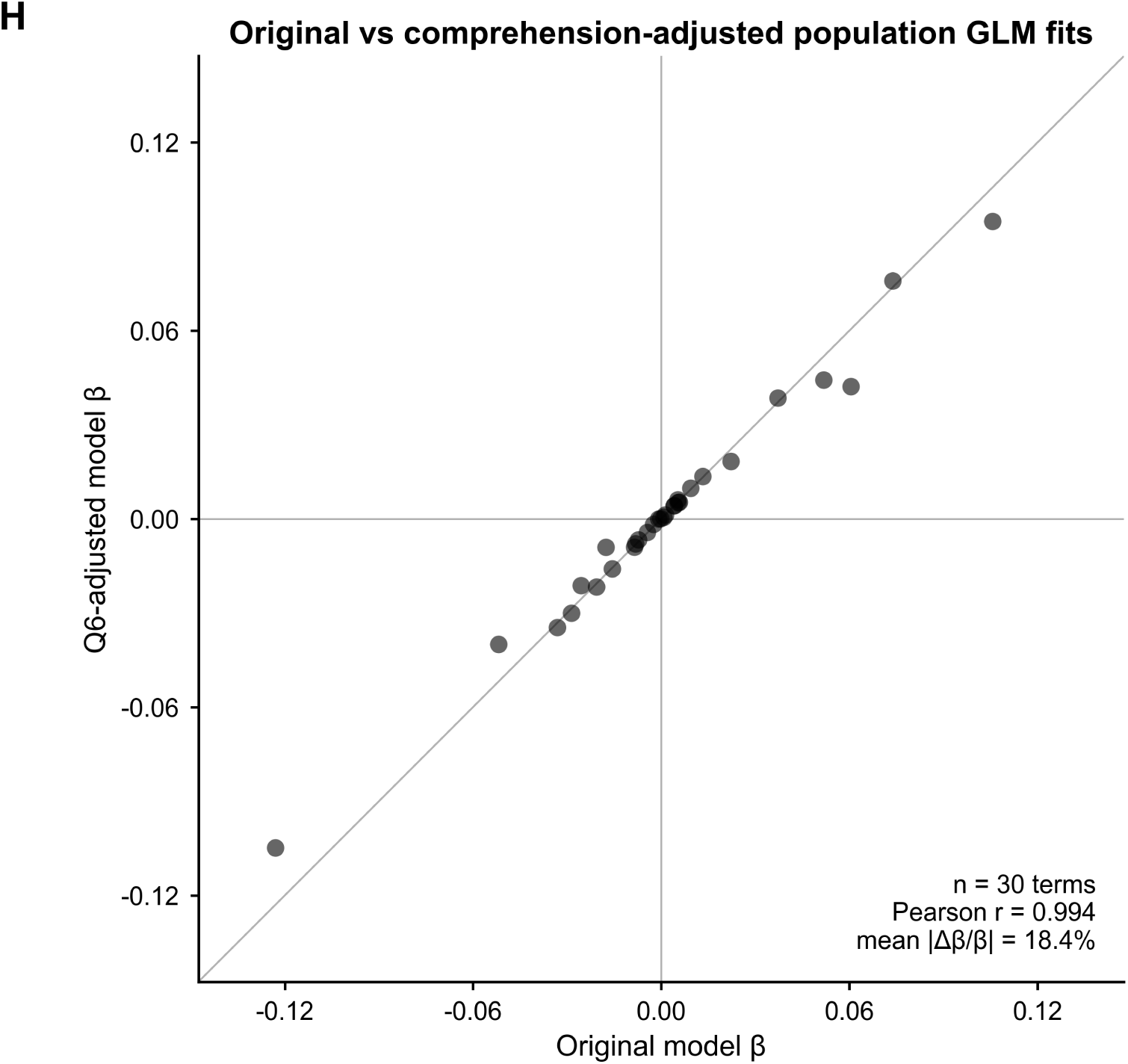
Symptom, response-time, fit-quality, classifier and comprehension characterization of the information-reflecting subgroup, and task-comprehension-adjusted robustness of the psychopathology results. Further characterization of the n=130 info-reflecting subpopulation — the individuals whose own choice-GLM Info weight was significantly positive in gain and significantly negative in loss (**Figure 6C**) — against the remaining n=1824 participants. The figure runs over three pages under one caption: **(A)** and **(B)** across the top of the first page with **(C)** and **(D)** beneath them; **(E)** across the top of the second with **(F)** and **(G)** beneath it; and **(H)** alone on the third. Panels **(A)**–**(G)** share the frozen subgroup definition (n=130 against n=1,824); **(H)** is a post-hoc, exploratory whole-sample check on the psychopathology results (Methods). The group comparisons in **(A)**, **(C)**, **(D)** and **(E)** are rank-based and reported raw and uncorrected (Methods); *, **, *** indicate *p <* 0.05, 0.01, 0.001 and *n.s.* is printed where a drawn comparison does not clear 0.05. **(A)** Raw questionnaire scores: a 3 × 3 grid of overlaid histograms, one per instrument (EAT-26, BIS-11, OCI-R, AUDIT, SCZ, SDS, STAI-T, AES, LSAS), each tile titled in that instrument’s **Figure 1B** color. Light gray is the non-info-reflecting subpopulation distribution (n=1824) and dark gray the info-reflecting subpopulation (n=130), drawn on shared bin edges with a per-tile count axis, so the subgroup’s bars are roughly fourteen times shorter throughout; a two-entry legend sits inside the OCI-R tile. Annotations are two-tailed rank-sum tests of the two groups’ scores, drawn where significant (unannotated tiles did not clear 0.05). Two instruments separate the groups, and both are CI-loaded: EAT-26 (median 58 vs. 51, *z* = 2.76, *p* = 5.8 × 10^-3^) and OCI-R (median 17 vs. 11, *z* = 3.02, *p* = 2.5 × 10^-3^), each marked **. The other seven are not significant (nearest SCZ, *p* = 0.13), including every AD-loaded instrument (SDS *p* = 0.88, STAI-T *p* = 0.45, AES *p* = 0.65) — the item-level counterpart of the selectively elevated CI factor score in **Figure 6E**. **(B)** The subgroup’s own average preferences: the mean across the n=130 members of each of the six individual choice-GLM weights, in gain (x-axis *Gain GLM weight (log odds)*) vs. loss (y-axis *Loss GLM weight (log odds)*), with SE bars smaller than the markers, a dashed unity line and solid zero rules; conventions as in **Figure 2D**, and loss-context E[Rew] and Info × E[Rew] are sign-flipped as elsewhere. Point color is a one-sample *t*-test against zero, significant in gain (green), loss (red), both (yellow) or neither (black), BH–FDR corrected over this panel’s 12 tests (6 regressors × 2 contexts). Info is the only economic-attribute weight whose sign reverses (gain +1.12 ± 0.06, loss −0.88 ± 0.06; both *p*_BH_ *<* 0.0001), which is the defining property of the group. On every other regressor the subgroup looks like a weaker version of the population rather than its mirror image: E[Rew] stays positive in both contexts (+0.86 ± 0.07 and +0.57 ± 0.06), the side bias Pos reverses in the population’s direction (−0.24 ± 0.04 and +0.12 ± 0.04), and Info × E[Rew] is positive in both (+0.052 ± 0.018, *p*_BH_ = 6.3 × 10^-3^; +0.046 ± 0.021, *p*_BH_ = 0.039). Unc[Rew] is green — negative in gain (−0.44 ± 0.08, *p*_BH_ *<* 0.0001) but indistinguishable from zero in loss (+0.04 ± 0.06, *p*_BH_ = 0.57) — and Info × Unc[Rew] is black in both. **(C)** Reaction times: three tiles, *Choice RT*, *Collect RT* and *Fixate RT*, with a single y-axis label (*Reaction time (ms)*) and per-tile scales and tick labels, each holding four boxes — Gain (green) and Loss (red) for the info-reflecting subpopulation (saturated) and for *Not* (light). Within-group gain-vs-loss brackets are Wilcoxon signed-rank tests, between-group brackets are Mann–Whitney U tests, and group × context interactions are Mann–Whitney tests on the within-participant gain-minus-loss differences. Choice RT is the only event on which the groups differ, and the subgroup is faster, not slower: median 1373 vs. 1599 ms in gain (*p <* 0.0001) and 1536 vs. 1712 ms in loss (*p <* 0.0001), with both groups slowing by a comparable amount from gain to loss (both within-group *p <* 0.0001; interaction *p* = 0.94). Collect RT differs between contexts only in the non-info-reflecting subpopulation (*p* = 0.041) and never between groups (*p* ≥ 0.30; interaction *p* = 0.24), and no Fixate RT comparison is significant (nearest *p* = 0.069; interaction *p* = 0.26). The subgroup is therefore not characterized by generalized slowing, and its gain-to-loss change in speed is the same as everyone else’s. **(D)** Total questionnaire completion time (y-axis *Questionnaire time (min)*) for the info-reflecting subpopulation (dark gray) and *Not* (light gray), with a Mann–Whitney bracket. The two do not differ (median 20.2 vs. 20.0 min; mean 22.1 vs. 21.9; *p* = 0.95), so subpopulation membership is not a by-product of hurried questionnaire responding. **(E)** Quality of the individual choice-GLM fits, in two tiles: *GLM prediction accuracy* (y-axis *p(correct)*) and *Mean log-lik per trial* (y-axis *log-lik/trial*). Each tile holds four boxes in the drawn order Gain (info-reflecting subpopulation), Loss (info-reflecting subpopulation), Gain (other), Loss (other), with the group names on a second label line and the color convention of **(C)**; tests are as in **(C)**. Both groups are fit less well in loss than in gain (all four within-group comparisons *p <* 0.0001). In the gain context the subgroup is indistinguishable from the rest (median accuracy 0.833 vs. 0.833, *p* = 0.69; median log-likelihood −0.379 vs. −0.391, *p* = 0.80). In the loss context it is modestly worse (0.780 vs. 0.807, *p* = 9.2 × 10^-3^; −0.467 vs. −0.437, *p* = 0.024) — a gap of under three percentage points of correctly predicted choices — and the group × context interaction is not significant for either metric (*p* = 0.063 and *p* = 0.087). Info-reflecting subpopulation members are therefore not a pool of noisy or poorly fit participants. **(F)** Diagnostics for the L1-regularized logistic classifier of **Figure 6F–G**. Left: the null distribution of held-out AUC over 1000 model-level label permutations, each with a full nested cross-validation refit (gray histogram; x-axis *AUC (nul l)*, y-axis *Count*), with a red vertical rule at the observed value labeled inline (*Observed AUC = 0.641*) with the permutation p-value printed beneath it (*p <* 0.001; the null has mean 0.499 and SD 0.032, and its maximum over 1000 permutations is 0.623, i.e. below the observed AUC). Right: nested cross-validated performance (10-fold outer × 5-fold inner) as horizontal bars against a gray dashed chance line at 0.5 (x-axis *Score*) — balanced accuracy 0.596 (**) and ROC AUC 0.641 (***) — each starred by its own nested-CV label-permutation test. Sensitivity and specificity are deliberately not drawn here: each is a single threshold-dependent operating point that does not exceed its own permutation null, and they are reported, with that caveat, in the **Figure 6G** caption. Because ROC AUC does not depend on class prevalence, we note alongside it the prevalence-sensitive summary of the same held-out predictions: area under the precision–recall curve 0.128 against a chance baseline of 0.067 (the positive-class base rate), and positive predictive value 9.9% at the 0.5 threshold (Methods). **(G)** Comprehension-filtered version of the **Figure 6C** density scatter, restricted to the N = 1526 of 1954 participants who answered the Q6 task-comprehension question correctly in both contexts. Axes are *Gain GLM weight* and *Loss GLM weight*, and the regressor name (*Info*) is annotated in bold at the upper left rather than carried in the axis titles. The filled blue log-density contour is the comprehension-passing sample, magenta points are the info-reflecting subpopulation members who pass the filter, the white crosshair is that sample’s mean ± SEM (+0.802 ± 0.015 in gain, +0.516 ± 0.016 in loss), the dashed line is unity and the solid lines mark zero; the gray outline cloud of non-members drawn in earlier versions is deliberately omitted, since the magenta points are already a visible subset. 73 of the 130 subgroup members (56%) pass the comprehension filter against a 78% retention rate in the sample as a whole, and those 73 occupy the same lower-right region of the plane as the full set: the information-reflecting pattern is attenuated but not created by comprehension failures, and it is plainly present among participants who demonstrably understood the task. **(H)** Post-hoc, exploratory: original versus Q6-adjusted population GLM coefficients. The companion to **Supplementary Figure 7B**, drawn by the same routine on the same 30 adjusted terms — each of the five economic regressors (E[Rew], Unc[Rew], Info, Info × E[Rew], Info × Unc[Rew]) crossed with each of the three psychopathology factors (AD, CI, SW), entered both as a factor interaction and as a context × factor interaction. The x-axis (*Original model β*) is the published population (cluster-robust) choice GLM; the y-axis (*Q6-adjusted model β*) is that same model refit on the full n=1,954 with a *z*-scored Q6 comprehension-pass indicator added to the moderator block — the “Q6 adjustment” (Methods), whose restriction-based counterpart defines panel **(G)**, so the two ways of using Q6 sit in one figure. Gray rules mark zero on each axis, the diagonal is the identity line, and the annotation reports the number of terms, their Pearson correlation and the mean absolute fractional change in coefficient. Adjusting for comprehension leaves the coefficient pattern essentially intact: the 30 terms correlate at *r* = 0.994 (*p* = 2.3 × 10^-28^), 29 of the 30 terms keep their sign (the single exception is the smallest term, |*β*| ≈ 10^-4^), and no term’s rank relative to the others is materially altered. What the adjustment does is shrink the estimates, close to uniformly: the best-fitting slope through the origin is 0.882, so adjusted coefficients are on average about 12% smaller than their published counterparts. The mean absolute fractional change (18.4%) exceeds the 12% implied by that slope because a fractional change is dominated by the smallest coefficients, where the same absolute shift is a larger share of the estimate.

**Table 1: Supplementary Table 1.**
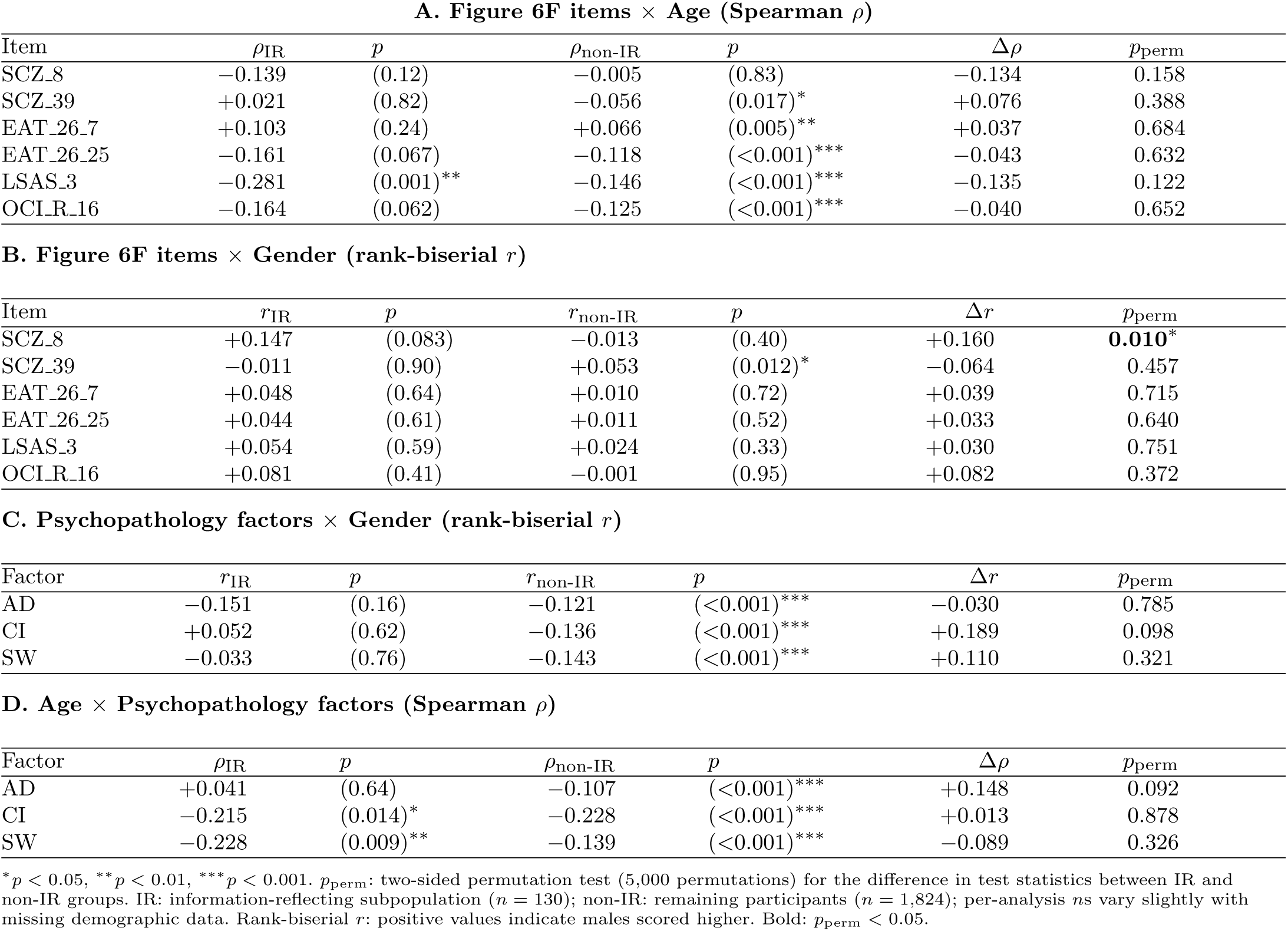
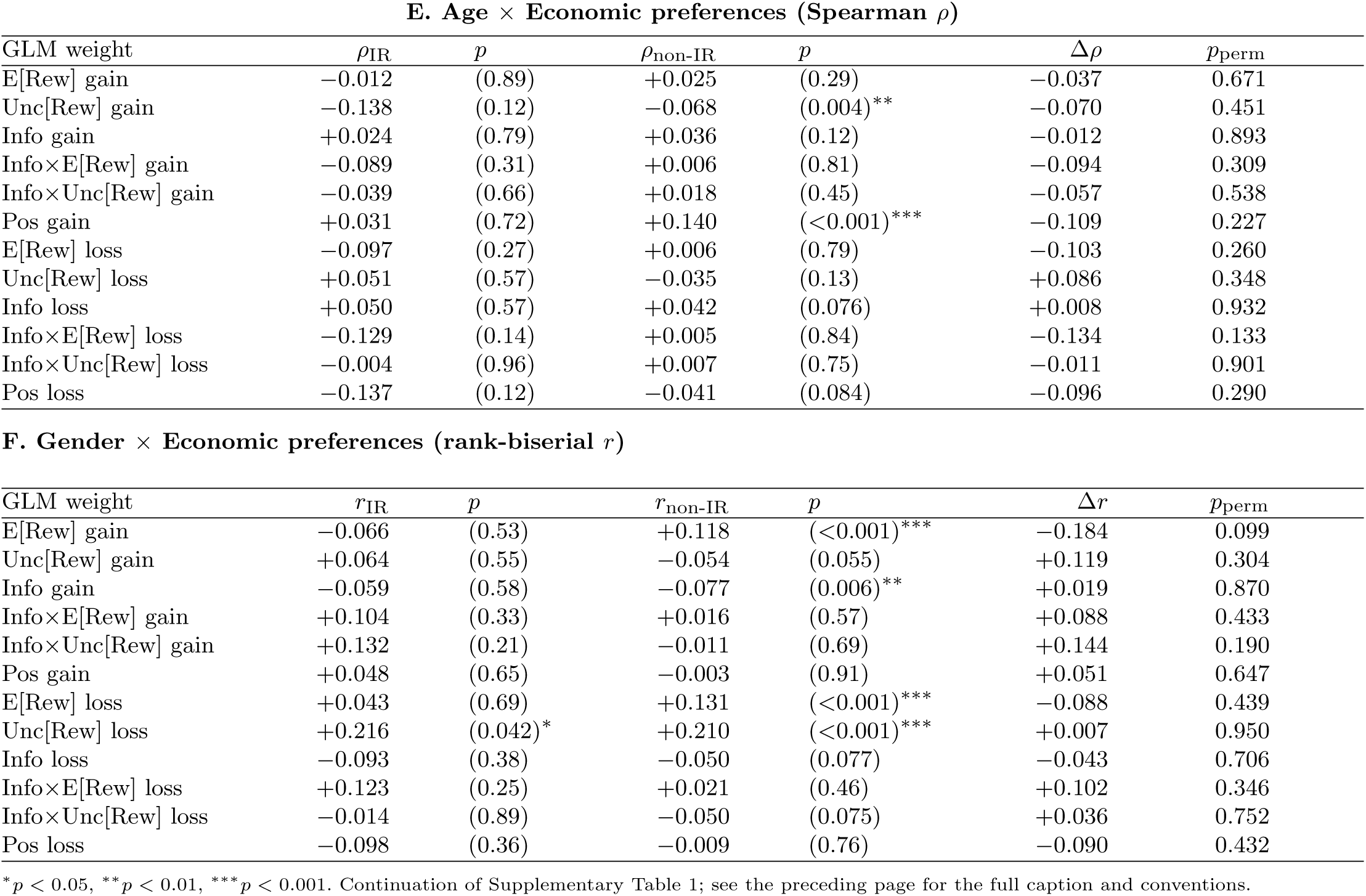
Demographic relationships are not moderated by information-reflecting subpopulation membership. Every analysis is run twice—once within the information-reflecting subpopulation (IR; *n* = 130) and once within the remaining participants (non-IR; *n* = 1,824)—and the two resulting statistics are then compared by a label-permutation test (5,000 permutations; *p*_perm_). Per-analysis *n*s vary slightly with missing demographic data. **(A)** Age effects on the six sparse-classifier questionnaire items of **Figure 6F**, as Spearman *ρ* within each group; the permutation test evaluates *H*_0_: *ρ*_IR_ = *ρ*_non-IR_. **(B)** Gender effects on the same six items, as Mann–Whitney rank-biserial *r* (positive = males scored higher, as in the footnote); the permutation test evaluates *H*_0_: *r*_IR_ = *r*_non-IR_. **(C)** Gender effects on the psychopathology factor scores (AD, CI, SW), same statistic and test as **(B)**. **(D)** Age–psychopathology correlations, same statistic and test as **(A)**. **(E)** Age–economic preference correlations, over the six individual GLM weights in each context. **(F)** Gender–economic preference effects, over the same twelve weights. Across all 42 comparisons only one permutation test reached significance (SCZ 8 × gender, Panel B; *p*_perm_ = 0.010). Two within-group patterns are nonetheless worth noting: the gender differences in psychopathology factor scores that are robust among the non-IR participants (all *p <* 0.001) are absent within the IR subgroup (all *p >* 0.15; Panel C), and LSAS 3 shows a stronger negative age correlation inside the subpopulation (*ρ*_IR_ = −0.281, *p* = 0.001) than outside it (*ρ*_non-IR_ = −0.146; *p*_perm_ = 0.122; Panel A). Overall, demographic relationships with questionnaire items, psychopathology factors, and economic preferences are not strongly moderated by information-reflecting subpopulation membership, though these within-subpopulation patterns are potentially consistent with a partially distinct demographic–symptom structure.

